# Long-term reliability and stability of parameterized resting state EEG: Evidence from a five-year follow-up

**DOI:** 10.64898/2026.03.03.709208

**Authors:** Polina Politanskaia, Jacinta Bywater, Anna J. Finley, Hannah A.D Keage, Nicholas J. Kelley, Daniel McKeown, Victor Schinazi, Douglas J Angus

## Abstract

Several aspects of parameterized neural activity, including the aperiodic exponent and individual peak alpha frequency, have emerged as promising biomarkers for ageing, pathology, and cognitive decline. Their potential clinical application is tempered by a lack of evidence on long-term temporal stability. Existing investigations have largely relied on cross-sectional designs or considered stability for up to 90 days. Here, we examined five-year reliability, stability, and age-related changes in periodic and aperiodic neural activity using electroencephalography in adults aged 20-70 years. Resting-state EEG was recorded in two sessions, approximately five years apart. We extracted the aperiodic exponent, aperiodic offset, peak alpha power, and individual alpha peak frequency and examined test-retest reliability at both the channel and cluster levels. All parameters demonstrated fair to excellent test-retest reliability (intraclass correlations = 0.51-0.88). Linear mixed models revealed that individual peak alpha frequency decreased, the aperiodic exponent flattened, and parameterized alpha power remained unchanged. There were no interactions between time and age. Our findings suggest that parameterized activity is reliable over long timeframes, and demonstrates changes consistent with ageing-related processes. Spectral parameterization may provide a means of characterizing within-person neurophysiological changes across adulthood. Future research should explore the utility of identifying deviations that may indicate pathology.

## Introduction

Understanding changes in neural activity across the lifespan is necessary for distinguishing between healthy and pathological aging. Since its development over 100 years ago, electroencephalography (EEG) has emerged as a critical tool for neuroscientists and clinicians hoping to understand how, why, and in whom neural activity changes. Traditionally, EEG research has focused on oscillatory (periodic) activity within canonical bands. Notably, variation in the amplitude and peak frequencies of alpha (8-13 Hz) oscillations has been linked to attention, memory, and executive control (Buzsáki & Watson, 2012; Immink et al., 2021; Klimesch, 1997). However, emerging evidence highlights the importance of non-oscillatory (aperiodic) activity as a distinct and complementary index of neural functioning. Despite recent advances in spectral parameterization techniques (Donoghue et al., 2020) and the understanding of the physiological significance of aperiodic activity (Gao et al., 2017), the temporal reliability and stability of these measures remain poorly characterized. One reason for this is because almost all studies use cross-sectional data or have examined reliability and change over short timeframes, reducing their utility as reliable markers of neural ageing or indicators of pathological deviation. To address this gap, we investigate within-person changes in both aperiodic and periodic components of the EEG power spectrum across a five-year period. We show that parametrized aperiodic and periodic activity are reliable over time through rank order stability across a five-year timespan. Additionally, we demonstrate that these measures change systematically as a function of time.

### Characterizing Aperiodic and Periodic Neural Activity

The majority of the neural power spectrum is comprised of non-oscillatory (i.e., aperiodic) activity, which follows a 1/f^χ-like distribution, such that as frequency increases, power decreases. The most used method for parameterizing aperiodic activity is SpecParam (formerly FOOOF; Donoghue et al., 2020), which characterizes aperiodic activity into two parameters: the aperiodic exponent (χ) and the offset. The exponent reflects the rate at which power decreases across frequencies, while the offset indicates the overall magnitude of neural activity across the frequency spectrum. Computational and pharmacological studies suggest that the aperiodic exponent is a non-invasive index of circuit-level excitation:inhibition (E:I) balance (Gao et al., 2017; McKeon et al., 2023; Waschke et al., 2021), with greater exponents (i.e., steeper slopes) associated with greater cortical inhibition. Conversely, lower exponents (i.e., flatter slopes) are linked to increased excitability, reduced inhibition, and greater cortical noise (Voytek et al., 2015). Alterations in the offset parameter reflect changes in neuronal firing rates and arousal states (Manning et al., 2009).

These aperiodic parameters show systematic variation in healthy aging. Cross-sectional studies report that the aperiodic exponent increases with age from infancy through early adolescence (Hill et al., 2022; Schaworonkow & Voytek, 2021), before gradually decreasing from early adulthood to older age (Finley et al., 2022; Merkin et al., 2022). Beyond age-related alterations, systematic variation in the aperiodic exponent has been linked to systematic variation in cognitive functioning. Individuals with lower (i.e., flatter) exponents tend to perform worse on cognitive tasks that assess response time, working memory capacity, and attentional control (Euler et al., 2024; Pathania et al., 2022; Voytek et al., 2015). Moreover, consistent with the E:I balance model, the exponent is altered in multiple developmental and neurological disorders associated with altered excitatory and inhibitory neurotransmission, including Parkinson’s disease (Agouram et al., 2025; Helson et al., 2023; McKeown et al., 2023; Monchy et al., 2024), ADHD (Ostlund et al., 2021), and autism spectrum disorder (Shuffrey et al., 2022). Similar findings have been observed for the aperiodic offset, with lower offsets associated with poorer verbal fluency (McKeown et al., 2025), and reduced cognitive control (Clements et al., 2021).

Alpha oscillations (8-13 Hz) also show age-related changes with slowing of the individual alpha peak frequency (IAPF) linked to cognitive decline in older adults, and higher IAPF predicting better performance across multiple domains (Angelakis et al., 2004; Grandy, Werkle-Bergner, Chicherio, Lövdén, et al., 2013; Grandy, Werkle-Bergner, Chicherio, Schmiedek, et al., 2013). Power within the canonical alpha band (i.e., the unparameterised power between 8 −13 Hz) also decreases with age, and lower power is associated with reduced attentional capacity and executive control (Clements et al., 2021). These findings align with the broader literature, which consistently shows that fluctuations in alpha power are associated with attentional engagement and task performance (Klimesch, 1997; Klimesch et al., 2007). Recent work has also demonstrated that aperiodic and periodic activity are jointly important in predicting cognitive function, with one study finding that the aperiodic exponent and IAPF interact to predict cognitive decline over a 10-year period (Finley et al., 2024). In the current study, we focus on alpha power and IAPF due to their established sensitivity to age-related cognitive changes and their potential as biomarkers, and the consistency with which oscillatory ‘peaks’ within the alpha band can be detected using parameterization tools.

### Reliability and Stability of Aperiodic Activity and Alpha Oscillations

For aperiodic and periodic EEG features to serve as reliable markers of neural ageing or indicators of pathological deviation, we must establish both their test-retest reliability (i.e., the rank-order of individual differences over time) and their trajectories of mean-level change with age (i.e., their stability). Moreover, although cross-sectional data show that aperiodic activity is correlated with age (Finley et al., 2022; Smith et al., 2022; Tran et al., 2020; Voytek et al., 2015), it remains unknown if the magnitude of change in aperiodic measures varies with age.

Previous studies examining the test-retest reliability of parameterized aperiodic and periodic activity have yielded mixed results. In young adults, the intra-class correlations (ICC) for the aperiodic exponent are good to excellent over a 30-minute timeframe (ICC’s .78–.93; Pathania et al., 2021), and good over measures taken 90 minutes apart and 30 days later (average ICC’s .64-73; McKeown et al., 2024). In school-aged children (i.e., age 4–11 years), test-retest reliability across approximately six days is highly variable, with good reliability for the aperiodic exponent (ICC = .70), and fair reliability for the aperiodic offset (ICC = .53) in autistic children, but poor to fair reliability in typically developing children (aperiodic exponent ICC = .28, aperiodic offset ICC = .48; Levin et al., 2020).

The few studies examining test-retest reliability in older adults have focused on periodic activity, finding that both parameterized alpha power and IAPF have excellent reliability across approximately seven days in both younger (20-35 years old; ICC’s .78- .83) and older (60-80 years old; ICC’s .75-.81) adults (Popov et al., 2023), and good to excellent reliability in autistic and typically developing children (ICC’s .62-.86; Levin et al., 2020). Over longer timeframes, alpha power has extremely good test-retest reliability in younger adults, with Pearson’s *r* correlations between .72 and .85 observed for measures taken 12 years apart (Tenke et al., 2018).

Given the links to health and pathological aging, and potential as a biomarker, understanding the longer-term reliability of aperiodic activity, individual alpha peak frequency, and parameterized alpha power is necessary to establish how reliably these measures track individuals over the extended timeframes across which cognitive decline and pathology-related changes may be expected to occur, and to examine if and how these metrics change over time.

### The Current Study

In this study, we investigated the test-retest reliability, stability, and age-related changes in both aperiodic (exponent and offset), and periodic (IAPF and alpha power) components of the EEG power spectrum over a five-year period in a sample of normally aging adults. We use longitudinal data from the Dortmund Vital Study (Gajewski et al., 2022; Getzmann et al., 2024) to systematically examine the reliability and stability of these metrics in adults aged 20 to 70 years. Although we did not preregister our hypotheses, our work was guided by the following empirically supported predictions. First, we anticipated that the test-retest reliability of the aperiodic exponent, aperiodic offset, IAPF, and alpha power would be at least fair (e.g., ICC >.40; Cicchetti, 1994). Second, we predicted that the aperiodic exponent, aperiodic offset, IAPF, and alpha power would be negatively associated with age, such that older adults would have lower exponents, lower offsets, slower IAPF, and lower alpha. Third, the aperiodic and alpha measures were predicted to decrease over the five-year period. Finally, we predicted that the magnitude of this decrease would be moderated by age, with older adults showing a greater decrease (Finkel et al., 2003; Pfefferbaum & Sullivan, 2015; Zaninotto et al., 2018). In addition, we explored whether these trajectories differ by sex, though we made no specific directional predictions.

## Method

### Data Availability

Detailed descriptions of the original data collection and processing procedures can be found in Getzmann et al. (2024). All EEG recordings are publicly available in BIDS format available at https://doi.org/10.18112/openneuro.ds005385.v1.0.2). Jupyter and *R*-markdown notebooks and scripts for reproducing our results are available at https://github.com/MindSpaceLab/Aperiodic_Test_Retest_5year

### Participants

The sample was assessed at two sessions. Commencing in 2021, Session 1 data were collected from 608 participants aged 20 to 70 years (*M* = 44.10, *SD* = 14.50), including 376 females (61.8%) and 232 males (38.2%). Most participants were right-handed (*n* = 566, 93.1%), with 40 left-handed (6.6%) and two participants with missing handedness information. For Session 2, approximately five years later, 208 participants (34% of those in Session 1) returned for a follow-up assessment. The follow-up sample included 130 females (62.5%) and 78 males (37.5%), with 201 right-handed (96.6%) and 7 left-handed (3.4%) participants. Participants with both session 1 and session 2 data (*M* = 47.05, *SD* = 13.75) were older than those with only session 1 data (*M* = 42.54, *SD* = 14.68, *t*(441.4) = 3.74, *p* <.001), and had greater peak alpha power over occipitoparietal sensors in the eyes-closed condition (both sessions *M* = 1.22, *SD* = 0.45, *t*(325) = 2.16, *p* = .031). There were no other significant differences in demographic or EEG metrics (see supplementary info).

Participants were recruited from the Dortmund area (Germany) through local companies, public institutions, and media advertisements. Exclusion criteria were severe neurological (e.g., dementia, Parkinson’s disease, stroke), cardiovascular, oncological, eye diseases, psychiatric disorders, significant head injury/surgery/implants, psychotropic/neuroleptic medication use, and limited physical fitness or mobility. Participants taking blood thinners, hormones, antihypertensives, or cholesterol-reducers were permitted. Written informed consent was obtained from all participants before study commencement. The research protocol adhered to the World Medical Association’s Code of Ethics (Declaration of Helsinki) and received approval from the local Ethical Committee of the Leibniz Research Centre for Working Environment and Human Factors in Dortmund, Germany (ClinicalTrials.gov Identifier: NCT05155397, approval numbers: A93-1 and A93-3 for the follow-up testing).

Participants with poor EEG data quality (excessive artifacts or technical failures) were excluded by the original authors (Getzmann et al., 2024), although the exact number of participants excluded was unspecified. We excluded participants if any of the following criteria were met: either reference channel (TP9/TP10) was identified as bad, insufficient artifact-free data (<50% artifact-free epochs per condition), poor SpecParam model fits (r^2^ < .80) on > 50% of channels, and an absence of alpha peaks on >50% of channels. Criteria relating to reference channels and artifactual data were applied in a list-wise fashion, while criteria relating to model fit and peak detection were applied on an analysis-wise basis^1^. Demographic characteristics for the retained sample for each analysis are presented in Table 1.

**Table 1.** Demographic and SpecParam Model Characteristics for Analysis of Aperiodic and Alpha Activity.

|  | Aperiodic |  | Alpha |  |
| --- | --- | --- | --- | --- |
|  | Eyes-Closed | Eyes-Open | Eyes-Closed | Eyes-Open |
|  | *n* = 178 | *n* = 161 | *n* = 175 | *n* = 147 |
| Age | 46.92 (13.83) | 46.67 (13.68) | 46.83 (13.95) | 46.78 (13.88) |
|  | [20 – 70] | [22 – 70] | [20 – 70] | [22 – 70] |
| Sex | 116 F / 62 M | 106 F / 55 M | 114 F / 61 M | 97 F / 50 M |
| Epochs Session 1 | 87.20 (8.44) | 86.84 (7.35) | 87.09 (8.44) | 86.95 (7.15) |
|  | [24 – 107] | [47 – 110] | [24 – 107] | [47 – 110] |
| Epochs % Session 1 | 93.88 (6.46) | 93.40 (7.26) | 93.01 (6.49) | 93.53 (7.09) |
|  | [56.38 – 100] | [51.65 – 100] | [56.38 – 100] | [51.65 – 100] |
| Epochs Session 2 | 87.88 (5.46) | 86.58 (5.97) | 87.86 (5.49) | 86.57 (6.21) |
|  | [67 – 116] | [49 – 108] | [67 – 116] | [49 – 108] |
| Epochs % Session 2 | 95.25 (3.61) | 94.11 (5.48) | 95.23 (3.64) | 94.05 (5.66) |
|  | [72.83 – 100] | [53.85 – 100] | [72.83 – 100] | [53.85 – 100] |
| Mean $r^2$ Session 1 | .98 (.01) | .98 (.01) | .98 (.01) | .98 (.01) |
|  | [.94 – .99] | [.94 – .99] | [.94 – .99] | [.94 – .99] |
| Mean $r^2$ Session 2 | .98 (.01) | .98 (.01) | .98 (.01) | .98 (.01) |
|  | [.92 – .99] | [.93 – .99] | [.92 – .99] | [.93 – .99] |
| MAE Session 1 | .05 (.01) | .04 (.01) | .05 (.01) | .04 (.01) |
|  | [.03 – .08] | [.03 – .06] | [.03 – .08] | [.03 – .06] |

<sup>1</sup> We opted for this approach rather than exclusion at the list-wise level. Many participants did not have clear alpha peaks in the eyes-open condition, and a list-wise approach would have reduced the analysable sample for all analyses, rather than just those involving individual peak alpha frequency, or alpha amplitude.

|  |  |  |  |  |
| --- | --- | --- | --- | --- |
| MAE Session 2 | .05 (.01) | .04 (.01) | .05 (.01) | .04 (.01) |
|  | [.03 – .09] | [.03 – .07] | [.03 – .09] | [.03 – .07] |

Note. Each analysis consists of partially overlapping subsets of the same sample. Standard deviations are presented in parentheses, ranges are presented in brackets. MAE = Mean Absolute Error

### Materials

Demographic information, including age, sex, and handedness, was collected during the intake process (Getzmann et al., 2024). Only pre-task resting-state EEG recordings were analysed in the present study. Post-task recordings were excluded to avoid potential confounding effects of cognitive exertion and fatigue. Resting-state EEG was recorded in two conditions: eyes closed (EC) and eyes open (EO).

### Procedure

Participants were invited to the laboratory for a morning testing session. Sessions were scheduled at approximately the same time for all participants to promote a consistent state of wakefulness. Upon arrival, participants provided informed consent and completed health and demographic questionnaires. All testing took place in a quiet room designed to reduce environmental noise and electrical interference (Getzmann et al., 2024). After completing the questionnaires, participants were fitted with a 64-channel EEG cap by a trained researcher in preparation for the recording.

The testing session began with the pre-task resting-state EEG recording, consisting of three minutes of EC followed by three minutes of EO. For the EC condition, participants were instructed to sit quietly, remain still, and keep their eyes closed while maintaining a state of relaxed wakefulness. For the EO participants, they were asked to maintain their gaze on a fixation cross presented on a monitor positioned approximately 1 meter in front of them to minimize eye movements while keeping a similarly relaxed state. Approximately five years after the initial session, participants were invited back for a follow-up session (Session 2) that followed identical procedures to the first session. The full details of the cognitive tasks and experimental procedures are described extensively in Getzmann et al. (2024).

### Physiological Recording and Data Reduction

The resting-state EEG was recorded using a 64-channel elastic cap with electrodes connected using saline solution positioned according to the international 10–20 system, with FCz as the reference electrode during recording. Data were acquired with a BrainVision BrainAmp DC amplifier and the BrainVision Recorder software (Brain Products GmbH). EEG signals were digitised at 1000 Hz. Electrode impedances were kept below 10 kΩ during each resting session (Getzmann et al., 2024).

We processed the raw resting-state EEG data using MNE-Python (Gramfort et al., 2013). The data were first downsampled to 250 Hz, and bad channels were identified and removed using RANSAC (Session 1 Eyes Closed: *M* = 1.18, *SD* = 1.62, max = 9; Session 1 Eyes Open: *M* = 1.49, *SD* = 1.87, max = 11; Session 2 Eyes Closed: *M* = 0.94, *SD* = 1.58, max = 9; Session 2 Eyes Open: *M* = 1.59, *SD* = 2.00, max = 11). The remaining good channels were then re-referenced to the average of the TP9 and TP10 electrodes. After re-referencing, FCz was recreated as an active channel.

Then, we used Independent Components Analysis (ICA) reduce the dimensionality of the data to 30 independent components for further artifact correction. In keeping with published guidelines (Pion-Tonachini et al., 2019), ICA was applied to a 1–100 Hz bandpass-filtered copy of the data. Components associated with eye movements, muscle activity, and cardiac signals were automatically identified and removed from the unfiltered data (Pion-Tonachini et al., 2019). Bad channels were then replaced by interpolation. Resting-state EEG data was then band-pass filtered between 0.1 and 40 Hz, and segmented into 2000 ms, non-overlapping epochs. Epochs with peak-to-peak voltages exceeding 200 μV on any channel were rejected and excluded from further analysis.

Power Spectral Density was estimated using Fast Fourier Transform (Welch method, 2000ms Hamming window, 50% overlap). PSDs were then averaged for each channel. PSDs for each channel, participant, and condition were parameterised using the SpecParam package (v1.01; Donoghue et al., 2020). Each PSD was decomposed into periodic (oscillatory) and aperiodic (1/f) components within the 3–40 Hz range (peak width limits = 1–8 Hz, maximum peaks = infinite, minimum peak height = 0.1 µV², peak threshold = 2.0, aperiodic mode ‘fixed’). From each SpecParam model fit, we extracted the aperiodic exponent and aperiodic offset, the peak alpha frequency and peak alpha power in the 8–13 Hz range, and the model R² and mean absolute error (MAE).

### Statistical Analyses

To identify regions of interest for ICC and HLM analyses, we used Principal Components Analysis (PCA) followed by k-means clustering (Euler et al., 2024; McKeown et al., 2025). For each measure and condition combination (e.g, Exponent Eyes-Closed), channel-wise data were first submitted to a PCA. Visual inspection of scree-plots indicated that each measure and condition combination was best represented by a two-factor solution. To confirm the number of clusters and channel membership within clusters, we then used k-means clustering. We fit a range of k-means models, from two to eleven, and used the silhouette scores to determine the optimum number of clusters for each measure and condition, selecting the k-means model with the largest silhouette score (Table S1). The resulting clusters are presented in Figure 1^2^. We note that the identified clusters are diffuse, potentially due to the spatial smoothing introduced by the interpolation of missing channels. For cluster-level analysis, we calculated the mean measurement for each participant within each cluster for each measurement type, condition, and session.

**Figure 1.**
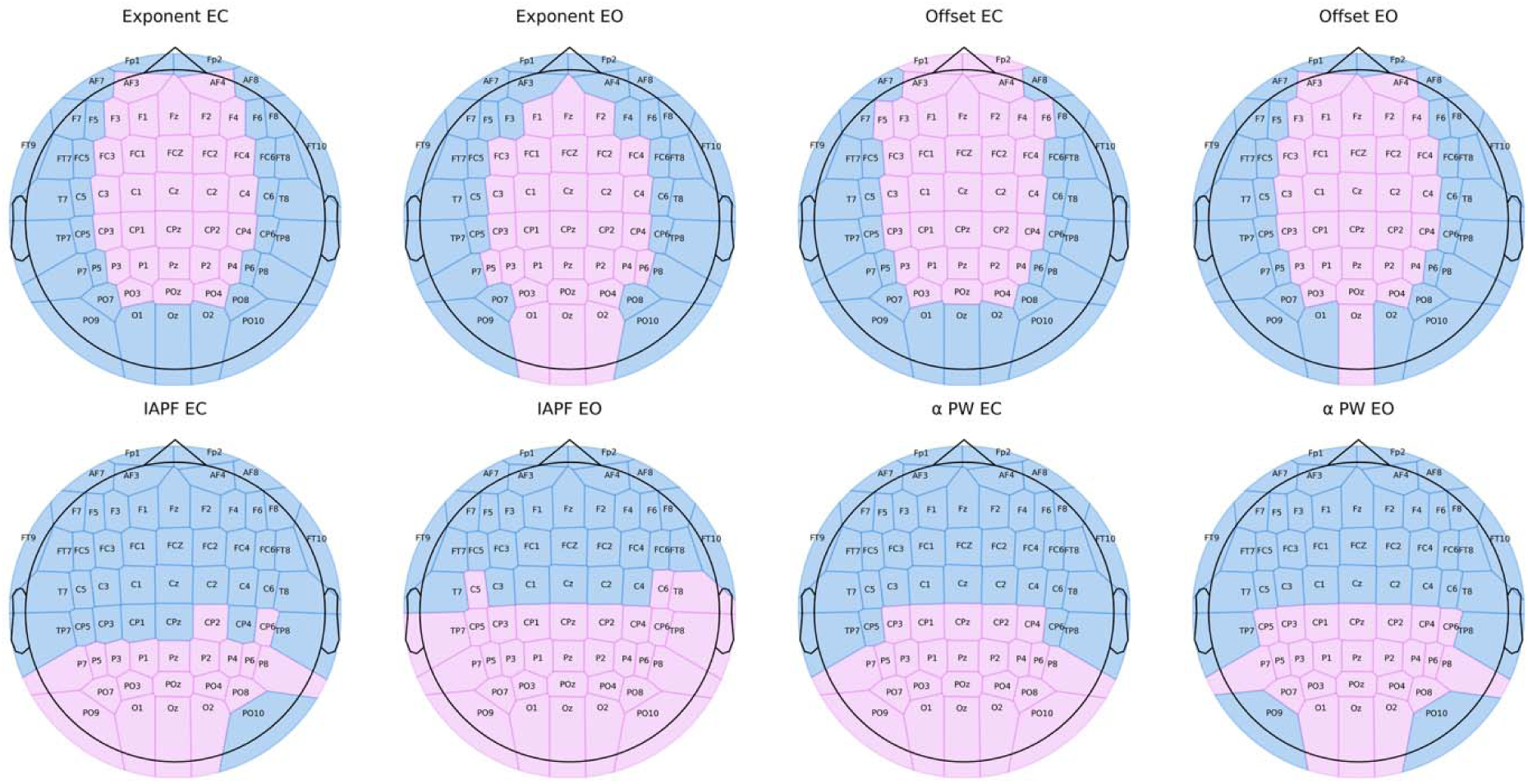
Topographic plots depicting k-means derived clusters for each measure and condition. Note: IAPF = individual alpha peak frequency, PW = power, EC = eyes-closed, EO = eyes-open.

**Figure 2.**
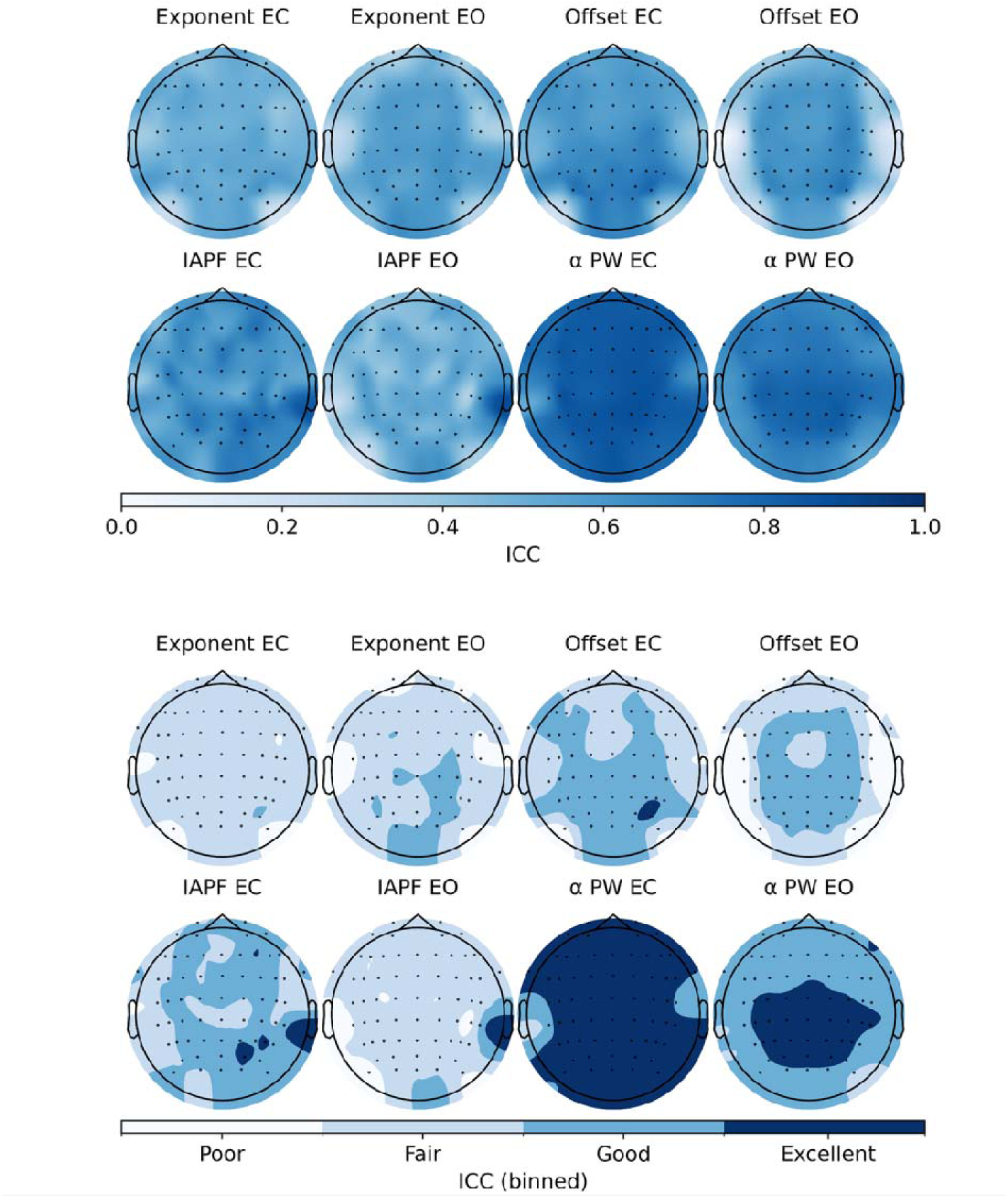
Topographic plots depicting Intraclass Correlations for the Aperiodic Exponent, Aperiodic Offset, Individual Alpha Peak Frequency, and Alpha Power. Note: The upper panel shows individual ICCs at the channel level. The lower panel shows ICCs at the channel level, after binning into thresholds (poor = 0 – .39, Fair = .40 – .59, Good = .60 – .74, Excellent = .75 – 1.00). IAPF = individual alpha peak frequency, PW = power, EC = eyes-closed, EO = eyes-open.

To examine test-retest reliability, we first calculated intra-class correlations at the individual channel level and then at the cluster level for each measure and condition (McKeown et al., 2024), using *pyirr* (de Klerk, 2022; Gamer et al., 2005) in Python. In keeping with our aim to evaluate the potential of parameterised activity as a reliable biomarker, we calculated ICCs using a two-way random-effects model (ICC[2,1]). We used the following thresholds for interpreting ICC values: <.40, poor; .40-.59, fair; .60- .74, good; .75-1.00, excellent (Cicchetti, 1994; Popov et al., 2023).

To characterise the effects of session, age, and sex, we conducted hierarchical linear models using *lmer* in *R* (Bates et al., 2014), applying these at the cluster level. We first ran models including the main effects of session, age, and sex, with participant ID treated as a random intercept, and mean centering age:

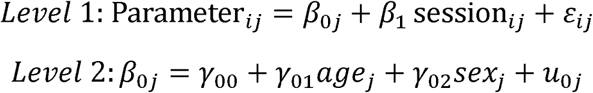

We also ran models including an interaction term between age and session, where level 1 is identical to the main effects only models:

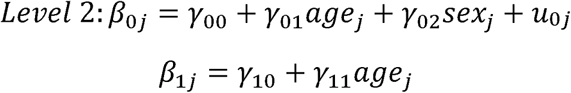

To explore potential non-linear changes in neural activity, we ran a separate set of models that included age as a quadratic term.

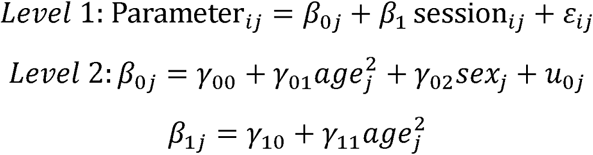

Lastly, we conducted exploratory analyses examining the interaction between session and sex.

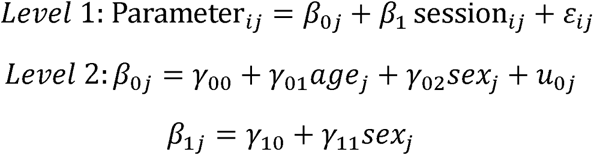

To identify which models (main effects vs interaction) provided the best fit to the data, we used the Akaike Information Criterion (AIC). To control for multiple comparisons in the linear models, we applied a Benjamini-Hochberg (Benjamini & Hochberg, 1995) false discovery rate correction within each parameter and coefficient family (e.g., eyes-open midline, eyes-closed midline, eyes-open perimeter, eyes-closed perimeter). Separate sets of corrections were applied to both hypothesis driven and exploratory models, and we report both uncorrected and corrected *p*-values.

## Results

### Test-Retest Reliability of EEG Parameters

Topographic plots of the ICC values for each measure and condition are presented in Figure 2. Across all measures and conditions, we observed fair to excellent reliability across five-years, with central sites typically yielding the strongest ICC values. Although lower than in previous reports over shorter timeframes (McKeown et al., 2024), the aperiodic exponent in eyes-closed and eyes-open conditions still had fair to good reliability. The aperiodic offset, as in previous studies, showed good reliability across frontocentral and parietal midline electrodes. Parameterized alpha power had extremely good test-retest reliability, particularly in the eyes-closed condition. The reliability of IAPF was generally good in the eyes-closed condition, but fair in the eyes-open condition.

At the cluster level, reliability was fair to excellent (Table 2). The aperiodic exponent had fair to good reliability across both midline and perimeter clusters in the eyes-closed and eyes-open conditions, respectively. The aperiodic offset had good reliability across both clusters and eyes-closed and eyes-open conditions. IAPF showed excellent reliability over the occipitoparietal cluster and good reliability over the frontotemporal cluster in the eyes-closed condition, and good reliability over both occipitoparietal and frontotemporal clusters in the eyes-open condition. Alpha power showed excellent reliability across both occipitoparietal and frontotemporal clusters in the eyes-closed condition, and good reliability across both clusters in the eyes-open condition.

**Table 2.** Test-Retest Reliability of EEG Measures by Cluster and Condition.

| Measure | Cluster | Condition | ICC | *F* | *n* |
| --- | --- | --- | --- | --- | --- |
| Aperiodic Exponent | Midline | Eyes Closed | 0.54<br>[0.43, 0.64] | 3.42 | 178 |
|  | Perimeter | Eyes Closed | 0.58<br>[0.47, 0.67] | 3.82 | 178 |
|  | Midline | Eyes Open | 0.62<br>[0.50, 0.71] | 4.44 | 161 |
|  | Perimeter | Eyes Open | 0.62<br>[0.50, 0.71] | 4.55 | 161 |
| Aperiodic Offset | Midline | Eyes Closed | 0.68<br>[0.57, 0.76] | 5.55 | 178 |
|  | Perimeter | Eyes Closed | 0.72<br>[0.62, 0.79] | 6.49 | 178 |
|  | Midline | Eyes Open | 0.67<br>[0.55, 0.76] | 5.50 | 161 |
|  | Perimeter | Eyes Open | 0.65<br>[0.53, 0.74] | 5.16 | 161 |
| IAPF | Occipitoparietal | Eyes Closed | 0.84<br>[0.79, 0.88] | 12.49 | 175 |
|  | Frontotemporal | Eyes Closed | 0.80<br>[0.74, 0.85] | 9.18 | 175 |
|  | Occipitoparietal | Eyes Open | 0.74<br>[0.66, 0.81] | 6.82 | 147 |
|  | Frontotemporal | Eyes Open | 0.65<br>[0.54, 0.73] | 4.62 | 147 |
| Alpha Power | Occipitoparietal | Eyes Closed | 0.87<br>[0.84, 0.91] | 14.95 | 175 |
|  | Frontotemporal | Eyes Closed | 0.87<br>[0.83, 0.90] | 15.16 | 175 |
|  | Occipitoparietal | Eyes Open | 0.82<br>[0.76, 0.87] | 10.15 | 147 |
|  | Frontotemporal | Eyes Open | 0.79<br>[0.72, 0.84] | 8.32 | 147 |

Note. IAPF = Individual Alpha Peak Frequency. 95% CI are presented in brackets. All *p*-values were <.001.

Taken together, these findings demonstrate that both periodic and aperiodic EEG parameters exhibit good to excellent stability over a five-year period, supporting their validity as trait-like neural markers. This suggests that rank-order differences in neural activity are maintained over time in a typically aging sample, providing a foundation for longitudinal studies of age-related changes

### Time and Age Effects on Aperiodic and Alpha Activity

Next, we examined longitudinal changes in both aperiodic and periodic alpha activity between Session 1 and Session 2 using linear mixed-effects models (Tables 3-10). We observed significant flattening of the aperiodic exponent (Tables 3-4, Figure 3), with a decrease between Session 1 and Session 2 for both the midline and perimeter clusters in eyes-closed and eyes-open conditions. We also observed consistent associations between age and the aperiodic exponent, with older adults showing lower exponents across both clusters and conditions.

**Table 3.** Hierarchical Linear Models of Aperiodic Exponent Stability (Eyes-Closed)

| Predictors | Midline |  |  |  |  |  |  | Perimeter |  |  |  |  |  |  |
| --- | --- | --- | --- | --- | --- | --- | --- | --- | --- | --- | --- | --- | --- | --- |
|  | Estimates | std. Error | std. Beta | CI | Statistic | p | p<br>(FDR.adj) | Estimates | std. Error | std. Beta | CI | Statistic | p | p<br>(FDR.adj) |
| Intercept | 1.299 | 0.043 | 0.080 | 1.213 – 1.384 | 29.896 | <0.001 |  | 1.017 | 0.042 | 0.069 | 0.933 – 1.100 | 23.968 | <0.001 |  |
| Session | -0.053 | 0.024 | -0.081 | -0.100 – -0.007 | -2.267 | 0.025 | 0.033 | -0.045 | 0.022 | -0.069 | -0.089 – -0.001 | -2.009 | 0.046 | 0.046 |
| Age | -0.007 | 0.001 | -0.301 | -0.010 – -0.004 | -4.878 | <0.001 | <0.001 | -0.005 | 0.002 | -0.229 | -0.008 – -0.002 | -3.542 | 0.001 | 0.001 |
| Sex | -0.076 | 0.043 | -0.230 | -0.161 – 0.009 | -1.774 | 0.078 | 0.295 | -0.064 | 0.044 | -0.197 | -0.152 – 0.023 | -1.455 | 0.147 | 0.295 |
| **Random Effects** |  |  |  |  |  |  |  |  |  |  |  |  |  |  |
| $\sigma^2$ | 0.05 | | | | | | | 0.04 | | | | | | |
| $\tau_{00}$ | 0.05<sub>participant_id</sub> | | | | | | | 0.06<sub>participant_id</sub> | | | | | | |
| ICC | 0.50 |  |  |  |  |  |  | 0.56 |  |  |  |  |  |  |
| N | 178<sub>participant_id</sub> |  |  |  |  |  |  | 178<sub>participant_id</sub> |  |  |  |  |  |  |
| Observations | 356 |  |  |  |  |  |  | 356 |  |  |  |  |  |  |
| Marginal R<sup>2</sup> / Conditional R<sup>2</sup> | 0.116 / 0.554 |  |  |  |  |  |  | 0.071 / 0.590 |  |  |  |  |  |  |
| AIC | 168.574 |  |  |  |  |  |  | 160.304 |  |  |  |  |  |  |

Note: Repeated-measured EEG data nested within participant, indicated by participant\_id. Sex coded as 0 = female, 1 = male.

**Table 4.** Hierarchical Linear Models of Aperiodic Exponent Stability (Eyes-Open)

| Predictors | Midline |  |  |  |  |  |  | Perimeter |  |  |  |  |  |  |
| --- | --- | --- | --- | --- | --- | --- | --- | --- | --- | --- | --- | --- | --- | --- |
|  | Estimates | std. Error | std. Beta | CI | Statistic | p | p<br>(FDR.adj) | Estimates | std. Error | std. Beta | CI | Statistic | p | p<br>(FDR.adj) |
| Intercept | 1.275 | 0.037 | 0.053 | 1.201 – 1.348 | 34.196 | <0.001 |  | 0.931 | 0.036 | -0.026 | 0.859 – 1.002 | 25.682 | <0.001 |  |
| Session | -0.070 | 0.019 | -0.122 | -0.108 – -0.032 | -3.629 | <0.001 | 0.001 | -0.069 | 0.018 | -0.125 | -0.105 – -0.033 | -3.749 | <0.001 | 0.001 |
| Age | -0.006 | 0.001 | -0.287 | -0.009 – -0.003 | -4.248 | <0.001 | <0.001 | -0.004 | 0.001 | -0.213 | -0.007 – -0.002 | -3.053 | 0.003 | 0.003 |
| Sex | -0.044 | 0.041 | -0.155 | -0.125 – 0.036 | -1.093 | 0.276 | 0.368 | 0.021 | 0.041 | 0.077 | -0.059 – 0.101 | 0.525 | 0.601 | 0.601 |
| **Random Effects** |  |  |  |  |  |  |  |  |  |  |  |  |  |  |
| $\sigma^2$ | 0.03 | | | | | | | 0.03 | | | | | | |
| $\tau_{00}$ | 0.04<sub>participant_id</sub> | | | | | | | 0.05<sub>participant_id</sub> | | | | | | |
| ICC | 0.60 |  |  |  |  |  |  | 0.63 |  |  |  |  |  |  |
| N | 161<sub>participant_id</sub> |  |  |  |  |  |  | 161<sub>participant_id</sub> |  |  |  |  |  |  |
| Observations | 322 |  |  |  |  |  |  | 322 |  |  |  |  |  |  |
| Marginal R<sup>2</sup> / Conditional R<sup>2</sup> | 0.106 / 0.641 |  |  |  |  |  |  | 0.060 / 0.648 |  |  |  |  |  |  |
| AIC | 39.256 |  |  |  |  |  |  | 23.615 |  |  |  |  |  |  |

Note: Repeated-measured EEG data nested within participant, indicated by participant\_id. Sex coded as 0 = female, 1 = male.

**Figure 3.**
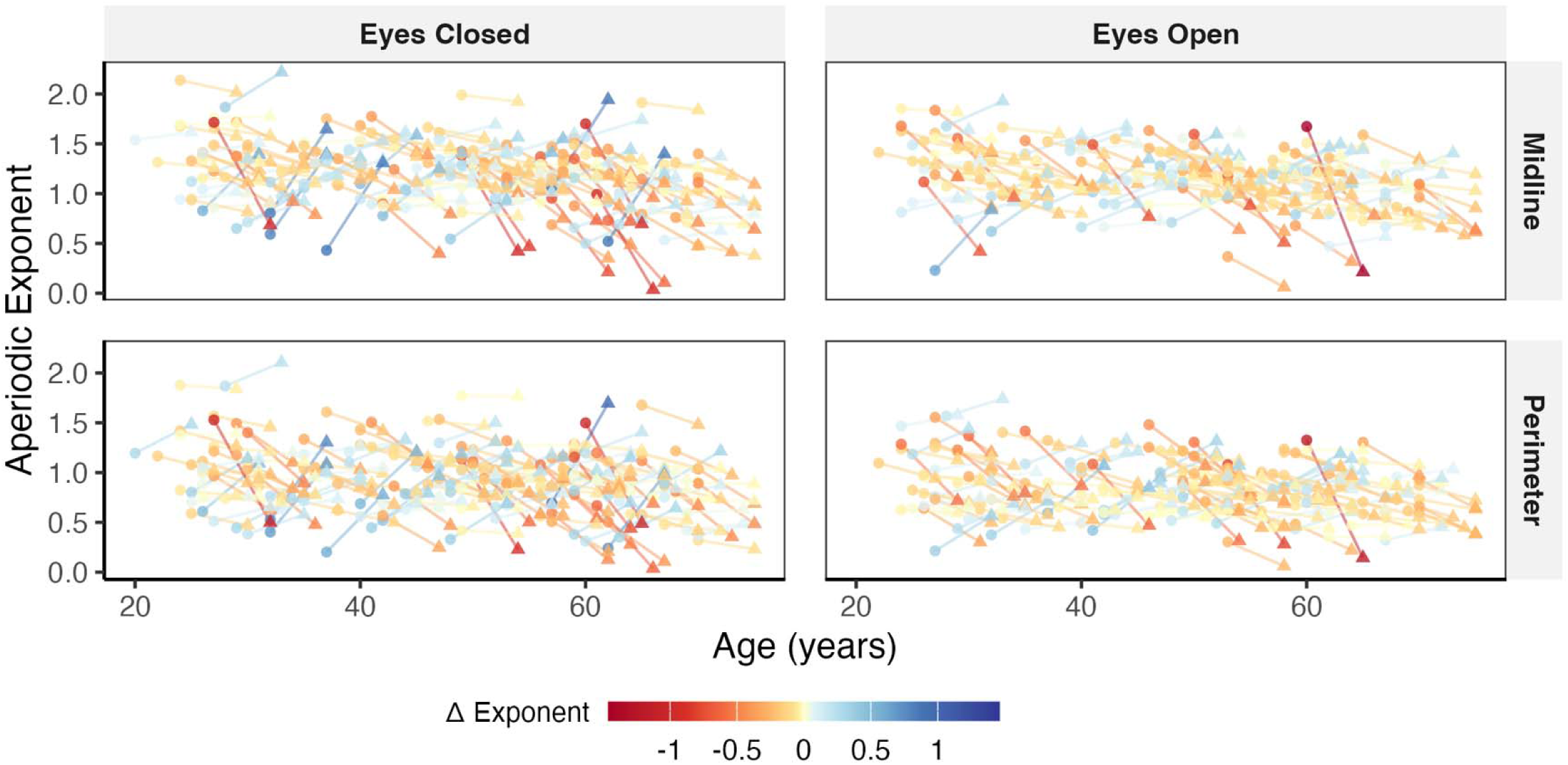
Aperiodic exponent for midline and perimeter clusters in each session. Note. Circles indicate each participant’s measure at session 1, and triangles indicate their measure five years later at session 2. Points and lines are shaded to show the direction and magnitude of change between session 1 and session 2.

For the aperiodic offset, we also observed decreases over time and negative associations with age (Tables 5-6, Figure 4), with these effects present across both clusters and both eyes-open and eyes-closed conditions. We also observed main effects of sex: male participants had lower aperiodic offsets in the midline cluster and the perimeter cluster in eyes-closed conditions, and in the midline cluster in eyes-open conditions, but not in the perimeter cluster.

**Table 5.**
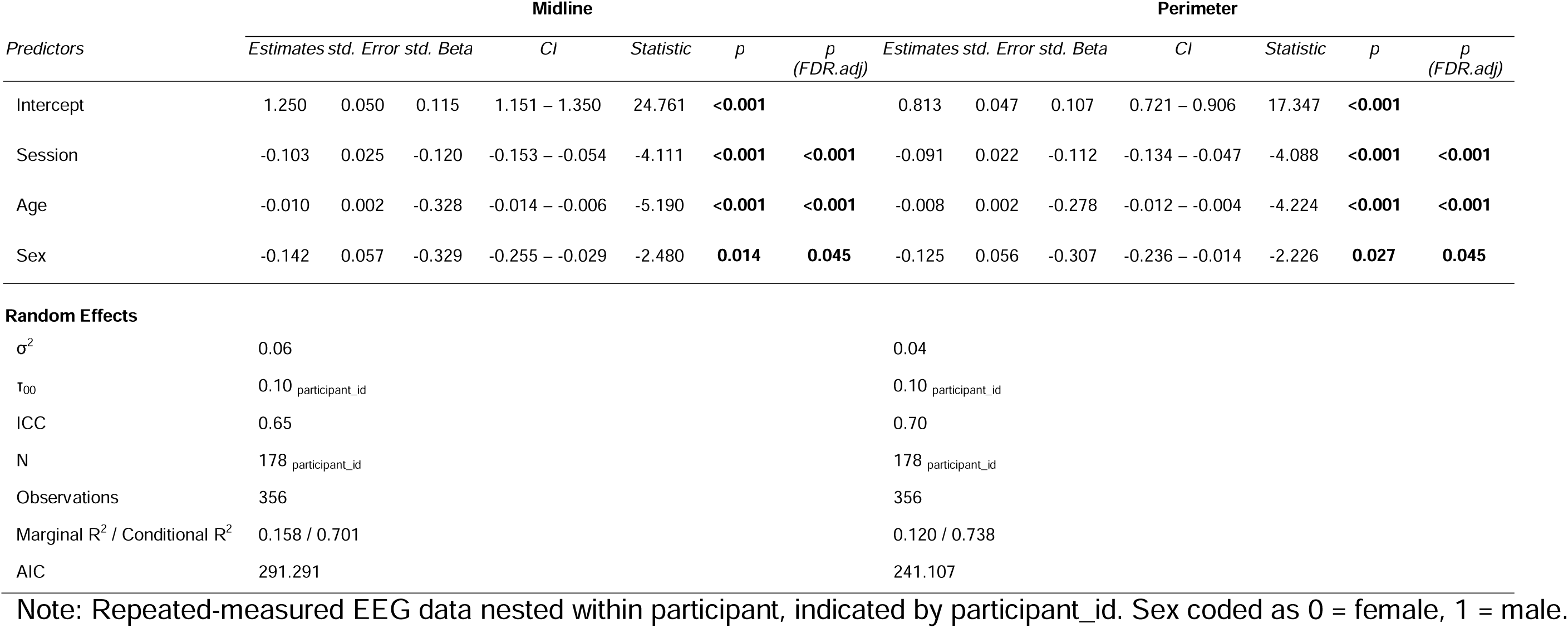
Hierarchical Linear Models of Aperiodic Offset Stability (Eyes-Closed)

**Table 6.**
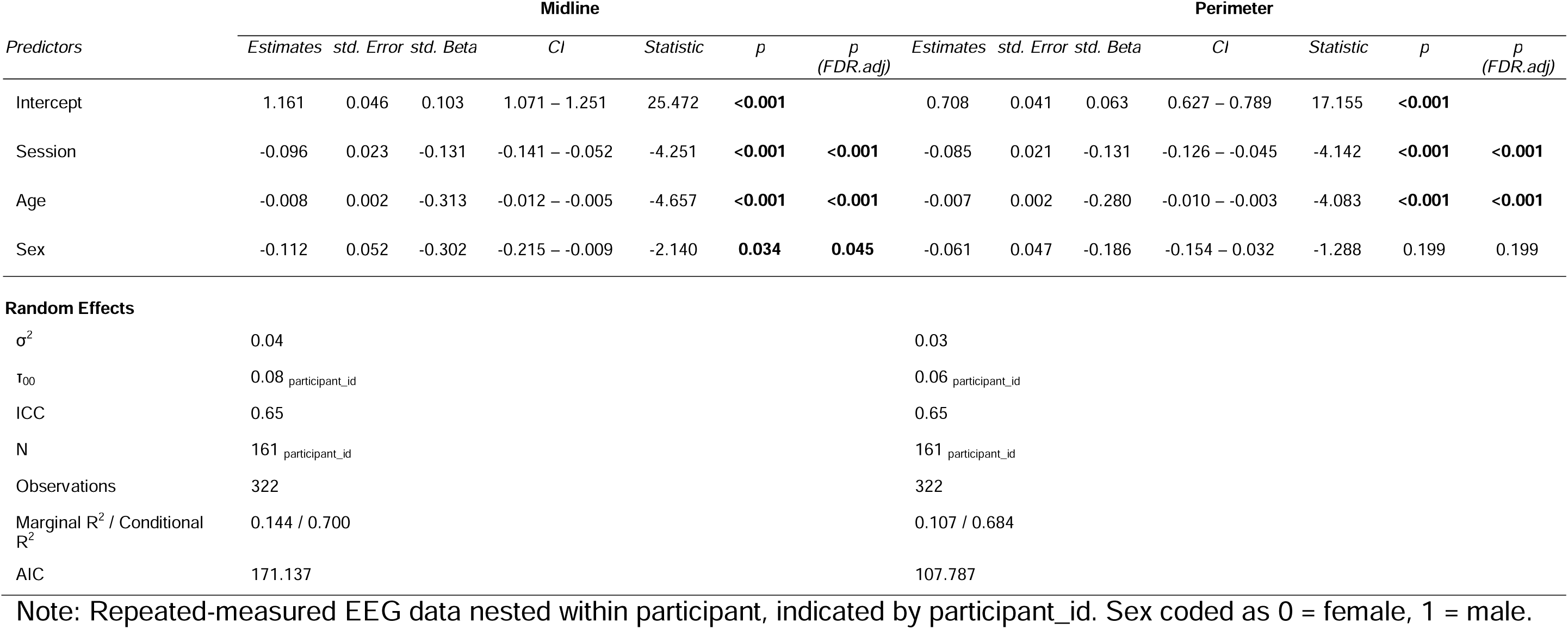
Hierarchical Linear Models of Aperiodic Offset Stability (Eyes-Open)

**Figure 4.**
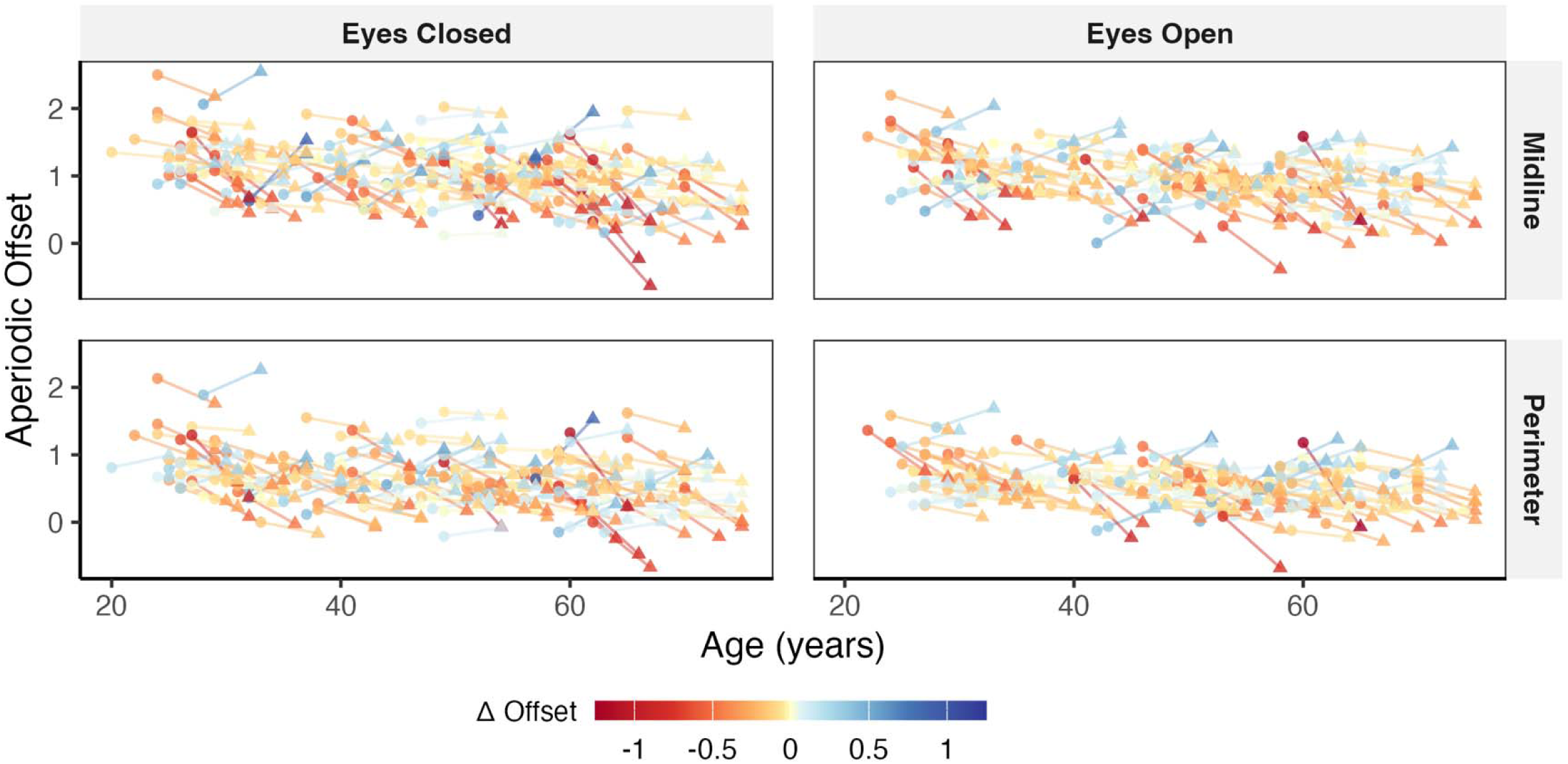
Aperiodic offset for midline and perimeter clusters in each session. Note. Circles indicate each participant’s measure at session 1, and triangles indicate their measure five years later at session 2. Points and lines are shaded to show the direction and magnitude of change between session 1 and session 2.

Individual Alpha Peak Frequency also showed significant decreases over time, such that IAPF was slower at Session 2 compared to Session 1 for the occipitoparietal cluster in eyes-closed and eyes-open conditions, but not for the frontotemporal cluster, once adjusting for FDR (Tables 7-8, Figure 5). IAPF was also negatively associated with age, with older adults showing slower central frequencies than younger adults in the occipitoparietal cluster during eyes-closed but not eyes-open, and in the frontotemporal cluster during eyes-closed and eyes-open. Lastly, although we observed a significant time effect on alpha power in the frontotemporal cluster during the eyes-closed condition, this did not survive FDR correction (Tables 9-10, Figure 6). There were no other significant effects (before, or after FDR correction).

**Table 7.**
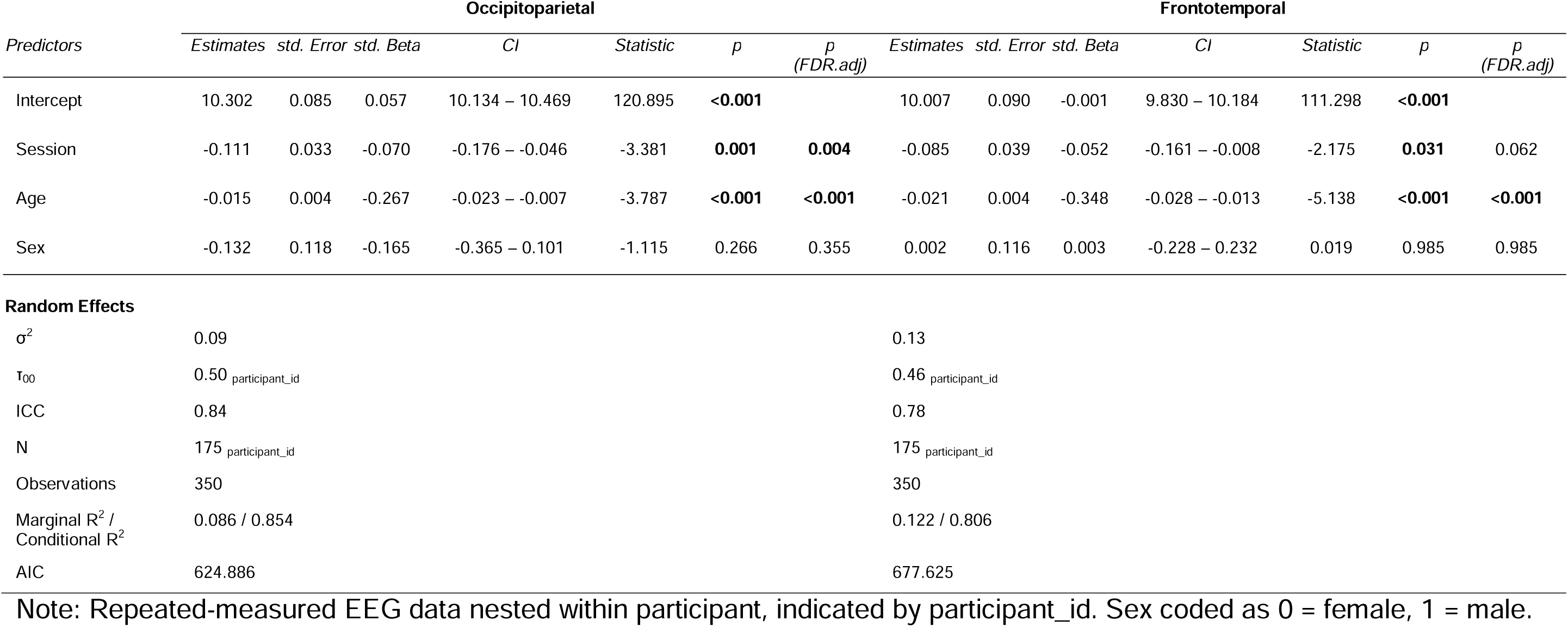
Hierarchical Linear Models of Individual Alpha Peak Frequency Stability (Eyes-Closed)

**Table 8.**
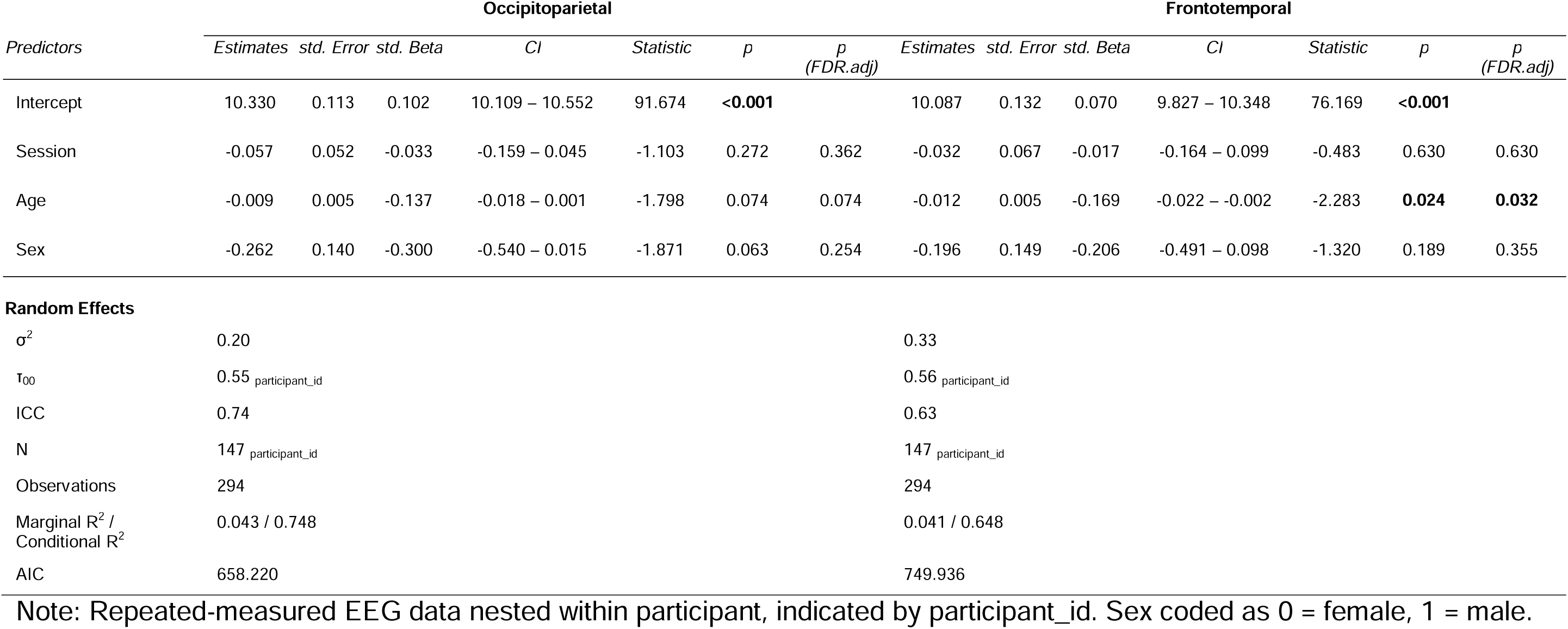
Hierarchical Linear Models of Individual Alpha Peak Frequency Stability (Eyes-Open)

**Figure 5.**
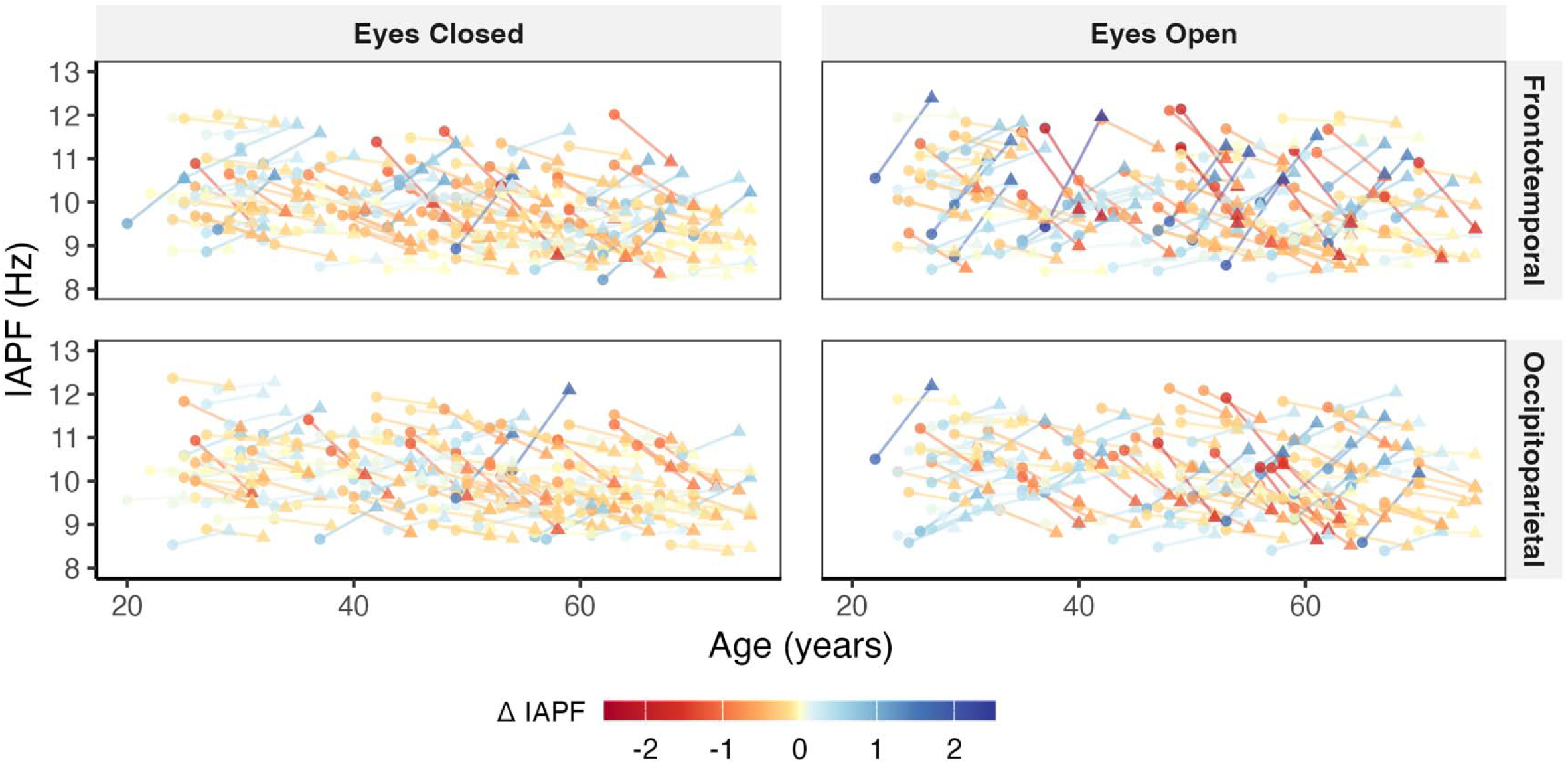
Individual alpha peak frequency for occipitoparietal and temporal clusters in each session. Note. Circles indicate each participant’s measure at session 1, and triangles indicate their measure five years later at session 2. Points and lines are shaded to show the direction and magnitude of change between session 1 and session 2.

**Table 9.**
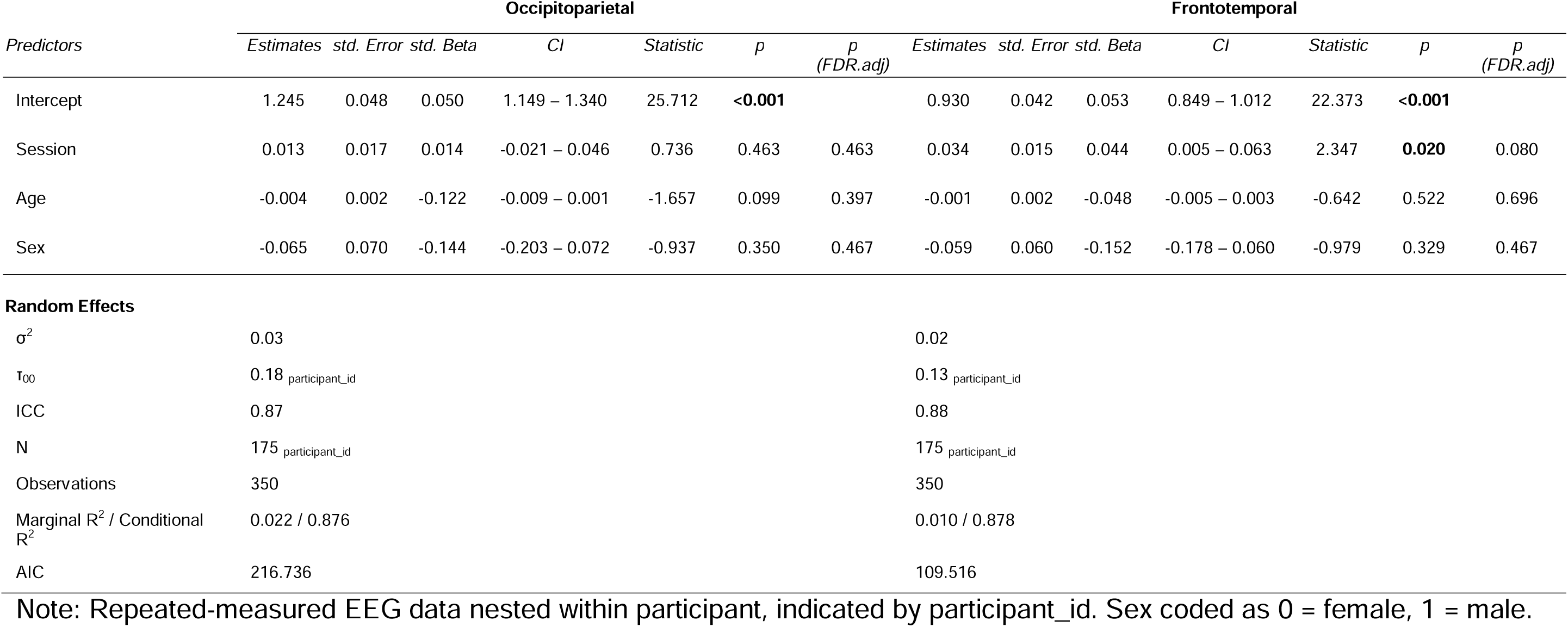
Hierarchical Linear Models of Alpha Power Stability (Eyes-Closed)

**Table 10.**
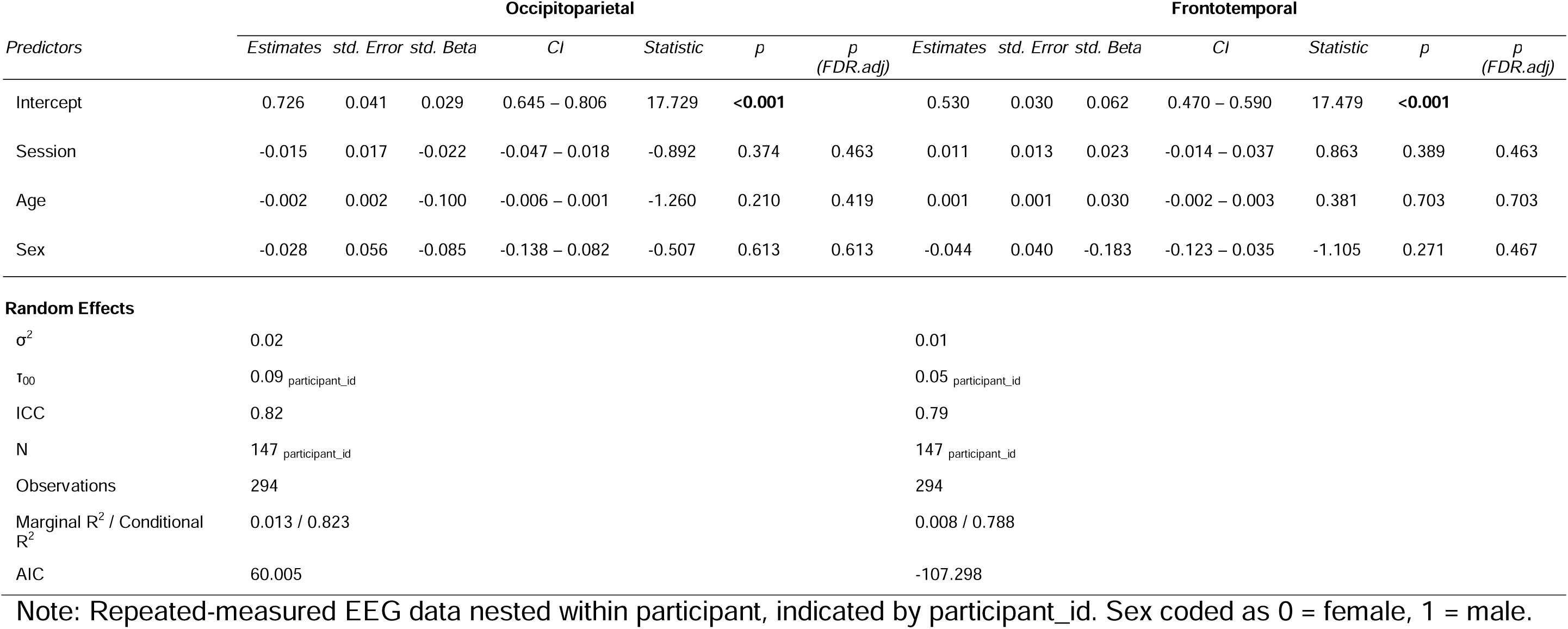
Hierarchical Linear Models of Alpha Power Stability (Eyes-Open)

**Figure 6.**
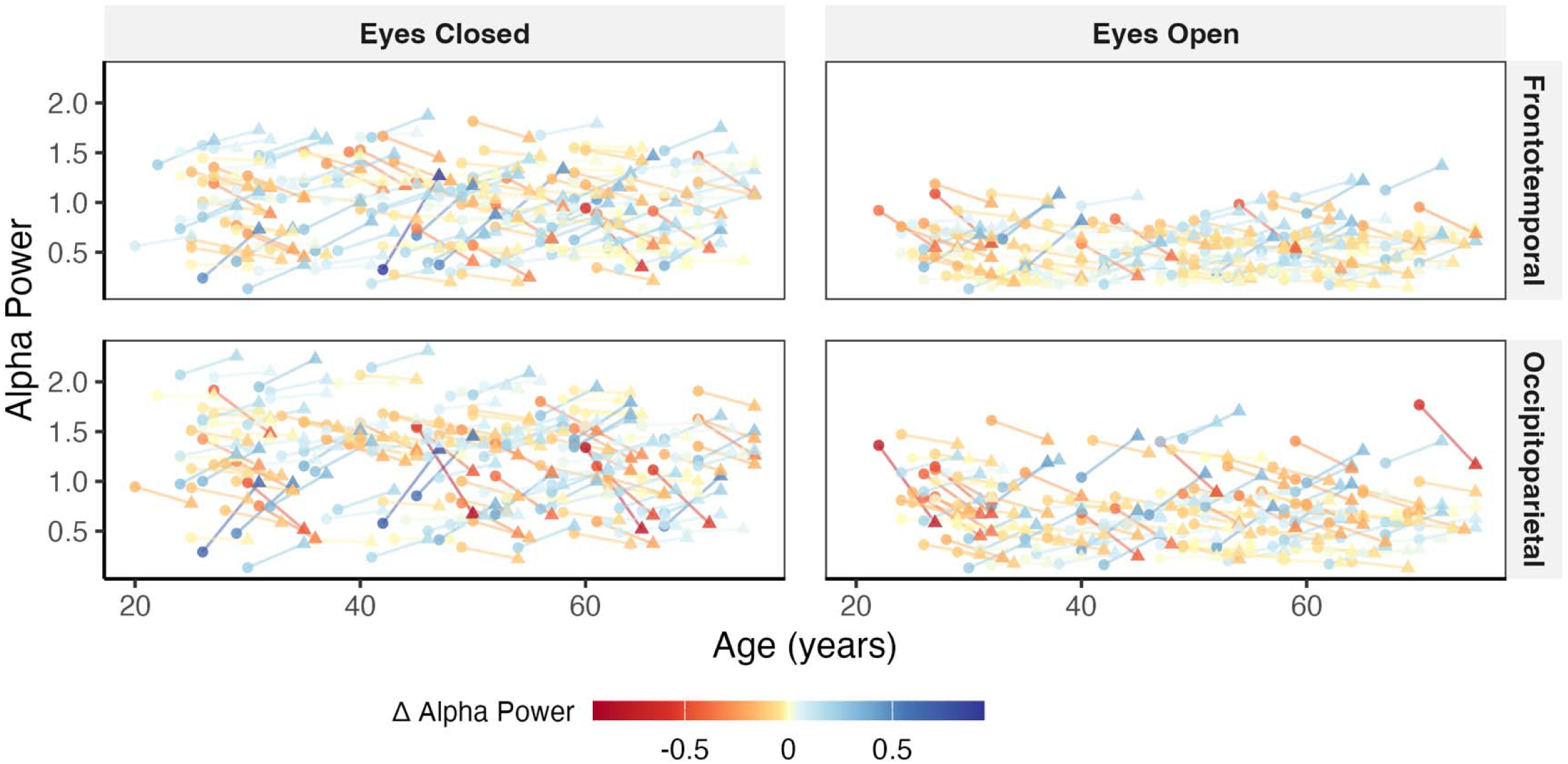
Alpha power for occipitoparietal and temporal clusters in each session. Note. Circles indicate each participant’s measure at session 1, and triangles indicate their measure five years later at session 2. Points and lines are shaded to show the direction and magnitude of change between session 1 and session

In the analyses where we observed significant independent effects of both session and age, the effect of age (std. Beta ranges = −0.213 – −0.348) tended to be larger in magnitude than the effect of session (std. Beta ranges = −0.052 – −0.213).

As shown in the supplementary info, we did not observe any interaction between session and age, session and age^2^, or session and sex, suggesting that the time-related changes in aperiodic and periodic activity we observed were not affected by participants’ age (when treated as linear or quadratic) at Session 1, or their sex. That is, we did not observe patterns consistent with a model of accelerated change in aperiodic or periodic activity with age, or increased (or decreased) change as a function of participant sex. The inclusion of the interaction term removed most significant main effects, except for session for the exponent (midline and perimeter clusters, eyes-open condition), for session, age and sex for the offset (midline and perimeter clusters, eyes-closed and eyes-open conditions), and session and age for IAPF (frontotemporal cluster, eyes-closed condition). However, in all instances, the AIC was lowest for the main-effects-only models, suggesting that these provide a better fit to the data than the interaction-term models.

## Discussion

We examined the stability of aperiodic and periodic across five years in a sample of younger, midlife, and older adults. Four central findings emerged. First, all parameters demonstrated fair to excellent test-retest reliability. Second, older adults exhibited lower aperiodic exponents and slower IAPF than younger adults in longitudinal models. Third, alpha power did not change over time, despite significant decreases in the aperiodic exponent, aperiodic offset, and IAPF. Fourth, and contrary to our hypotheses, we found no evidence of greater decreases in the aperiodic exponent, aperiodic offset, IAPF, or alpha power in older (relative to younger) adults.

### Test-Retest Reliability of EEG Parameters

The observed test–retest reliability across all spectral parameters supports their use as stable potential biomarkers, with particularly high reliability seen for IAPF and alpha power. Importantly, our finding that the aperiodic exponent and aperiodic offset show fair reliability over a five-year time frame provides strong support for their temporal stability and extends previous short-term findings (McKeown et al., 2024; Pathania et al., 2021). Although likely sensitive to methodological factors such as data quality, analysis parameters, and spectral-fitting choices (Donoghue et al., 2020; Ostlund et al., 2022), aperiodic activity nonetheless showed consistent temporal stability across five years, supporting its use as a non-invasive and longitudinal marker of E:I balance and neural spiking rates. Unlike our previous work (McKeown et al., 2024), which found that the test-retest reliability of the aperiodic exponent is poorer in the eyes-open than in the eyes-closed condition, here we observed the opposite.

We observed some variation in the reliability of alpha parameters across topography and condition. The reliability of alpha power was slightly lower in the eyes-open condition (particularly over frontotemporal sensors), consistent with previous studies showing that environmental variability and attentional demands reduce alpha power consistency (Barry et al., 2007; Cooper et al., 2003). Notably, even the reduced eyes-open alpha power reliability exceeds the fair reliability (ICC > 0.60). These findings align with prior research demonstrating that periodic features, especially IAPF, reflect enduring individual differences in neural architecture and cognitive functioning that are largely independent of momentary state fluctuations (Grandy, Werkle-Bergner, Chicherio, Schmiedek, et al., 2013). These reliability coefficients exceed conventional thresholds for excellent reliability in clinical contexts (Cicchetti, 1994).

### Modelling Changes in Aperiodic and Periodic Activity

Our findings replicate and extend the findings observed in previous cross-sectional datasets, though to reflect a range of developmental and, in older adults, pathophysiological changes. Variation in alpha frequency has been linked to changes in white-mater integrity (Babiloni et al., 2008; Kramberger et al., 2017), while variation in the aperiodic exponent and offset has been attributed to changes in a variety of mechanisms that impact E:I balance and spiking rates (e.g., gamma-aminobutyric acid receptor gene expression; Duma et al., 2024; amyloid-beta and tau proteins; Martínez-Cañada et al., 2023; pathophysiological alterations to GABAergic neurons; Wiest et al., 2023). Collectively, these associations have been conceptualized as reflecting a gradual loss of coordination in neural activity. Parameterized measures of alpha power might be expected to differ from absolute power measures in older participants, given that age-related alterations in peak alpha frequency and in the aperiodic exponent can both influence power within and adjacent to the alpha band (Finley et al., 2022; Klimesch, 1999; Merkin et al., 2022). We, however, did not observe an association between age and alpha power, including when we examined absolute (rather than parameterized) power (see supplementary info, tables S36-S45), suggesting that the stability of alpha power in our study reflects features of the sample rather than an artefact of how alpha was measured.

Previous studies have been limited by their reliance on data covering short time scales, with an exclusive focus on reliability within those intervals. Others have examined changes in activity across early life development, finding increases in the aperiodic exponent and offset in infancy (Schaworonkow & Voytek, 2021) and early childhood (Sacks et al., 2025). Others have reported decreases in the aperiodic exponent and offset over two years in adolescents (McSweeney et al., 2021). Beyond this, research has been limited to cross-sectional analyses of age (Finley et al., 2022; Hill et al., 2022; Merkin et al., 2022), leaving it unclear whether these findings reflect highly replicable cohort effects or changes in neural activity, underlying cortical structures, and function. Here, for the first time, we show that in addition to a negative association with age, aperiodic activity changes longitudinally over the lifespan. The aperiodic exponent flattened across the five years between sessions and the aperiodic offset decreased in height. The absence of any significant age and session interactions for these measures is consistent with, but does not confirm, that these patterns of change do not accelerate with age. Although changes between two sessions cannot be uniquely attributed to chronological ageing rather than to other time-varying factors, these longitudinal changes are consistent with theoretical accounts linking lower exponents to reduced E:I balance and diminished synchronization of neural population activity with advancing age (Gao et al., 2017; Voytek et al., 2015). The concurrent decrease in aperiodic offset may reflect reductions in overall neural firing rates or broadband power, potentially indexing declines in cortical metabolic activity or neuronal density between sessions.

We also observed clear changes in parameterized alpha activity, although this was limited to IAPF. The observed slowing of IAPF is consistent with previous cross-sectional and longitudinal studies demonstrating reductions in alpha peak frequency with age (Klimesch, 1999), which are thought to reflect changes in thalamocortical network dynamics and white matter integrity. We, however, did not observe changes in peak alpha power across sessions. The stability of alpha power observed in our study appears to contradict extensive literature reporting age-related decreases in power (Babiloni et al., 2006). Previous studies typically examined alpha power within fixed frequency bands (e.g., 8–12 Hz), which would systematically underestimate power in individuals with slower IAPF, as conventional spectral boundaries may be biased against older adults or any group with a slowed IAPF (Scally et al., 2018). However, in exploratory analyses examining absolute alpha power, we observed the same pattern of stability, suggesting that this null effect is a function of the sample, rather than potential IAPF-related confounds.

Lastly, we observed differences in aperiodic activity by participant sex. Even after controlling for age, male participants tended to have lower offsets than female participants. Few other studies have specifically examined sex or gender differences in aperiodic activity, and differences appear specific to developmental stages. In infants and young children, females have been found to have higher aperiodic exponents and lower offsets than males (Sacks et al., 2025). In adolescents, females tend to have lower aperiodic exponents and offsets than males (McSweeney et al., 2021). The inconsistent direction of sex differences across developmental stages may relate to sex-based variation in cortical and subcortical structure and organization, which may differently influence E:I balance and spiking rates across the lifespan (Koolschijn & Crone, 2013; Lenroot & Giedd, 2010).

### Implications, Limitations and Future Directions

Our findings have several implications for the use of parameterised activity in cross-sectional and longitudinal research. First, the acceptable-to-excellent reliability we report over a five-year period provides additional support for research examining individual differences in these metrics as correlates or predictors of aging, cognitive change, and age-related pathophysiology. Second, our data provide a benchmark for power analysis for both cross-sectional and longitudinal studies examining paramatiersed activity (Hedge et al., 2018). Third, albeit beyond the scope of our study, our findings that aperiodic metrics have fair or greater relevance support their use as intervention targets. Fourth, our observation that the aperiodic exponent, aperiodic offset, and IAPF decreased between sessions, whereas peak alpha power does not, suggests that the neurophysiological processes underlying these metrics may follow different developmental trajectories in adulthood.

There are several limitations to our work. First, approximately 1/3^rd^ of participants had data for both sessions, and these participants were typically older and had greater alpha power compared to participants with data for only session 1. Because these data derive from an initial public release of an ongoing study (Gajewski et al., 2022; Getzmann et al., 2024), these differences are a property of the dataset, which focused only on participants who could have returned for session 2 between 2021 and 2024. As data collection for session 1 occurred between 2016 and 2023, the tranche of data released in Getzmann et al. (2024) only includes session 2 data from participants who completed their session 1 visits between 2016 and 2019. These differences, combined with possible sample attrition, may bias our estimates of reliability, and associations with age. ICCs may be biased upwards as participants with better signal quality may be more likely to return for the second session. Cross-sectional activity–age associations may be more conservative than reported if, among older participants, those healthy enough to return for session 2 represent a positively selected subset relative to non-returners: the differences between younger and older participants may therefore underestimate the differences that would be observed in a fully representative sample. Replication in future waves of the Dortmund Vital Study (Gajewski et al., 2022) will be necessary to establish generalizability, and more precise estimates of cohort-level reliability and change.

Second, because the current tranche of data includes only two sessions, we are only able to establish that change occurred between sessions, but cannot characterize the shape of this change as linear, accelerating, decelerating, or non-monotonic. Although we did not observe a significant interaction between age and session, this indicates that the change across sessions was not significantly moderated by age, but does not establish linearity. We are also unable to rule out regression to the mean as a potential explanation for change between sessions. Third, the available data provide only limited information about participants. It is possible that changes in lifestyle, health condition, and various environmental factors could contribute to both the reliability and stability of the metrics we examined. Similarly, potential age-related changes are inherently confounded with potential session-related changes. That is, beyond alteration in neural activity that can be attributed to chronological age, there are multiple individual, environmental (e.g., COVID-19; Jiang et al., 2024), and recording factors that differ between sessions, which could conceivably bias our results. Although the reported protocol was held constant across sessions, the impact of unmeasured sources of variance cannot be ruled out. The session-effect changes we observed are, however consistent with cross-sectional age associations, but this is convergent, rather than direct evidence.

Finally, although cross-sectional associations between aperiodic activity and cognitive performance are well documented, we were unable to examine them in the current data. It therefore remains unclear whether within-person changes in these parameters predict subsequent cognitive trajectories.

## Conclusion

Recent advances in spectral parameterization techniques have enabled researchers to disentangle aperiodic and periodic components of neural activity, linking these measures to constructs that were previously difficult to quantify non-invasively (e.g., E:I balance), and establishing their promise as markers of ageing, neurodegeneration, and psychiatric illness. However, realizing this potential requires establishing the psychometric properties of these measures and understanding how they change over time. Building on work evaluating reliability over short timescales, we showed that aperiodic activity and parameterized alpha activity demonstrate acceptable levels of reliability over a five-year period in younger, midlife, and older adults. Critically, this reliability was observed despite concurrent associations with age and within-person changes in these metrics over time. These findings support the viability of aperiodic and periodic parameters for use in longitudinal research, particularly when derived using spectral parameterization approaches such as SpecParam. However, clear differences in reliability across neural measures parameterized using the same approach highlight the need to carefully consider which parameters are best suited to specific research questions and populations.

## Acknowledgements

Thanks: We thank Getzmann and colleagues for making the data available, and the participants for volunteering their time.

## Author contributions

Conceptualization: Polina Politanskaia, Jacinta Bywater, Anna J. Finley, Hannah A. D. Keage, Nicholas J. Kelley, Daniel McKeown, Victor Schinazi, and Douglas J. Angus.

Data curation: Douglas J. Angus.

Formal analysis: Polina Politanskaia and Douglas J. Angus.

Methodology: Polina Politanskaia and Douglas J. Angus.

Project administration: Douglas J. Angus.

Software: Douglas J. Angus.

Supervision: Douglas J. Angus.

Validation: Polina Politanskaia and Douglas J. Angus.

Visualization: Douglas J. Angus.

Writing - original draft: Polina Politanskaia and Douglas J. Angus.

Writing - review & editing: Polina Politanskaia, Jacinta Bywater, Anna J. Finley, Hannah A. D. Keage, Nicholas J. Kelley, Daniel McKeown, Victor Schinazi, and Douglas J. Angus.

## Footnotes

1 We opted for this approach rather than exclusion at the list-wise level. Many participants did not have clear alpha peaks in the eyes-open condition, and a list-wise approach would have reduced the analysable sample for all analyses, rather than just those involving individual peak alpha frequency, or alpha amplitude.

2 To ensure that ROIs were identical across sessions, we opted to apply the clusters identified for session one data to session two data. The clusters identified for session two alone were, however, largely identical to session one clusters.

