## Supplementary Info for "Long-term reliability and stability of parameterized resting state EEG: Evidence from a five-year follow-up"

**Silhouette Scores**

Silhouette scores for the k-means clustering analyses are presented in Table S1. Although the scores for k=2 solutions were lower than are typically recommended for clustering analyses, k=2 solutions were consistently optimal across all parameters, conditions, and sessions, converging with two-component solutions indicated by the PCA. Moreover, the resulting clusters were generally consistent with the topography of scalp recorded EEG (e.g., midline dominant aperiodic activity). The modest absolute silhouette values are not surprising for scalp EEG given volume conduction, and the clustering was used to define topographically interpretable regions of interest rather than to identify discrete, unique subgroups of channels.

**Table S1**

*Silhouette Scores for each Measure, Condition, and Session.*

| Measure | Condition | Session | k=2 | k=3 | k=4 | k=5 | k=6 | k=7 | k=8 | k=9 | k=10 |
| --- | --- | --- | --- | --- | --- | --- | --- | --- | --- | --- | --- |
| Exponent | EC | 1 | 0.33 | 0.28 | 0.23 | 0.26 | 0.20 | 0.22 | 0.21 | 0.20 | 0.19 |
| Exponent | EC | 2 | 0.32 | 0.27 | 0.26 | 0.18 | 0.18 | 0.16 | 0.18 | 0.17 | 0.16 |
| Exponent | EO | 1 | 0.34 | 0.29 | 0.18 | 0.16 | 0.13 | 0.14 | 0.14 | 0.12 | 0.13 |
| Exponent | EO | 2 | 0.34 | 0.23 | 0.22 | 0.17 | 0.17 | 0.13 | 0.14 | 0.14 | 0.15 |
| Offset | EC | 1 | 0.40 | 0.30 | 0.21 | 0.24 | 0.24 | 0.24 | 0.22 | 0.20 | 0.18 |
| Offset | EC | 2 | 0.38 | 0.24 | 0.21 | 0.23 | 0.23 | 0.23 | 0.22 | 0.23 | 0.18 |
| Offset | EO | 1 | 0.38 | 0.27 | 0.17 | 0.21 | 0.21 | 0.20 | 0.18 | 0.16 | 0.16 |
| Offset | EO | 2 | 0.38 | 0.23 | 0.20 | 0.21 | 0.21 | 0.18 | 0.19 | 0.17 | 0.16 |
| IAPF | EC | 1 | 0.18 | 0.15 | 0.10 | 0.12 | 0.11 | 0.07 | 0.07 | 0.07 | 0.08 |
| IAPF | EC | 2 | 0.18 | 0.13 | 0.11 | 0.10 | 0.07 | 0.08 | 0.07 | 0.07 | 0.08 |
| IAPF | EO | 1 | 0.14 | 0.11 | 0.13 | 0.09 | 0.08 | 0.08 | 0.07 | 0.07 | 0.07 |
| IAPF | EO | 2 | 0.13 | 0.11 | 0.12 | 0.09 | 0.09 | 0.08 | 0.08 | 0.07 | 0.07 |
| Alpha power | EC | 1 | 0.45 | 0.31 | 0.27 | 0.25 | 0.28 | 0.28 | 0.21 | 0.21 | 0.20 |
| Alpha power | EC | 2 | 0.47 | 0.28 | 0.25 | 0.23 | 0.21 | 0.22 | 0.19 | 0.18 | 0.17 |
| Alpha power | EO | 1 | 0.42 | 0.26 | 0.24 | 0.20 | 0.18 | 0.18 | 0.17 | 0.12 | 0.14 |
| Alpha power | EO | 2 | 0.40 | 0.23 | 0.18 | 0.12 | 0.12 | 0.11 | 0.14 | 0.13 | 0.14 |
| Absolute alpha power | EC | 1 | 0.38 | 0.34 | 0.29 | 0.30 | 0.29 | 0.29 | 0.26 | 0.22 | 0.23 |
| Absolute alpha power | EC | 2 | 0.38 | 0.35 | 0.31 | 0.28 | 0.26 | 0.28 | 0.30 | 0.24 | 0.26 |
| Absolute alpha power | EO | 1 | 0.38 | 0.26 | 0.28 | 0.30 | 0.28 | 0.24 | 0.25 | 0.24 | 0.22 |
| Absolute alpha power | EO | 2 | 0.36 | 0.28 | 0.26 | 0.29 | 0.27 | 0.27 | 0.21 | 0.20 | 0.18 |

Note; EC = eyes closed, EO = eyes open.

**Absolute Alpha Power**

We also performed parallel analyses on absolute alpha power. That is, power over the entire canonical alpha band, without adjusting for aperiodic activity. These analyses also yielded a two-cluster solution, which resembled the midline vs. perimeter clustering observed for the aperiodic exponent and aperiodic offset (Figure S1).

ICCs for absolute alpha power at the channel level are presented in Figure S2, and cluster level are presented in Table S2. As with parameterized alpha metrics, absolute alpha power generally had excellent reliability across Session 1 and Session2 measures for both eyes-open and eyes-closed conditions, with greater ICCs during eyes-closed.

**Figure S1**

*Topographic plots showing cluster solutions for Absolute Alpha Power.*

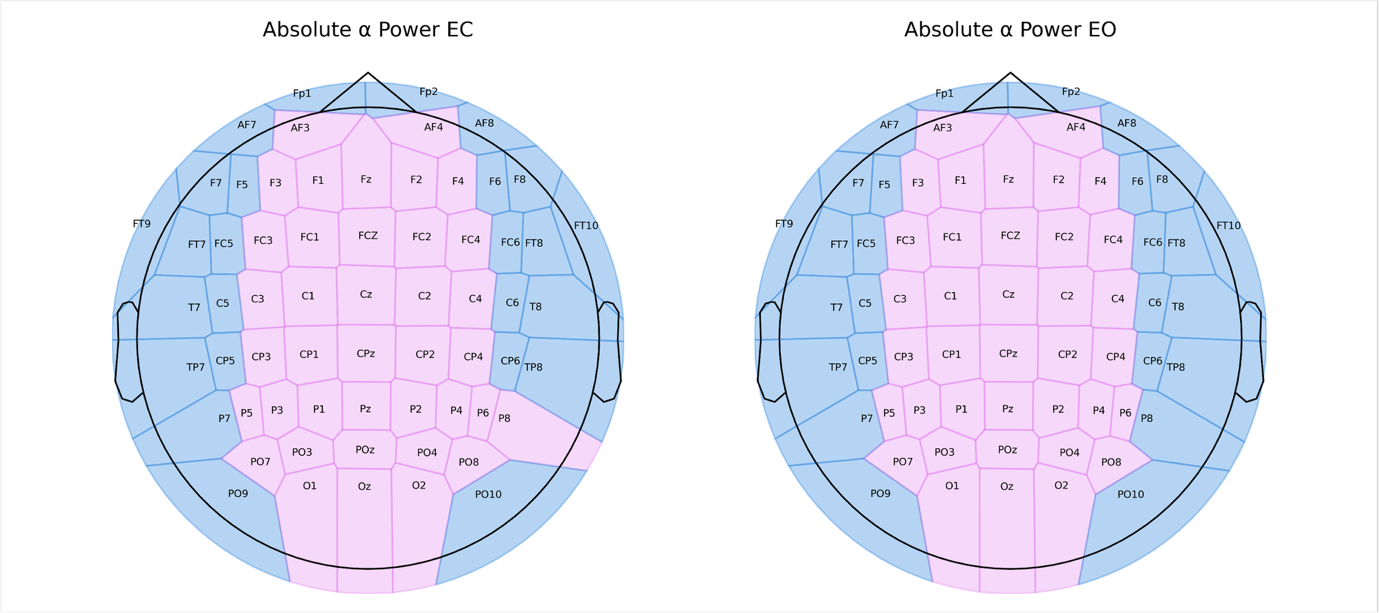

**Figure S2**

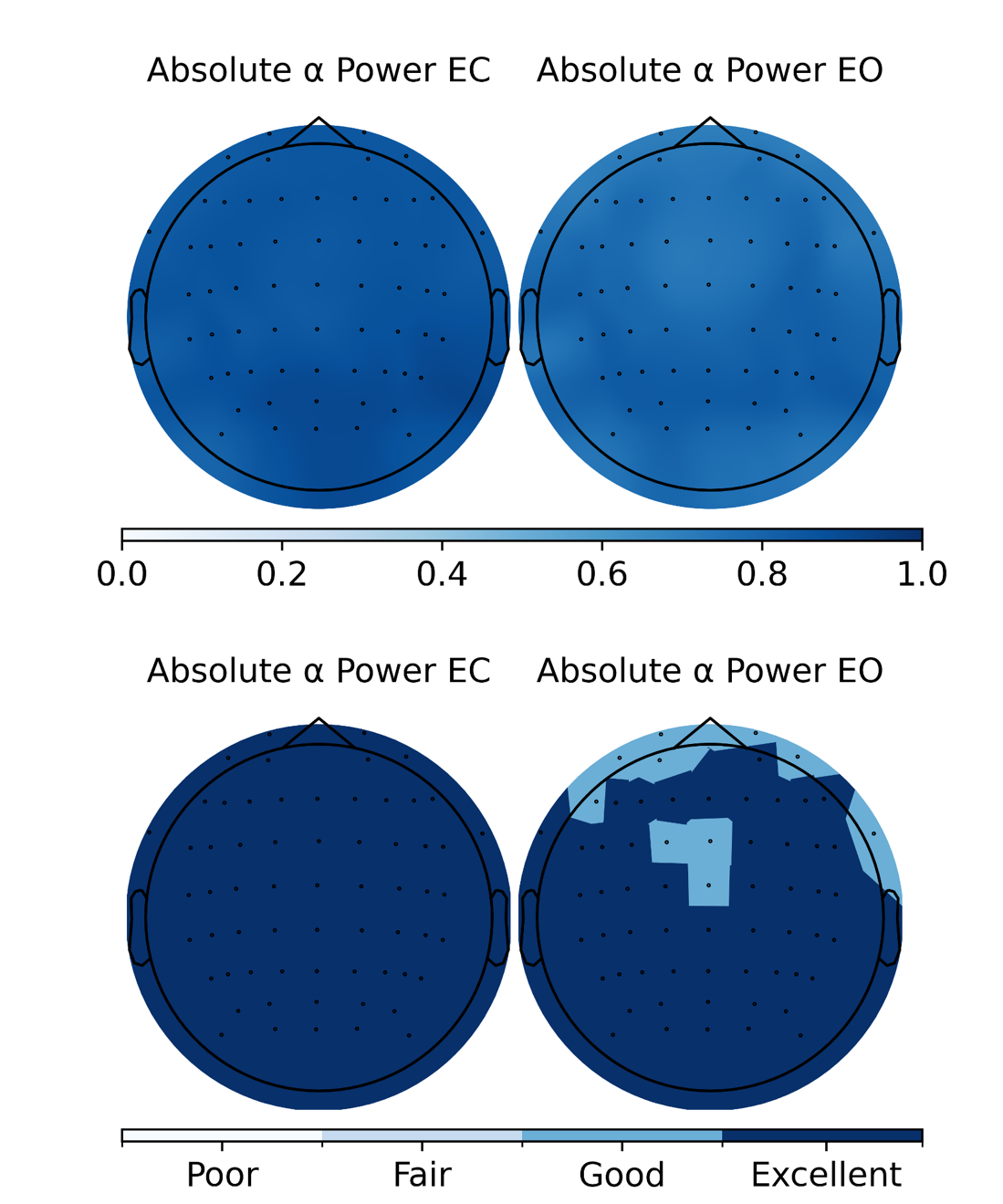
*Topographic plots depicting Intraclass Correlations for the Absolute Alpha Power*

Note: The upper panel shows individual ICCs at the channel level. The lower panel shows ICCs at the channel level, after binning into thresholds (poor = 0 – .39, Fair = .40 – .59, Good = .60 – .74, Excellent = .75 – 1.00). EC = eyes-closed, EO = eyes-open.

**Table S2**

*Test-Retest Reliability of Absolute Alpha Power by Cluster and Condition*

| Measure | Cluster | Condition | ICC | *F* | *n* |
| --- | --- | --- | --- | --- | --- |
| Absolute Alpha Power | Midline | Eyes Closed | 0.90  [0.86, 0.92] | 14.95 | 175 |
|  | Perimeter | Eyes Closed | 0.90  [0.86, 0.92] | 15.16 | 175 |
|  | Midline | Eyes Open | 0.83  [0.77, 0.87] | 10.15 | 147 |
|  | Perimeter | Eyes Open | 0.84  [0.78, 0.88] | 8.32 | 147 |

Note. 95% CI are presented in brackets. All *p*-values were <.001.

### **Session 1 vs. Session 1 & 2 participants.**

As noted in the manuscript, participants with both session 1 and session 2 data (*M* = 47.05, *SD* = 13.75) were older than those with only session 1 data (*M* = 42.54, *SD* = 14.68, *t*(441.4) = 3.74, *p* <.001), and had greater peak alpha power over occipitoparietal sensors in the eyes-closed condition (both sessions *M* = 1.22, *SD* = 0.45, *t*(325) = 2.16, *p* = .031). As shown below, there were no differences in any other metrics.

**Table S3.**

*Comparison of Demographic and Spectral Parameters Between Participants Completing Session 1 Only and Participants Completing Both Sessions*

|  |  |  | **Session 1 Only** | | | **Both Sessions** | | |  |  |
| --- | --- | --- | --- | --- | --- | --- | --- | --- | --- | --- |
| **Measure** | **Cond.** | **Cl.** | ***n*** | ***M*** | ***SD*** | ***n*** | ***M*** | ***SD*** | ***t*** | ***p*** |
| Exponent | EC | 1 | 392 | 1.180 | 0.363 | 178 | 1.198 | 0.312 | −0.630 | .529 |
|  | EC | 2 | 392 | 0.911 | 0.349 | 178 | 0.934 | 0.324 | −0.772 | .440 |
|  | EO | 1 | 368 | 1.146 | 0.297 | 161 | 1.174 | 0.276 | −1.059 | .290 |
|  | EO | 2 | 368 | 0.835 | 0.283 | 161 | 0.858 | 0.276 | −0.867 | .387 |
| Offset | EC | 1 | 392 | 1.071 | 0.450 | 178 | 1.069 | 0.396 | 0.056 | .955 |
|  | EC | 2 | 392 | 0.653 | 0.427 | 178 | 0.656 | 0.387 | −0.067 | .947 |
|  | EO | 1 | 368 | 1.012 | 0.373 | 161 | 1.005 | 0.336 | 0.214 | .831 |
|  | EO | 2 | 368 | 0.612 | 0.311 | 161 | 0.585 | 0.305 | 0.920 | .358 |
| IAPF | EC | 1 | 386 | 10.157 | 0.847 | 175 | 10.102 | 0.801 | 0.734 | .464 |
|  | EC | 2 | 386 | 9.943 | 0.901 | 175 | 9.867 | 0.808 | 1.006 | .315 |
|  | EO | 1 | 323 | 10.250 | 0.906 | 147 | 10.161 | 0.867 | 1.011 | .313 |
|  | EO | 2 | 323 | 10.131 | 0.993 | 147 | 9.957 | 0.958 | 1.805 | .072 |
| Alpha power | EC | 1 | 386 | 1.135 | 0.436 | 175 | 1.223 | 0.453 | −2.161 | .031* |
|  | EC | 2 | 386 | 0.915 | 0.398 | 175 | 0.941 | 0.391 | −0.711 | .477 |
|  | EO | 1 | 323 | 0.689 | 0.359 | 147 | 0.695 | 0.344 | −0.162 | .872 |
|  | EO | 2 | 323 | 0.539 | 0.280 | 147 | 0.528 | 0.240 | 0.435 | .664 |
| Absolute Alpha power | EC | 1 | 386 | 1.340 | 0.468 | 175 | 1.386 | 0.449 | -1.089 | .277 |
|  | EC | 2 | 386 | 1.051 | 0.430 | 175 | 1.076 | 0.407 | -0.667 | .505 |
|  | EO | 1 | 323 | 0.974 | 0.383 | 147 | 0.963 | 0.345 | 0.293 | .770 |
|  | EO | 2 | 323 | 0.727 | 0.343 | 147 | 0.700 | 0.330 | 0.824 | .411 |
| Age | — | — | 400 | 42.542 | 14.683 | 207 | 47.053 | 13.754 | −3.742 | .0002* |
| Sex |  |  | 400 | 246F / 154M | | 207 | 129F / 78M | | *χ*² = 0.012 | .913 |

Note; EC = eyes closed, EO = eyes open. For the exponent, offset, and absolute alpha power, C1 = midline cluster, C2 perimeter cluster. For IAPF and alpha power, C1 = occipitoparietal cluster, C2 = frontotemporal cluster

**Cross-sectional associations with Age.**

In addition to the HLMs, we also explored cross-sectional associates between age and aperiodic activity, and age and parameterized alpha activity. We observed significant negative correlations between age and the aperiodic exponent, aperiodic offset, and IAPF. We did not observe robust associations between age and peak alpha power. Absolute alpha power was only correlated with age for eyes closed recordings, over cluster 1.

**Figure S3.**

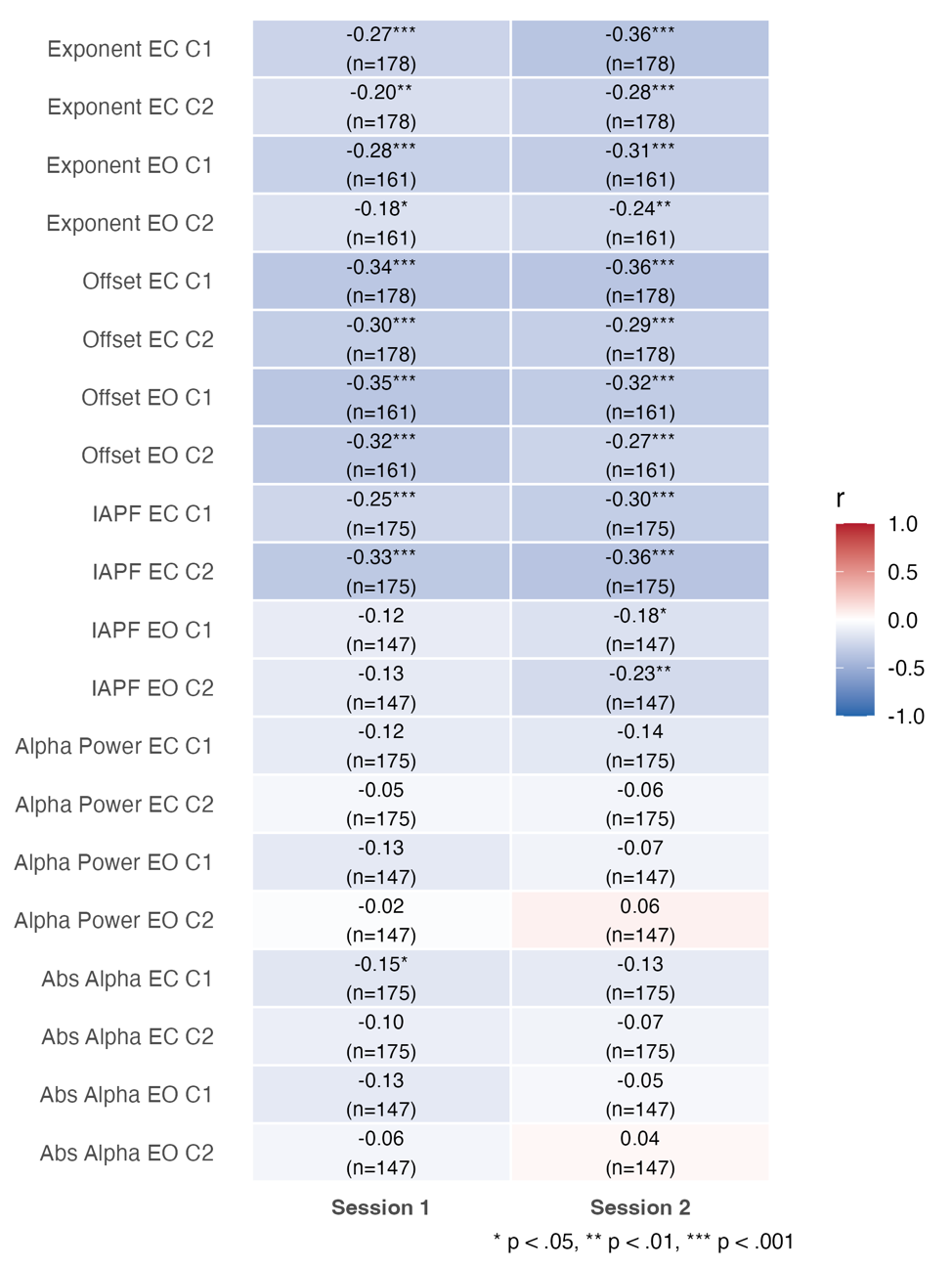
Cross-sectional associations between age and EEG measures

Note. EC = eyes closed, EO = eyes open. For the exponent, offset, and absolute alpha power, C1 = midline cluster, C2 perimeter cluster. For IAPF and alpha power, C1 = occipitoparietal cluster, C2 = frontotemporal cluster. Abs Alpha = Absolute alpha power.

**Hierarchical Linear Models**

Here, we present the outputs of the additional HLM models. These include models fitting session X age, age^2^, session X age^2^, and session X sex. In all cases, the interaction terms are non-significant, and the model fit statistics are poorer than those of the main-effect-only models. In addition, we also fitted models with absolute alpha power as the outcome variable. In these models, we conducted the same main-effect-only models, as well as session X age, age^2^, session X age^2^, and session X sex models.

**Table S4**

*Hierarchical Linear Models of Aperiodic Exponent Stability, with Session X Age interactions (Eyes-Closed)*

|  | **Midline** | | | | | | | **Perimeter** | | | | | | |
| --- | --- | --- | --- | --- | --- | --- | --- | --- | --- | --- | --- | --- | --- | --- |
| *Predictors* | *Estimates* | *std. Error* | *std. Beta* | *CI* | *Statistic* | *p* | *p*  *(FDR.adj)* | *Estimates* | *std. Error* | *std. Beta* | *CI* | *Statistic* | *p* | *p*  *(FDR.adj)* |
| Intercept | 1.286 | 0.044 | 0.080 | 1.200 – 1.372 | 29.312 | **<0.001** |  | 1.008 | 0.043 | 0.069 | 0.924 – 1.093 | 23.483 | **<0.001** |  |
| Session | -0.045 | 0.024 | -0.081 | -0.092 – 0.002 | -1.870 | 0.063 | 0.084 | -0.039 | 0.023 | -0.069 | -0.084 – 0.006 | -1.722 | 0.087 | 0.087 |
| Age | -0.003 | 0.003 | -0.301 | -0.008 – 0.003 | -0.891 | 0.374 | 0.396 | -0.002 | 0.003 | -0.229 | -0.008 – 0.003 | -0.849 | 0.396 | 0.396 |
| Sex | -0.076 | 0.043 | -0.230 | -0.161 – 0.009 | -1.774 | 0.078 | 0.295 | -0.064 | 0.044 | -0.197 | -0.152 – 0.023 | -1.455 | 0.147 | 0.295 |
| Session X Age | -0.003 | 0.002 | -0.064 | -0.006 – 0.000 | -1.808 | 0.072 | 0.289 | -0.002 | 0.002 | -0.042 | -0.005 – 0.001 | -1.235 | 0.219 | 0.437 |
| **Random Effects** | | | | | | | | | | | | | | |
| σ^2^ | 0.05 | | | | | | | 0.04 | | | | | | |
| τ_00_ | 0.05 _participant_id_ | | | | | | | 0.06 _participant_id_ | | | | | | |
| ICC | 0.50 | | | | | | | 0.56 | | | | | | |
| N | 178 _participant_id_ | | | | | | | 178 _participant_id_ | | | | | | |
| Observations | 356 | | | | | | | 356 | | | | | | |
| Marginal R^2^ / Conditional R^2^ | 0.120 / 0.560 | | | | | | | 0.072 / 0.592 | | | | | | |
| AIC | 178.239 | | | | | | | 171.800 | | | | | | |

Note: Repeated-measured EEG data nested within participant, indicated by participant_id. Sex coded as 0 = female, 1 = male.

**Table S5**

*Hierarchical Linear Models of Aperiodic Exponent Stability, with Session X Age interactions (Eyes-Open)*

|  | **Midline** | | | | | | | **Perimeter** | | | | | | |
| --- | --- | --- | --- | --- | --- | --- | --- | --- | --- | --- | --- | --- | --- | --- |
| *Predictors* | *Estimates* | *std. Error* | *std. Beta* | *CI* | *Statistic* | *p* | *p*  *(FDR.adj)* | *Estimates* | *std. Error* | *std. Beta* | *CI* | *Statistic* | *p* | *p*  *(FDR.adj)* |
| Intercept | 1.271 | 0.038 | 0.053 | 1.197 – 1.345 | 33.701 | **<0.001** |  | 0.927 | 0.037 | -0.026 | 0.855 – 0.999 | 25.293 | **<0.001** |  |
| Session | -0.067 | 0.020 | -0.122 | -0.106 – -0.029 | -3.436 | **0.001** | **0.002** | -0.066 | 0.019 | -0.125 | -0.103 – -0.029 | -3.540 | **0.001** | **0.002** |
| Age | -0.005 | 0.003 | -0.287 | -0.010 – 0.000 | -1.811 | 0.071 | 0.285 | -0.003 | 0.002 | -0.213 | -0.008 – 0.002 | -1.144 | 0.254 | 0.396 |
| Sex | -0.044 | 0.041 | -0.155 | -0.125 – 0.036 | -1.093 | 0.276 | 0.368 | 0.021 | 0.041 | 0.077 | -0.059 – 0.101 | 0.525 | 0.601 | 0.601 |
| Session X Age | -0.001 | 0.001 | -0.022 | -0.004 – 0.002 | -0.660 | 0.510 | 0.510 | -0.001 | 0.001 | -0.025 | -0.004 – 0.002 | -0.737 | 0.462 | 0.510 |
| **Random Effects** | | | | | | | | | | | | | | |
| σ^2^ | 0.03 | | | | | | | 0.03 | | | | | | |
| τ_00_ | 0.04 _participant_id_ | | | | | | | 0.05 _participant_id_ | | | | | | |
| ICC | 0.60 | | | | | | | 0.62 | | | | | | |
| N | 161 _participant_id_ | | | | | | | 161 _participant_id_ | | | | | | |
| Observations | 322 | | | | | | | 322 | | | | | | |
| Marginal R^2^ / Conditional R^2^ | 0.107 / 0.640 | | | | | | | 0.060 / 0.648 | | | | | | |
| AIC | 52.112 | | | | | | | 36.456 | | | | | | |

Note: Repeated-measured EEG data nested within participant, indicated by participant_id. Sex coded as 0 = female, 1 = male.

**Table S6**

*Hierarchical Linear Models of Aperiodic Offset Stability, with Session X Age interactions (Eyes-Closed)*

|  | **Midline** | | | | | | | **Perimeter** | | | | | | |
| --- | --- | --- | --- | --- | --- | --- | --- | --- | --- | --- | --- | --- | --- | --- |
| *Predictors* | *Estimates* | *std. Error* | *std. Beta* | *CI* | *Statistic* | *p* | *p*  *(FDR.adj)* | *Estimates* | *std. Error* | *std. Beta* | *CI* | *Statistic* | *p* | *p*  *(FDR.adj)* |
| Intercept | 1.242 | 0.051 | 0.115 | 1.141 – 1.342 | 24.316 | **<0.001** |  | 0.812 | 0.047 | 0.107 | 0.718 – 0.905 | 17.103 | **<0.001** |  |
| Session | -0.098 | 0.026 | -0.120 | -0.148 – -0.047 | -3.809 | **<0.001** | **<0.001** | -0.089 | 0.023 | -0.112 | -0.134 – -0.045 | -3.940 | **<0.001** | **<0.001** |
| Age | -0.007 | 0.003 | -0.328 | -0.014 – -0.001 | -2.160 | **0.031** | **0.031** | -0.008 | 0.003 | -0.278 | -0.014 – -0.001 | -2.437 | **0.015** | **0.020** |
| Sex | -0.142 | 0.057 | -0.329 | -0.255 – -0.029 | -2.480 | **0.014** | **0.045** | -0.125 | 0.056 | -0.307 | -0.236 – -0.014 | -2.226 | **0.027** | **0.045** |
| Session X Age | -0.002 | 0.002 | -0.032 | -0.006 – 0.002 | -1.089 | 0.277 | 0.794 | -0.000 | 0.002 | -0.007 | -0.004 – 0.003 | -0.263 | 0.793 | 0.794 |
| **Random Effects** | | | | | | | | | | | | | | |
| σ^2^ | 0.06 | | | | | | | 0.04 | | | | | | |
| τ_00_ | 0.10 _participant_id_ | | | | | | | 0.10 _participant_id_ | | | | | | |
| ICC | 0.65 | | | | | | | 0.70 | | | | | | |
| N | 178 _participant_id_ | | | | | | | 178 _participant_id_ | | | | | | |
| Observations | 356 | | | | | | | 356 | | | | | | |
| Marginal R^2^ / Conditional R^2^ | 0.158 / 0.702 | | | | | | | 0.120 / 0.737 | | | | | | |
| AIC | 302.885 | | | | | | | 254.062 | | | | | | |

Note: Repeated-measured EEG data nested within participant, indicated by participant_id. Sex coded as 0 = female, 1 = male.

**Table S7**

*Hierarchical Linear Models of Aperiodic Offset Stability, with Session X Age interactions (Eyes-Open)*

|  | **Midline** | | | | | | | **Perimeter** | | | | | | |
| --- | --- | --- | --- | --- | --- | --- | --- | --- | --- | --- | --- | --- | --- | --- |
| *Predictors* | *Estimates* | *std. Error* | *std. Beta* | *CI* | *Statistic* | *p* | *p*  *(FDR.adj)* | *Estimates* | *std. Error* | *std. Beta* | *CI* | *Statistic* | *p* | *p*  *(FDR.adj)* |
| Intercept | 1.159 | 0.046 | 0.103 | 1.069 – 1.250 | 25.142 | **<0.001** |  | 0.710 | 0.042 | 0.063 | 0.628 – 0.793 | 17.013 | **<0.001** |  |
| Session | -0.095 | 0.023 | -0.131 | -0.141 – -0.049 | -4.115 | **<0.001** | **<0.001** | -0.087 | 0.021 | -0.131 | -0.128 – -0.045 | -4.130 | **<0.001** | **<0.001** |
| Age | -0.008 | 0.003 | -0.313 | -0.014 – -0.002 | -2.524 | **0.012** | **0.020** | -0.008 | 0.003 | -0.280 | -0.013 – -0.002 | -2.700 | **0.007** | **0.020** |
| Sex | -0.112 | 0.052 | -0.302 | -0.215 – -0.009 | -2.140 | **0.034** | **0.045** | -0.061 | 0.047 | -0.186 | -0.154 – 0.032 | -1.288 | 0.199 | 0.199 |
| Session X Age | -0.000 | 0.002 | -0.008 | -0.004 – 0.003 | -0.262 | 0.794 | 0.794 | 0.001 | 0.002 | 0.012 | -0.002 – 0.004 | 0.384 | 0.702 | 0.794 |
| **Random Effects** | | | | | | | | | | | | | | |
| σ^2^ | 0.04 | | | | | | | 0.03 | | | | | | |
| τ_00_ | 0.08 _participant_id_ | | | | | | | 0.06 _participant_id_ | | | | | | |
| ICC | 0.65 | | | | | | | 0.64 | | | | | | |
| N | 161 _participant_id_ | | | | | | | 161 _participant_id_ | | | | | | |
| Observations | 322 | | | | | | | 322 | | | | | | |
| Marginal R^2^ / Conditional R^2^ | 0.144 / 0.699 | | | | | | | 0.107 / 0.682 | | | | | | |
| AIC | 184.028 | | | | | | | 120.791 | | | | | | |

Note: Repeated-measured EEG data nested within participant, indicated by participant_id. Sex coded as 0 = female, 1 = male.

**Table S8**

*Hierarchical Linear Models of Individual Alpha Peak Frequency Stability, with Session X Age interactions (Eyes-Closed)*

|  | **Occipitoparietal** | | | | | | | **Frontotemporal** | | | | | | |
| --- | --- | --- | --- | --- | --- | --- | --- | --- | --- | --- | --- | --- | --- | --- |
| *Predictors* | *Estimates* | *std. Error* | *std. Beta* | *CI* | *Statistic* | *p* | *p*  *(FDR.adj)* | *Estimates* | *std. Error* | *std. Beta* | *CI* | *Statistic* | *p* | *p*  *(FDR.adj)* |
| Intercept | 10.290 | 0.086 | 0.057 | 10.121 – 10.458 | 120.029 | **<0.001** |  | 9.997 | 0.091 | -0.001 | 9.819 – 10.176 | 110.243 | **<0.001** |  |
| Session | -0.103 | 0.034 | -0.070 | -0.169 – -0.037 | -3.082 | **0.002** | **0.010** | -0.078 | 0.040 | -0.052 | -0.157 – -0.000 | -1.975 | **0.050** | 0.100 |
| Age | -0.011 | 0.005 | -0.267 | -0.022 – -0.000 | -2.033 | **0.043** | 0.086 | -0.017 | 0.006 | -0.348 | -0.029 – -0.006 | -2.957 | **0.003** | **0.013** |
| Sex | -0.132 | 0.118 | -0.165 | -0.365 – 0.101 | -1.115 | 0.266 | 0.355 | 0.002 | 0.116 | 0.003 | -0.228 – 0.232 | 0.019 | 0.985 | 0.985 |
| Session X Age | -0.003 | 0.002 | -0.025 | -0.008 – 0.002 | -1.238 | 0.218 | 0.371 | -0.002 | 0.003 | -0.019 | -0.008 – 0.003 | -0.806 | 0.421 | 0.421 |
| **Random Effects** | | | | | | | | | | | | | | |
| σ^2^ | 0.09 | | | | | | | 0.13 | | | | | | |
| τ_00_ | 0.50 _participant_id_ | | | | | | | 0.46 _participant_id_ | | | | | | |
| ICC | 0.84 | | | | | | | 0.78 | | | | | | |
| N | 175 _participant_id_ | | | | | | | 175 _participant_id_ | | | | | | |
| Observations | 350 | | | | | | | 350 | | | | | | |
| Marginal R^2^ / Conditional R^2^ | 0.087 / 0.854 | | | | | | | 0.123 / 0.805 | | | | | | |
| AIC | 635.609 | | | | | | | 688.892 | | | | | | |

Note: Repeated-measured EEG data nested within participant, indicated by participant_id. Sex coded as 0 = female, 1 = male.

**Table S9**

*Hierarchical Linear Models of Individual Alpha Peak Frequency Stability, with Session X Age interactions (Eyes-Open)*

|  | **Occipitoparietal** | | | | | | | **Frontotemporal** | | | | | | |
| --- | --- | --- | --- | --- | --- | --- | --- | --- | --- | --- | --- | --- | --- | --- |
| *Predictors* | *Estimates* | *std. Error* | *std. Beta* | *CI* | *Statistic* | *p* | *p*  *(FDR.adj)* | *Estimates* | *std. Error* | *std. Beta* | *CI* | *Statistic* | *p* | *p*  *(FDR.adj)* |
| Intercept | 10.314 | 0.114 | 0.102 | 10.090 – 10.538 | 90.748 | **<0.001** |  | 10.060 | 0.134 | 0.070 | 9.797 – 10.323 | 75.293 | **<0.001** |  |
| Session | -0.046 | 0.053 | -0.033 | -0.150 – 0.058 | -0.875 | 0.383 | 0.510 | -0.014 | 0.068 | -0.017 | -0.148 – 0.119 | -0.212 | 0.833 | 0.833 |
| Age | -0.003 | 0.007 | -0.137 | -0.017 – 0.012 | -0.346 | 0.729 | 0.848 | -0.002 | 0.009 | -0.169 | -0.019 – 0.016 | -0.192 | 0.848 | 0.848 |
| Sex | -0.262 | 0.140 | -0.300 | -0.540 – 0.015 | -1.871 | 0.063 | 0.254 | -0.196 | 0.149 | -0.206 | -0.491 – 0.098 | -1.320 | 0.189 | 0.355 |
| Session X Age | -0.004 | 0.004 | -0.032 | -0.011 – 0.003 | -1.089 | 0.278 | 0.371 | -0.007 | 0.005 | -0.048 | -0.016 – 0.003 | -1.381 | 0.169 | 0.371 |
| **Random Effects** | | | | | | | | | | | | | | |
| σ^2^ | 0.20 | | | | | | | 0.32 | | | | | | |
| τ_00_ | 0.55 _participant_id_ | | | | | | | 0.56 _participant_id_ | | | | | | |
| ICC | 0.74 | | | | | | | 0.63 | | | | | | |
| N | 147 _participant_id_ | | | | | | | 147 _participant_id_ | | | | | | |
| Observations | 294 | | | | | | | 294 | | | | | | |
| Marginal R^2^ / Conditional R^2^ | 0.044 / 0.748 | | | | | | | 0.043 / 0.650 | | | | | | |
| AIC | 668.381 | | | | | | | 758.872 | | | | | | |

Note: Repeated-measured EEG data nested within participant, indicated by participant_id. Sex coded as 0 = female, 1 = male.

**Table S10**

*Hierarchical Linear Models of Alpha Power Stability, with Session X Age interactions (Eyes-Closed)*

|  | **Occipitoparietal** | | | | | | | **Frontotemporal** | | | | | | |
| --- | --- | --- | --- | --- | --- | --- | --- | --- | --- | --- | --- | --- | --- | --- |
| *Predictors* | *Estimates* | *std. Error* | *std. Beta* | *CI* | *Statistic* | *p* | *p*  *(FDR.adj)* | *Estimates* | *std. Error* | *std. Beta* | *CI* | *Statistic* | *p* | *p*  *(FDR.adj)* |
| Intercept | 1.242 | 0.049 | 0.050 | 1.146 – 1.337 | 25.497 | **<0.001** |  | 0.929 | 0.042 | 0.053 | 0.847 – 1.012 | 22.206 | **<0.001** |  |
| Session | 0.015 | 0.018 | 0.014 | -0.020 – 0.049 | 0.831 | 0.407 | 0.543 | 0.035 | 0.015 | 0.044 | 0.006 – 0.065 | 2.354 | **0.020** | 0.079 |
| Age | -0.003 | 0.003 | -0.122 | -0.009 – 0.003 | -0.960 | 0.338 | 0.519 | -0.001 | 0.003 | -0.048 | -0.006 – 0.004 | -0.330 | 0.741 | 0.741 |
| Sex | -0.065 | 0.070 | -0.144 | -0.203 – 0.072 | -0.937 | 0.350 | 0.467 | -0.059 | 0.060 | -0.152 | -0.178 – 0.060 | -0.979 | 0.329 | 0.467 |
| Session X Age | -0.001 | 0.001 | -0.011 | -0.003 – 0.002 | -0.570 | 0.569 | 0.759 | -0.000 | 0.001 | -0.006 | -0.002 – 0.002 | -0.295 | 0.768 | 0.768 |
| **Random Effects** | | | | | | | | | | | | | | |
| σ^2^ | 0.03 | | | | | | | 0.02 | | | | | | |
| τ_00_ | 0.18 _participant_id_ | | | | | | | 0.13 _participant_id_ | | | | | | |
| ICC | 0.87 | | | | | | | 0.88 | | | | | | |
| N | 175 _participant_id_ | | | | | | | 175 _participant_id_ | | | | | | |
| Observations | 350 | | | | | | | 350 | | | | | | |
| Marginal R^2^ / Conditional R^2^ | 0.022 / 0.876 | | | | | | | 0.010 / 0.877 | | | | | | |
| AIC | 229.962 | | | | | | | 123.304 | | | | | | |

Note: Repeated-measured EEG data nested within participant, indicated by participant_id. Sex coded as 0 = female, 1 = male.

**Table S11**

*Hierarchical Linear Models of Alpha Power Stability, with Session X Age interactions (Eyes-Open)*

|  | **Occipitoparietal** | | | | | | | **Frontotemporal** | | | | | | |
| --- | --- | --- | --- | --- | --- | --- | --- | --- | --- | --- | --- | --- | --- | --- |
| *Predictors* | *Estimates* | *std. Error* | *std. Beta* | *CI* | *Statistic* | *p* | *p*  *(FDR.adj)* | *Estimates* | *std. Error* | *std. Beta* | *CI* | *Statistic* | *p* | *p*  *(FDR.adj)* |
| Intercept | 0.732 | 0.041 | 0.029 | 0.651 – 0.813 | 17.783 | **<0.001** |  | 0.536 | 0.031 | 0.062 | 0.476 – 0.596 | 17.575 | **<0.001** |  |
| Session | -0.019 | 0.017 | -0.022 | -0.052 – 0.014 | -1.136 | 0.258 | 0.515 | 0.007 | 0.013 | 0.023 | -0.019 – 0.033 | 0.551 | 0.583 | 0.583 |
| Age | -0.005 | 0.003 | -0.100 | -0.010 – 0.000 | -1.843 | 0.066 | 0.266 | -0.002 | 0.002 | 0.030 | -0.006 – 0.002 | -0.862 | 0.390 | 0.519 |
| Sex | -0.028 | 0.056 | -0.085 | -0.138 – 0.082 | -0.507 | 0.613 | 0.613 | -0.044 | 0.040 | -0.183 | -0.123 – 0.035 | -1.105 | 0.271 | 0.467 |
| Session X Age | 0.002 | 0.001 | 0.033 | -0.001 – 0.004 | 1.350 | 0.179 | 0.358 | 0.001 | 0.001 | 0.042 | -0.000 – 0.003 | 1.573 | 0.118 | 0.358 |
| **Random Effects** | | | | | | | | | | | | | | |
| σ^2^ | 0.02 | | | | | | | 0.01 | | | | | | |
| τ_00_ | 0.09 _participant_id_ | | | | | | | 0.05 _participant_id_ | | | | | | |
| ICC | 0.82 | | | | | | | 0.79 | | | | | | |
| N | 147 _participant_id_ | | | | | | | 147 _participant_id_ | | | | | | |
| Observations | 294 | | | | | | | 294 | | | | | | |
| Marginal R^2^ / Conditional R^2^ | 0.014 / 0.824 | | | | | | | 0.010 / 0.790 | | | | | | |
| AIC | 71.812 | | | | | | | -95.652 | | | | | | |

Note: Repeated-measured EEG data nested within participant, indicated by participant_id. Sex coded as 0 = female, 1 = male.

**Table S12**

*Hierarchical Linear Models of Aperiodic Exponent Stability (Eyes-Closed) with Age^2^*

|  | **Midline** | | | | | | | **Perimeter** | | | | | | |
| --- | --- | --- | --- | --- | --- | --- | --- | --- | --- | --- | --- | --- | --- | --- |
| *Predictors* | *Estimates* | *std. Error* | *std. Beta* | *CI* | *Statistic* | *p* | *p*  *(FDR.adj)* | *Estimates* | *std. Error* | *std. Beta* | *CI* | *Statistic* | *p* | *p*  *(FDR.adj)* |
| Intercept | 1.346 | 0.049 | 0.228 | 1.249 – 1.443 | 27.349 | **<0.001** |  | 1.042 | 0.048 | 0.180 | 0.947 – 1.137 | 21.573 | **<0.001** |  |
| Session | -0.053 | 0.024 | -0.081 | -0.100 – -0.007 | -2.267 | **0.025** | **0.033** | -0.045 | 0.022 | -0.069 | -0.089 – -0.001 | -2.009 | **0.046** | **0.046** |
| Age² | -0.000 | 0.000 | -0.127 | -0.001 – -0.000 | -2.679 | **0.008** | **0.032** | -0.000 | 0.000 | -0.094 | -0.000 – 0.000 | -1.509 | 0.133 | 0.177 |
| Sex | -0.094 | 0.045 | -0.292 | -0.182 – -0.006 | -2.099 | **0.037** | 0.149 | -0.079 | 0.045 | -0.245 | -0.168 – 0.011 | -1.741 | 0.083 | 0.154 |
| **Random Effects** | | | | | | | | | | | | | | |
| σ^2^ | 0.05 | | | | | | | 0.04 | | | | | | |
| τ_00_ | 0.06 _participant_id_ | | | | | | | 0.06 _participant_id_ | | | | | | |
| ICC | 0.53 | | | | | | | 0.58 | | | | | | |
| N | 178 _participant_id_ | | | | | | | 178 _participant_id_ | | | | | | |
| Observations | 356 | | | | | | | 356 | | | | | | |
| Marginal R^2^ / Conditional R^2^ | 0.057 / 0.555 | | | | | | | 0.029 / 0.590 | | | | | | |
| AIC | 189.040 | | | | | | | 175.341 | | | | | | |

Note: Repeated-measured EEG data nested within participant, indicated by participant_id. Sex coded as 0 = female, 1 = male.

**Table S13**

*Hierarchical Linear Models of Aperiodic Exponent Stability (Eyes-Open) with Age^2^*

|  | **Midline** | | | | | | | **Perimeter** | | | | | | |
| --- | --- | --- | --- | --- | --- | --- | --- | --- | --- | --- | --- | --- | --- | --- |
| *Predictors* | *Estimates* | *std. Error* | *std. Beta* | *CI* | *Statistic* | *p* | *p*  *(FDR.adj)* | *Estimates* | *std. Error* | *std. Beta* | *CI* | *Statistic* | *p* | *p*  *(FDR.adj)* |
| Intercept | 1.305 | 0.044 | 0.178 | 1.218 – 1.391 | 29.616 | **<0.001** |  | 0.949 | 0.043 | 0.071 | 0.865 – 1.033 | 22.255 | **<0.001** |  |
| Session | -0.070 | 0.019 | -0.122 | -0.108 – -0.032 | -3.629 | **<0.001** | **0.001** | -0.069 | 0.018 | -0.125 | -0.105 – -0.033 | -3.749 | **<0.001** | **0.001** |
| Age² | -0.000 | 0.000 | -0.099 | -0.000 – 0.000 | -1.726 | 0.086 | 0.173 | -0.000 | 0.000 | -0.078 | -0.000 – 0.000 | -1.123 | 0.263 | 0.263 |
| Sex | -0.067 | 0.042 | -0.231 | -0.150 – 0.017 | -1.581 | 0.116 | 0.154 | 0.005 | 0.041 | 0.021 | -0.076 – 0.087 | 0.131 | 0.896 | 0.896 |
| **Random Effects** | | | | | | | | | | | | | | |
| σ^2^ | 0.03 | | | | | | | 0.03 | | | | | | |
| τ_00_ | 0.05 _participant_id_ | | | | | | | 0.05 _participant_id_ | | | | | | |
| ICC | 0.63 | | | | | | | 0.64 | | | | | | |
| N | 161 _participant_id_ | | | | | | | 161 _participant_id_ | | | | | | |
| Observations | 322 | | | | | | | 322 | | | | | | |
| Marginal R^2^ / Conditional R^2^ | 0.040 / 0.641 | | | | | | | 0.022 / 0.648 | | | | | | |
| AIC | 58.527 | | | | | | | 36.548 | | | | | | |

Note: Repeated-measured EEG data nested within participant, indicated by participant_id. Sex coded as 0 = female, 1 = male.

**Table S14**

*Hierarchical Linear Models of Aperiodic Exponent Stability, with Session X Age^2^ interactions (Eyes-Closed)*

|  | **Midline** | | | | | | | **Perimeter** | | | | | | |
| --- | --- | --- | --- | --- | --- | --- | --- | --- | --- | --- | --- | --- | --- | --- |
| *Predictors* | *Estimates* | *std. Error* | *std. Beta* | *CI* | *Statistic* | *p* | *p*  *(FDR.adj)* | *Estimates* | *std. Error* | *std. Beta* | *CI* | *Statistic* | *p* | *p*  *(FDR.adj)* |
| Intercept | 1.302 | 0.062 | 0.228 | 1.179 – 1.424 | 20.903 | **<0.001** |  | 1.015 | 0.060 | 0.180 | 0.896 – 1.133 | 16.802 | **<0.001** |  |
| Session | -0.024 | 0.035 | -0.064 | -0.092 – 0.044 | -0.692 | 0.490 | 0.490 | -0.027 | 0.033 | -0.066 | -0.092 – 0.038 | -0.813 | 0.417 | 0.490 |
| Age | -0.000 | 0.000 | -0.127 | -0.001 – 0.000 | -0.390 | 0.697 | 0.933 | -0.000 | 0.000 | -0.094 | -0.000 – 0.000 | -0.190 | 0.850 | 0.933 |
| Sex | -0.094 | 0.045 | -0.292 | -0.182 – -0.006 | -2.099 | **0.037** | 0.149 | -0.079 | 0.045 | -0.245 | -0.168 – 0.011 | -1.741 | 0.083 | 0.154 |
| Session X Age² | -0.000 | 0.000 | -0.017 | -0.000 – 0.000 | -1.156 | 0.249 | 0.487 | -0.000 | 0.000 | -0.002 | -0.000 – 0.000 | -0.748 | 0.455 | 0.487 |
| **Random Effects** | | | | | | | | | | | | | | |
| σ^2^ | 0.05 | | | | | | | 0.04 | | | | | | |
| τ_00_ | 0.06 _participant_id_ | | | | | | | 0.06 _participant_id_ | | | | | | |
| ICC | 0.53 | | | | | | | 0.58 | | | | | | |
| N | 178 _participant_id_ | | | | | | | 178 _participant_id_ | | | | | | |
| Observations | 356 | | | | | | | 356 | | | | | | |
| Marginal R^2^ / Conditional R^2^ | 0.058 / 0.556 | | | | | | | 0.030 / 0.590 | | | | | | |
| AIC | 205.785 | | | | | | | 192.970 | | | | | | |

Note: Repeated-measured EEG data nested within participant, indicated by participant_id. Sex coded as 0 = female, 1 = male.

**Table S15**

*Hierarchical Linear Models of Aperiodic Exponent Stability, with Session X Age^2^ interactions (Eyes-Open)*

|  | **Midline** | | | | | | | **Perimeter** | | | | | | |
| --- | --- | --- | --- | --- | --- | --- | --- | --- | --- | --- | --- | --- | --- | --- |
| *Predictors* | *Estimates* | *std. Error* | *std. Beta* | *CI* | *Statistic* | *p* | *p*  *(FDR.adj)* | *Estimates* | *std. Error* | *std. Beta* | *CI* | *Statistic* | *p* | *p*  *(FDR.adj)* |
| Intercept | 1.281 | 0.054 | 0.178 | 1.174 – 1.387 | 23.647 | **<0.001** |  | 0.928 | 0.052 | 0.071 | 0.825 – 1.031 | 17.782 | **<0.001** |  |
| Session | -0.054 | 0.028 | -0.106 | -0.110 – 0.002 | -1.892 | 0.060 | 0.121 | -0.055 | 0.027 | -0.109 | -0.108 – -0.001 | -2.014 | **0.046** | 0.121 |
| Age | -0.000 | 0.000 | -0.099 | -0.000 – 0.000 | -0.366 | 0.715 | 0.933 | -0.000 | 0.000 | -0.078 | -0.000 – 0.000 | -0.084 | 0.933 | 0.933 |
| Sex | -0.067 | 0.042 | -0.231 | -0.150 – 0.017 | -1.581 | 0.116 | 0.154 | 0.005 | 0.041 | 0.021 | -0.076 – 0.087 | 0.131 | 0.896 | 0.896 |
| Session X Age² | -0.000 | 0.000 | -0.017 | -0.000 – 0.000 | -0.754 | 0.452 | 0.487 | -0.000 | 0.000 | -0.015 | -0.000 – 0.000 | -0.697 | 0.487 | 0.487 |
| **Random Effects** | | | | | | | | | | | | | | |
| σ^2^ | 0.03 | | | | | | | 0.03 | | | | | | |
| τ_00_ | 0.05 _participant_id_ | | | | | | | 0.05 _participant_id_ | | | | | | |
| ICC | 0.63 | | | | | | | 0.64 | | | | | | |
| N | 161 _participant_id_ | | | | | | | 161 _participant_id_ | | | | | | |
| Observations | 322 | | | | | | | 322 | | | | | | |
| Marginal R^2^ / Conditional R^2^ | 0.041 / 0.641 | | | | | | | 0.022 / 0.648 | | | | | | |
| AIC | 76.371 | | | | | | | 54.567 | | | | | | |

Note: Repeated-measured EEG data nested within participant, indicated by participant_id. Sex coded as 0 = female, 1 = male.

**Table S16**

*Hierarchical Linear Models of Aperiodic Offset Stability (Eyes-Closed) with Age^2^*

|  | **Midline** | | | | | | | **Perimeter** | | | | | | |
| --- | --- | --- | --- | --- | --- | --- | --- | --- | --- | --- | --- | --- | --- | --- |
| *Predictors* | *Estimates* | *std. Error* | *std. Beta* | *CI* | *Statistic* | *p* | *p*  *(FDR.adj)* | *Estimates* | *std. Error* | *std. Beta* | *CI* | *Statistic* | *p* | *p*  *(FDR.adj)* |
| Intercept | 1.274 | 0.060 | 0.237 | 1.156 – 1.392 | 21.247 | **<0.001** |  | 0.811 | 0.056 | 0.201 | 0.701 – 0.921 | 14.499 | **<0.001** |  |
| Session | -0.103 | 0.025 | -0.120 | -0.153 – -0.054 | -4.111 | **<0.001** | **<0.001** | -0.091 | 0.022 | -0.112 | -0.134 – -0.047 | -4.088 | **<0.001** | **<0.001** |
| Age | -0.000 | 0.000 | -0.098 | -0.001 – 0.000 | -1.347 | 0.180 | 0.719 | -0.000 | 0.000 | -0.072 | -0.000 – 0.000 | -0.391 | 0.696 | 0.928 |
| Sex | -0.172 | 0.061 | -0.401 | -0.292 – -0.052 | -2.833 | **0.005** | **0.016** | -0.152 | 0.059 | -0.370 | -0.268 – -0.036 | -2.595 | **0.010** | **0.016** |
| **Random Effects** | | | | | | | | | | | | | | |
| σ^2^ | 0.06 | | | | | | | 0.04 | | | | | | |
| τ_00_ | 0.12 _participant_id_ | | | | | | | 0.12 _participant_id_ | | | | | | |
| ICC | 0.68 | | | | | | | 0.73 | | | | | | |
| N | 178 _participant_id_ | | | | | | | 178 _participant_id_ | | | | | | |
| Observations | 356 | | | | | | | 356 | | | | | | |
| Marginal R^2^ / Conditional R^2^ | 0.060 / 0.702 | | | | | | | 0.045 / 0.739 | | | | | | |
| AIC | 319.729 | | | | | | | 263.126 | | | | | | |

Note: Repeated-measured EEG data nested within participant, indicated by participant_id. Sex coded as 0 = female, 1 = male.

**Table S17**

*Hierarchical Linear Models of Aperiodic Offset Stability (Eyes-Open) with Age^2^*

|  | **Midline** | | | | | | | **Perimeter** | | | | | | |
| --- | --- | --- | --- | --- | --- | --- | --- | --- | --- | --- | --- | --- | --- | --- |
| *Predictors* | *Estimates* | *std. Error* | *std. Beta* | *CI* | *Statistic* | *p* | *p*  *(FDR.adj)* | *Estimates* | *std. Error* | *std. Beta* | *CI* | *Statistic* | *p* | *p*  *(FDR.adj)* |
| Intercept | 1.165 | 0.055 | 0.196 | 1.056 – 1.274 | 21.076 | **<0.001** |  | 0.698 | 0.050 | 0.137 | 0.600 – 0.796 | 14.077 | **<0.001** |  |
| Session | -0.096 | 0.023 | -0.131 | -0.141 – -0.052 | -4.251 | **<0.001** | **<0.001** | -0.085 | 0.021 | -0.131 | -0.126 – -0.045 | -4.142 | **<0.001** | **<0.001** |
| Age | -0.000 | 0.000 | -0.066 | -0.000 – 0.000 | -0.556 | 0.579 | 0.928 | 0.000 | 0.000 | -0.050 | -0.000 – 0.000 | 0.009 | 0.993 | 0.993 |
| Sex | -0.140 | 0.055 | -0.382 | -0.249 – -0.031 | -2.540 | **0.012** | **0.016** | -0.082 | 0.049 | -0.256 | -0.180 – 0.015 | -1.675 | 0.096 | 0.096 |
| **Random Effects** | | | | | | | | | | | | | | |
| σ^2^ | 0.04 | | | | | | | 0.03 | | | | | | |
| τ_00_ | 0.09 _participant_id_ | | | | | | | 0.07 _participant_id_ | | | | | | |
| ICC | 0.68 | | | | | | | 0.67 | | | | | | |
| N | 161 _participant_id_ | | | | | | | 161 _participant_id_ | | | | | | |
| Observations | 322 | | | | | | | 322 | | | | | | |
| Marginal R^2^ / Conditional R^2^ | 0.050 / 0.701 | | | | | | | 0.031 / 0.684 | | | | | | |
| AIC | 196.284 | | | | | | | 128.768 | | | | | | |

Note: Repeated-measured EEG data nested within participant, indicated by participant_id. Sex coded as 0 = female, 1 = male.

**Table S18**

*Hierarchical Linear Models of Aperiodic Offset Stability, with Session X Age^2^ interactions (Eyes-Closed)*

|  | **Midline** | | | | | | | **Perimeter** | | | | | | |
| --- | --- | --- | --- | --- | --- | --- | --- | --- | --- | --- | --- | --- | --- | --- |
| *Predictors* | *Estimates* | *std. Error* | *std. Beta* | *CI* | *Statistic* | *p* | *p*  *(FDR.adj)* | *Estimates* | *std. Error* | *std. Beta* | *CI* | *Statistic* | *p* | *p*  *(FDR.adj)* |
| Intercept | 1.206 | 0.072 | 0.237 | 1.064 – 1.348 | 16.687 | **<0.001** |  | 0.769 | 0.066 | 0.201 | 0.638 – 0.900 | 11.567 | **<0.001** |  |
| Session | -0.058 | 0.037 | -0.106 | -0.130 – 0.015 | -1.570 | 0.118 | 0.118 | -0.062 | 0.033 | -0.111 | -0.127 – 0.002 | -1.914 | 0.057 | 0.076 |
| Age² | 0.000 | 0.000 | -0.098 | -0.000 – 0.001 | 0.508 | 0.612 | 0.816 | 0.000 | 0.000 | -0.072 | -0.000 – 0.001 | 0.650 | 0.516 | 0.816 |
| Sex | -0.172 | 0.061 | -0.401 | -0.292 – -0.052 | -2.833 | **0.005** | **0.016** | -0.152 | 0.059 | -0.370 | -0.268 – -0.036 | -2.595 | **0.010** | **0.016** |
| Session X Age² | -0.000 | 0.000 | -0.014 | -0.000 – 0.000 | -1.683 | 0.094 | 0.377 | -0.000 | 0.000 | -0.001 | -0.000 – 0.000 | -1.177 | 0.241 | 0.429 |
| **Random Effects** | | | | | | | | | | | | | | |
| σ^2^ | 0.06 | | | | | | | 0.04 | | | | | | |
| τ_00_ | 0.12 _participant_id_ | | | | | | | 0.12 _participant_id_ | | | | | | |
| ICC | 0.69 | | | | | | | 0.73 | | | | | | |
| N | 178 _participant_id_ | | | | | | | 178 _participant_id_ | | | | | | |
| Observations | 356 | | | | | | | 356 | | | | | | |
| Marginal R^2^ / Conditional R^2^ | 0.063 / 0.705 | | | | | | | 0.046 / 0.740 | | | | | | |
| AIC | 334.871 | | | | | | | 279.944 | | | | | | |

Note: Repeated-measured EEG data nested within participant, indicated by participant_id. Sex coded as 0 = female, 1 = male.

**Table S19**

*Hierarchical Linear Models of Aperiodic Offset Stability, with Session X Age^2^ interactions (Eyes-Open)*

|  | **Midline** | | | | | | | **Perimeter** | | | | | | |
| --- | --- | --- | --- | --- | --- | --- | --- | --- | --- | --- | --- | --- | --- | --- |
| *Predictors* | *Estimates* | *std. Error* | *std. Beta* | *CI* | *Statistic* | *p* | *p*  *(FDR.adj)* | *Estimates* | *std. Error* | *std. Beta* | *CI* | *Statistic* | *p* | *p*  *(FDR.adj)* |
| Intercept | 1.139 | 0.067 | 0.196 | 1.008 – 1.270 | 17.091 | **<0.001** |  | 0.665 | 0.060 | 0.137 | 0.547 – 0.783 | 11.085 | **<0.001** |  |
| Session | -0.079 | 0.034 | -0.123 | -0.145 – -0.012 | -2.346 | **0.020** | 0.076 | -0.063 | 0.030 | -0.113 | -0.123 – -0.003 | -2.064 | **0.041** | 0.076 |
| Age² | 0.000 | 0.000 | -0.066 | -0.000 – 0.001 | 0.220 | 0.826 | 0.826 | 0.000 | 0.000 | -0.050 | -0.000 – 0.001 | 0.799 | 0.425 | 0.816 |
| Sex | -0.140 | 0.055 | -0.382 | -0.249 – -0.031 | -2.540 | **0.012** | **0.016** | -0.082 | 0.049 | -0.256 | -0.180 – 0.015 | -1.675 | 0.096 | 0.096 |
| Session X Age² | -0.000 | 0.000 | -0.008 | -0.000 – 0.000 | -0.706 | 0.481 | 0.481 | -0.000 | 0.000 | -0.018 | -0.000 – 0.000 | -0.994 | 0.322 | 0.429 |
| **Random Effects** | | | | | | | | | | | | | | |
| σ^2^ | 0.04 | | | | | | | 0.03 | | | | | | |
| τ_00_ | 0.09 _participant_id_ | | | | | | | 0.07 _participant_id_ | | | | | | |
| ICC | 0.68 | | | | | | | 0.67 | | | | | | |
| N | 161 _participant_id_ | | | | | | | 161 _participant_id_ | | | | | | |
| Observations | 322 | | | | | | | 322 | | | | | | |
| Marginal R^2^ / Conditional R^2^ | 0.050 / 0.700 | | | | | | | 0.032 / 0.684 | | | | | | |
| AIC | 213.867 | | | | | | | 146.055 | | | | | | |

Note: Repeated-measured EEG data nested within participant, indicated by participant_id. Sex coded as 0 = female, 1 = male.

**Table S20**

*Hierarchical Linear Models of Individual Alpha Peak Frequency Stability (Eyes-Closed) with Age^2^*

|  | **Occipitoparietal** | | | | | | | **Frontotemporal** | | | | | | |
| --- | --- | --- | --- | --- | --- | --- | --- | --- | --- | --- | --- | --- | --- | --- |
| *Predictors* | *Estimates* | *std. Error* | *std. Beta* | *CI* | *Statistic* | *p* | *p*  *(FDR.adj)* | *Estimates* | *std. Error* | *std. Beta* | *CI* | *Statistic* | *p* | *p*  *(FDR.adj)* |
| Intercept | 10.380 | 0.106 | 0.189 | 10.172 – 10.588 | 98.379 | **<0.001** |  | 10.063 | 0.112 | 0.160 | 9.843 – 10.282 | 90.233 | **<0.001** |  |
| Session | -0.111 | 0.033 | -0.070 | -0.176 – -0.046 | -3.381 | **0.001** | **0.004** | -0.085 | 0.039 | -0.052 | -0.161 – -0.008 | -2.175 | **0.031** | 0.062 |
| Age | -0.001 | 0.000 | -0.111 | -0.001 – 0.000 | -1.682 | 0.094 | 0.320 | -0.000 | 0.000 | -0.133 | -0.001 – 0.000 | -1.411 | 0.160 | 0.320 |
| Sex | -0.174 | 0.121 | -0.225 | -0.413 – 0.066 | -1.433 | 0.154 | 0.205 | -0.061 | 0.124 | -0.077 | -0.305 – 0.183 | -0.492 | 0.623 | 0.623 |
| **Random Effects** | | | | | | | | | | | | | | |
| σ^2^ | 0.09 | | | | | | | 0.13 | | | | | | |
| τ_00_ | 0.53 _participant_id_ | | | | | | | 0.54 _participant_id_ | | | | | | |
| ICC | 0.85 | | | | | | | 0.80 | | | | | | |
| N | 175 _participant_id_ | | | | | | | 175 _participant_id_ | | | | | | |
| Observations | 350 | | | | | | | 350 | | | | | | |
| Marginal R^2^ / Conditional R^2^ | 0.032 / 0.854 | | | | | | | 0.015 / 0.806 | | | | | | |
| AIC | 641.038 | | | | | | | 705.381 | | | | | | |

Note: Repeated-measured EEG data nested within participant, indicated by participant_id. Sex coded as 0 = female, 1 = male.

**Table S21**

*Hierarchical Linear Models of Individual Alpha Peak Frequency Stability (Eyes-Open) with Age^2^*

|  | **Occipitoparietal** | | | | | | | **Frontotemporal** | | | | | | |
| --- | --- | --- | --- | --- | --- | --- | --- | --- | --- | --- | --- | --- | --- | --- |
| *Predictors* | *Estimates* | *std. Error* | *std. Beta* | *CI* | *Statistic* | *p* | *p*  *(FDR.adj)* | *Estimates* | *std. Error* | *std. Beta* | *CI* | *Statistic* | *p* | *p*  *(FDR.adj)* |
| Intercept | 10.369 | 0.136 | 0.194 | 10.101 – 10.638 | 75.998 | **<0.001** |  | 10.070 | 0.156 | 0.140 | 9.762 – 10.378 | 64.407 | **<0.001** |  |
| Session | -0.057 | 0.052 | -0.033 | -0.159 – 0.045 | -1.103 | 0.272 | 0.362 | -0.032 | 0.067 | -0.017 | -0.164 – 0.099 | -0.483 | 0.630 | 0.630 |
| Age | -0.000 | 0.000 | -0.079 | -0.001 – 0.000 | -0.710 | 0.479 | 0.638 | -0.000 | 0.000 | -0.057 | -0.001 – 0.001 | -0.045 | 0.964 | 0.964 |
| Sex | -0.289 | 0.141 | -0.339 | -0.568 – -0.010 | -2.050 | **0.042** | 0.169 | -0.227 | 0.151 | -0.246 | -0.526 – 0.071 | -1.504 | 0.135 | 0.205 |
| **Random Effects** | | | | | | | | | | | | | | |
| σ^2^ | 0.20 | | | | | | | 0.33 | | | | | | |
| τ_00_ | 0.56 _participant_id_ | | | | | | | 0.59 _participant_id_ | | | | | | |
| ICC | 0.74 | | | | | | | 0.64 | | | | | | |
| N | 147 _participant_id_ | | | | | | | 147 _participant_id_ | | | | | | |
| Observations | 294 | | | | | | | 294 | | | | | | |
| Marginal R^2^ / Conditional R^2^ | 0.028 / 0.748 | | | | | | | 0.013 / 0.648 | | | | | | |
| AIC | 666.033 | | | | | | | 760.170 | | | | | | |

Note: Repeated-measured EEG data nested within participant, indicated by participant_id. Sex coded as 0 = female, 1 = male.

**Table S22**

*Hierarchical Linear Models of Individual Alpha Peak Frequency Stability, with Session X Age^2^ interactions (Eyes-Closed)*

|  | **Occipitoparietal** | | | | | | | **Frontotemporal** | | | | | | |
| --- | --- | --- | --- | --- | --- | --- | --- | --- | --- | --- | --- | --- | --- | --- |
| *Predictors* | *Estimates* | *std. Error* | *std. Beta* | *CI* | *Statistic* | *p* | *p*  *(FDR.adj)* | *Estimates* | *std. Error* | *std. Beta* | *CI* | *Statistic* | *p* | *p*  *(FDR.adj)* |
| Intercept | 10.366 | 0.119 | 0.189 | 10.133 – 10.599 | 87.461 | **<0.001** |  | 10.152 | 0.128 | 0.160 | 9.900 – 10.404 | 79.238 | **<0.001** |  |
| Session | -0.102 | 0.049 | -0.073 | -0.198 – -0.006 | -2.094 | **0.038** | 0.051 | -0.144 | 0.057 | -0.075 | -0.257 – -0.031 | -2.514 | **0.013** | 0.075 |
| Age² | -0.000 | 0.000 | -0.111 | -0.001 – 0.000 | -1.110 | 0.268 | 0.187 | -0.001 | 0.000 | -0.133 | -0.002 – -0.000 | -1.995 | **0.047** | 0.357 |
| Sex | -0.174 | 0.121 | -0.225 | -0.413 – 0.066 | -1.433 | 0.154 | 0.623 | -0.061 | 0.124 | -0.077 | -0.305 – 0.183 | -0.492 | 0.623 | 0.205 |
| Session X Age² | -0.000 | 0.000 | 0.004 | -0.000 – 0.000 | -0.256 | 0.798 | 0.387 | 0.000 | 0.000 | 0.024 | -0.000 – 0.001 | 1.410 | 0.160 | 0.873 |
| **Random Effects** | | | | | | | | | | | | | | |
| σ^2^ | 0.10 | | | | | | | 0.13 | | | | | | |
| τ_00_ | 0.53 _participant_id_ | | | | | | | 0.54 _participant_id_ | | | | | | |
| ICC | 0.85 | | | | | | | 0.80 | | | | | | |
| N | 175 _participant_id_ | | | | | | | 175 _participant_id_ | | | | | | |
| Observations | 350 | | | | | | | 350 | | | | | | |
| Marginal R^2^ / Conditional R^2^ | 0.032 / 0.853 | | | | | | | 0.016 / 0.807 | | | | | | |
| AIC | 658.389 | | | | | | | 720.491 | | | | | | |

Note: Repeated-measured EEG data nested within participant, indicated by participant_id. Sex coded as 0 = female, 1 = male.

**Table S23**

*Hierarchical Linear Models of Individual Alpha Peak Frequency Stability, with Session X Age^2^ interactions (Eyes-Open)*

|  | **Occipitoparietal** | | | | | | | **Frontotemporal** | | | | | | |
| --- | --- | --- | --- | --- | --- | --- | --- | --- | --- | --- | --- | --- | --- | --- |
| *Predictors* | *Estimates* | *std. Error* | *std. Beta* | *CI* | *Statistic* | *p* | *p*  *(FDR.adj)* | *Estimates* | *std. Error* | *std. Beta* | *CI* | *Statistic* | *p* | *p*  *(FDR.adj)* |
| Intercept | 10.482 | 0.161 | 0.194 | 10.164 – 10.799 | 65.020 | **<0.001** |  | 10.088 | 0.192 | 0.140 | 9.709 – 10.466 | 52.451 | **<0.001** |  |
| Session | -0.132 | 0.077 | -0.030 | -0.284 – 0.020 | -1.711 | 0.089 | 0.119 | -0.044 | 0.100 | 0.003 | -0.242 – 0.154 | -0.441 | 0.660 | 0.660 |
| Age² | -0.001 | 0.001 | -0.079 | -0.002 – 0.000 | -1.453 | 0.147 | 0.295 | -0.000 | 0.001 | -0.057 | -0.001 – 0.001 | -0.157 | 0.875 | 0.875 |
| Sex | -0.289 | 0.141 | -0.339 | -0.568 – -0.010 | -2.050 | **0.042** | 0.169 | -0.227 | 0.151 | -0.246 | -0.526 – 0.071 | -1.504 | 0.135 | 0.205 |
| Session X Age² | 0.000 | 0.000 | -0.003 | -0.000 – 0.001 | 1.307 | 0.193 | 0.387 | 0.000 | 0.000 | -0.020 | -0.001 – 0.001 | 0.161 | 0.873 | 0.873 |
| **Random Effects** | | | | | | | | | | | | | | |
| σ^2^ | 0.19 | | | | | | | 0.33 | | | | | | |
| τ_00_ | 0.56 _participant_id_ | | | | | | | 0.59 _participant_id_ | | | | | | |
| ICC | 0.74 | | | | | | | 0.64 | | | | | | |
| N | 147 _participant_id_ | | | | | | | 147 _participant_id_ | | | | | | |
| Observations | 294 | | | | | | | 294 | | | | | | |
| Marginal R^2^ / Conditional R^2^ | 0.029 / 0.749 | | | | | | | 0.013 / 0.646 | | | | | | |
| AIC | 680.787 | | | | | | | 776.085 | | | | | | |

Note: Repeated-measured EEG data nested within participant, indicated by participant_id. Sex coded as 0 = female, 1 = male.

**Table S24**

*Hierarchical Linear Models of Alpha Power Stability (Eyes-Closed) with Age^2^*

|  | **Occipitoparietal** | | | | | | | **Frontotemporal** | | | | | | |
| --- | --- | --- | --- | --- | --- | --- | --- | --- | --- | --- | --- | --- | --- | --- |
| *Predictors* | *Estimates* | *std. Error* | *std. Beta* | *CI* | *Statistic* | *p* | *p*  *(FDR.adj)* | *Estimates* | *std. Error* | *std. Beta* | *CI* | *Statistic* | *p* | *p*  *(FDR.adj)* |
| Intercept | 1.257 | 0.060 | 0.061 | 1.139 – 1.374 | 21.106 | **<0.001** |  | 0.912 | 0.051 | 0.008 | 0.812 – 1.012 | 17.899 | **<0.001** |  |
| Session | 0.013 | 0.017 | 0.014 | -0.021 – 0.046 | 0.736 | 0.463 | 0.463 | 0.034 | 0.015 | 0.044 | 0.005 – 0.063 | 2.347 | **0.020** | 0.080 |
| Age² | -0.000 | 0.000 | 0.000 | -0.000 – 0.000 | -0.523 | 0.602 | 0.602 | 0.000 | 0.000 | 0.051 | -0.000 – 0.000 | 0.562 | 0.575 | 0.602 |
| Sex | -0.077 | 0.070 | -0.177 | -0.215 – 0.061 | -1.107 | 0.270 | 0.422 | -0.066 | 0.060 | -0.169 | -0.184 – 0.052 | -1.104 | 0.271 | 0.422 |
| **Random Effects** | | | | | | | | | | | | | | |
| σ^2^ | 0.03 | | | | | | | 0.02 | | | | | | |
| τ_00_ | 0.18 _participant_id_ | | | | | | | 0.13 _participant_id_ | | | | | | |
| ICC | 0.87 | | | | | | | 0.88 | | | | | | |
| N | 175 _participant_id_ | | | | | | | 175 _participant_id_ | | | | | | |
| Observations | 350 | | | | | | | 350 | | | | | | |
| Marginal R^2^ / Conditional R^2^ | 0.009 / 0.876 | | | | | | | 0.010 / 0.878 | | | | | | |
| AIC | 224.368 | | | | | | | 114.791 | | | | | | |

Note: Repeated-measured EEG data nested within participant, indicated by participant_id. Sex coded as 0 = female, 1 = male.

**Table S25**

*Hierarchical Linear Models of Alpha Power Stability (Eyes-Open) with Age^2^*

|  | **Occipitoparietal** | | | | | | | **Frontotemporal** | | | | | | |
| --- | --- | --- | --- | --- | --- | --- | --- | --- | --- | --- | --- | --- | --- | --- |
| *Predictors* | *Estimates* | *std. Error* | *std. Beta* | *CI* | *Statistic* | *p* | *p*  *(FDR.adj)* | *Estimates* | *std. Error* | *std. Beta* | *CI* | *Statistic* | *p* | *p*  *(FDR.adj)* |
| Intercept | 0.690 | 0.051 | -0.023 | 0.590 – 0.790 | 13.589 | **<0.001** |  | 0.491 | 0.037 | -0.032 | 0.419 – 0.564 | 13.345 | **<0.001** |  |
| Session | -0.015 | 0.017 | -0.022 | -0.047 – 0.018 | -0.892 | 0.374 | 0.463 | 0.011 | 0.013 | 0.023 | -0.014 – 0.037 | 0.863 | 0.389 | 0.463 |
| Age² | 0.000 | 0.000 | 0.056 | -0.000 – 0.000 | 1.019 | 0.310 | 0.602 | 0.000 | 0.000 | 0.088 | -0.000 – 0.000 | 1.872 | 0.063 | 0.253 |
| Sex | -0.032 | 0.056 | -0.095 | -0.142 – 0.078 | -0.576 | 0.565 | 0.565 | -0.039 | 0.039 | -0.164 | -0.117 – 0.038 | -1.005 | 0.317 | 0.422 |
| **Random Effects** | | | | | | | | | | | | | | |
| σ^2^ | 0.03 | | | | | | | 0.02 | | | | | | |
| τ_00_ | 0.18 _participant_id_ | | | | | | | 0.13 _participant_id_ | | | | | | |
| ICC | 0.87 | | | | | | | 0.88 | | | | | | |
| N | 175 _participant_id_ | | | | | | | 175 _participant_id_ | | | | | | |
| Observations | 350 | | | | | | | 350 | | | | | | |
| Marginal R^2^ / Conditional R^2^ | 0.009 / 0.876 | | | | | | | 0.010 / 0.878 | | | | | | |
| AIC | 224.368 | | | | | | | 114.791 | | | | | | |

Note: Repeated-measured EEG data nested within participant, indicated by participant_id. Sex coded as 0 = female, 1 = male.

**Table S26**

*Hierarchical Linear Models of Alpha Peak Power Stability, with Session X Age^2^ interactions (Eyes-Closed)*

|  | **Occipitoparietal** | | | | | | | **Frontotemporal** | | | | | | |
| --- | --- | --- | --- | --- | --- | --- | --- | --- | --- | --- | --- | --- | --- | --- |
| *Predictors* | *Estimates* | *std. Error* | *std. Beta* | *CI* | *Statistic* | *p* | *p*  *(FDR.adj)* | *Estimates* | *std. Error* | *std. Beta* | *CI* | *Statistic* | *p* | *p*  *(FDR.adj)* |
| Intercept | 1.251 | 0.066 | 0.061 | 1.122 – 1.381 | 18.999 | **<0.001** |  | 0.900 | 0.056 | 0.008 | 0.789 – 1.010 | 15.989 | **<0.001** |  |
| Session | 0.016 | 0.025 | 0.019 | -0.034 – 0.066 | 0.640 | 0.523 | 0.523 | 0.042 | 0.022 | 0.043 | -0.000 – 0.085 | 1.960 | 0.052 | 0.206 |
| Age² | -0.000 | 0.000 | 0.000 | -0.001 – 0.000 | -0.294 | 0.769 | 0.769 | 0.000 | 0.000 | 0.051 | -0.000 – 0.001 | 0.756 | 0.450 | 0.600 |
| Sex | -0.077 | 0.070 | -0.177 | -0.215 – 0.061 | -1.107 | 0.270 | 0.422 | -0.066 | 0.060 | -0.169 | -0.184 – 0.052 | -1.104 | 0.271 | 0.422 |
| Session X Age² | -0.000 | 0.000 | -0.005 | -0.000 – 0.000 | -0.195 | 0.846 | 0.846 | -0.000 | 0.000 | 0.001 | -0.000 – 0.000 | -0.509 | 0.611 | 0.815 |
| **Random Effects** | | | | | | | | | | | | | | |
| σ^2^ | 0.03 | | | | | | | 0.02 | | | | | | |
| τ_00_ | 0.18 _participant_id_ | | | | | | | 0.13 _participant_id_ | | | | | | |
| ICC | 0.87 | | | | | | | 0.88 | | | | | | |
| N | 175 _participant_id_ | | | | | | | 175 _participant_id_ | | | | | | |
| Observations | 350 | | | | | | | 350 | | | | | | |
| Marginal R^2^ / Conditional R^2^ | 0.009 / 0.875 | | | | | | | 0.010 / 0.877 | | | | | | |
| AIC | 243.050 | | | | | | | 133.577 | | | | | | |

Note: Repeated-measured EEG data nested within participant, indicated by participant_id. Sex coded as 0 = female, 1 = male.

**Table S27**

*Hierarchical Linear Models of Alpha Peak Power Stability, with Session X Age^2^ interactions (Eyes-Open)*

|  | **Occipitoparietal** | | | | | | | **Frontotemporal** | | | | | | |
| --- | --- | --- | --- | --- | --- | --- | --- | --- | --- | --- | --- | --- | --- | --- |
| *Predictors* | *Estimates* | *std. Error* | *std. Beta* | *CI* | *Statistic* | *p* | *p*  *(FDR.adj)* | *Estimates* | *std. Error* | *std. Beta* | *CI* | *Statistic* | *p* | *p*  *(FDR.adj)* |
| Intercept | 0.643 | 0.058 | -0.023 | 0.529 – 0.756 | 11.148 | **<0.001** |  | 0.470 | 0.043 | -0.032 | 0.386 – 0.554 | 11.004 | **<0.001** |  |
| Session | 0.017 | 0.025 | -0.005 | -0.032 – 0.066 | 0.692 | 0.490 | 0.523 | 0.025 | 0.019 | 0.034 | -0.013 – 0.064 | 1.293 | 0.198 | 0.396 |
| Age² | 0.000 | 0.000 | 0.056 | -0.000 – 0.001 | 1.929 | 0.055 | 0.109 | 0.000 | 0.000 | 0.088 | 0.000 – 0.001 | 1.989 | **0.048** | 0.109 |
| Sex | -0.032 | 0.056 | -0.095 | -0.142 – 0.078 | -0.576 | 0.565 | 0.565 | -0.039 | 0.039 | -0.164 | -0.117 – 0.038 | -1.005 | 0.317 | 0.422 |
| Session X Age² | -0.000 | 0.000 | -0.017 | -0.000 – 0.000 | -1.736 | 0.085 | 0.339 | -0.000 | 0.000 | -0.010 | -0.000 – 0.000 | -0.963 | 0.337 | 0.674 |
| **Random Effects** | | | | | | | | | | | | | | |
| σ^2^ | 0.02 | | | | | | | 0.01 | | | | | | |
| τ_00_ | 0.09 _participant_id_ | | | | | | | 0.04 _participant_id_ | | | | | | |
| ICC | 0.82 | | | | | | | 0.78 | | | | | | |
| N | 147 _participant_id_ | | | | | | | 147 _participant_id_ | | | | | | |
| Observations | 294 | | | | | | | 294 | | | | | | |
| Marginal R^2^ / Conditional R^2^ | 0.011 / 0.826 | | | | | | | 0.029 / 0.788 | | | | | | |
| AIC | 81.415 | | | | | | | -87.210 | | | | | | |

Note: Repeated-measured EEG data nested within participant, indicated by participant_id. Sex coded as 0 = female, 1 = male.

**Table S28**

*Hierarchical Linear Models of Aperiodic Exponent Stability, with Session X Sex interactions (Eyes-Closed)*

|  | **Midline** | | | | | | | **Perimeter** | | | | | | |
| --- | --- | --- | --- | --- | --- | --- | --- | --- | --- | --- | --- | --- | --- | --- |
| *Predictors* | *Estimates* | *std. Error* | *std. Beta* | *CI* | *Statistic* | *p* | *p*  *(FDR.adj)* | *Estimates* | *std. Error* | *std. Beta* | *CI* | *Statistic* | *p* | *p*  *(FDR.adj)* |
| Intercept | 1.346 | 0.050 | 0.080 | 1.247 – 1.445 | 26.761 | **<0.001** |  | 1.055 | 0.049 | 0.069 | 0.959 – 1.151 | 21.608 | **<0.001** |  |
| Session | -0.085 | 0.029 | -0.128 | -0.142 – -0.028 | -2.928 | **0.004** | **0.015** | -0.070 | 0.028 | -0.108 | -0.125 – -0.016 | -2.554 | **0.011** | **0.023** |
| Age | -0.007 | 0.001 | -0.301 | -0.010 – -0.004 | -4.878 | **<0.001** | **<0.001** | -0.005 | 0.002 | -0.229 | -0.008 – -0.002 | -3.542 | **0.001** | **0.001** |
| Sex | -0.212 | 0.085 | -0.230 | -0.380 – -0.044 | -2.483 | **0.014** | 0.054 | -0.174 | 0.083 | -0.197 | -0.337 – -0.011 | -2.103 | **0.036** | 0.073 |
| Session X Sex | 0.090 | 0.049 | 0.137 | -0.007 – 0.187 | 1.840 | 0.067 | 0.237 | 0.073 | 0.047 | 0.112 | -0.019 – 0.165 | 1.568 | 0.119 | 0.237 |
| **Random Effects** | | | | | | | | | | | | | | |
| σ^2^ | 0.05 | | | | | | | 0.04 | | | | | | |
| τ_00_ | 0.05 _participant_id_ | | | | | | | 0.06 _participant_id_ | | | | | | |
| ICC | 0.50 | | | | | | | 0.56 | | | | | | |
| N | 178 _participant_id_ | | | | | | | 178 _participant_id_ | | | | | | |
| Observations | 356 | | | | | | | 356 | | | | | | |
| Marginal R^2^ / Conditional R^2^ | 0.120 / 0.561 | | | | | | | 0.073 / 0.594 | | | | | | |
| AIC | 171.393 | | | | | | | 164.145 | | | | | | |

Note: Repeated-measured EEG data nested within participant, indicated by participant_id. Sex coded as 0 = female, 1 = male.

**Table S29**

*Hierarchical Linear Models of Aperiodic Exponent Stability, with Session X Sex interactions (Eyes-Open)*

|  | **Midline** | | | | | | | **Perimeter** | | | | | | |
| --- | --- | --- | --- | --- | --- | --- | --- | --- | --- | --- | --- | --- | --- | --- |
| *Predictors* | *Estimates* | *std. Error* | *std. Beta* | *CI* | *Statistic* | *p* | *p*  *(FDR.adj)* | *Estimates* | *std. Error* | *std. Beta* | *CI* | *Statistic* | *p* | *p*  *(FDR.adj)* |
| Intercept | 1.251 | 0.043 | 0.053 | 1.167 – 1.335 | 29.350 | **<0.001** |  | 0.908 | 0.041 | -0.026 | 0.827 – 0.989 | 21.999 | **<0.001** |  |
| Session | -0.054 | 0.024 | -0.095 | -0.101 – -0.007 | -2.285 | **0.024** | **0.024** | -0.053 | 0.023 | -0.097 | -0.098 – -0.009 | -2.367 | **0.019** | **0.024** |
| Age | -0.006 | 0.001 | -0.287 | -0.009 – -0.003 | -4.248 | **<0.001** | **<0.001** | -0.004 | 0.001 | -0.213 | -0.007 – -0.002 | -3.053 | **0.003** | **0.003** |
| Sex | 0.024 | 0.073 | -0.155 | -0.119 – 0.168 | 0.333 | 0.739 | 0.739 | 0.088 | 0.071 | 0.077 | -0.051 – 0.228 | 1.252 | 0.212 | 0.282 |
| Session X Sex | -0.046 | 0.040 | -0.080 | -0.126 – 0.034 | -1.133 | 0.259 | 0.259 | -0.045 | 0.039 | -0.081 | -0.121 – 0.031 | -1.161 | 0.247 | 0.259 |
| **Random Effects** | | | | | | | | | | | | | | |
| σ^2^ | 0.03 | | | | | | | 0.03 | | | | | | |
| τ_00_ | 0.04 _participant_id_ | | | | | | | 0.05 _participant_id_ | | | | | | |
| ICC | 0.60 | | | | | | | 0.63 | | | | | | |
| N | 161 _participant_id_ | | | | | | | 161 _participant_id_ | | | | | | |
| Observations | 322 | | | | | | | 322 | | | | | | |
| Marginal R^2^ / Conditional R^2^ | 0.108 / 0.642 | | | | | | | 0.061 / 0.649 | | | | | | |
| AIC | 44.551 | | | | | | | 28.940 | | | | | | |

Note: Repeated-measured EEG data nested within participant, indicated by participant_id. Sex coded as 0 = female, 1 = male.

**Table S30**

*Hierarchical Linear Models of Aperiodic Offset Stability, with Session X Sex interactions (Eyes-Closed)*

|  | **Midline** | | | | | | | **Perimeter** | | | | | | |
| --- | --- | --- | --- | --- | --- | --- | --- | --- | --- | --- | --- | --- | --- | --- |
| *Predictors* | *Estimates* | *std. Error* | *std. Beta* | *CI* | *Statistic* | *p* | *p*  *(FDR.adj)* | *Estimates* | *std. Error* | *std. Beta* | *CI* | *Statistic* | *p* | *p*  *(FDR.adj)* |
| Intercept | 1.291 | 0.057 | 0.115 | 1.178 – 1.404 | 22.500 | **<0.001** |  | 0.840 | 0.053 | 0.107 | 0.736 – 0.944 | 15.905 | **<0.001** |  |
| Session | -0.130 | 0.031 | -0.151 | -0.192 – -0.069 | -4.207 | **<0.001** | **<0.001** | -0.108 | 0.027 | -0.133 | -0.163 – -0.054 | -3.945 | **<0.001** | **<0.001** |
| Age | -0.010 | 0.002 | -0.328 | -0.014 – -0.006 | -5.190 | **<0.001** | **<0.001** | -0.008 | 0.002 | -0.278 | -0.012 – -0.004 | -4.224 | **<0.001** | **<0.001** |
| Sex | -0.259 | 0.097 | -0.329 | -0.450 – -0.067 | -2.659 | **0.008** | **0.033** | -0.201 | 0.090 | -0.307 | -0.377 – -0.025 | -2.245 | **0.025** | 0.051 |
| Session X Sex | 0.078 | 0.053 | 0.091 | -0.026 – 0.182 | 1.486 | 0.139 | 0.197 | 0.051 | 0.047 | 0.062 | -0.041 – 0.143 | 1.090 | 0.277 | 0.277 |
| **Random Effects** | | | | | | | | | | | | | | |
| σ^2^ | 0.06 | | | | | | | 0.04 | | | | | | |
| τ_00_ | 0.10 _participant_id_ | | | | | | | 0.10 _participant_id_ | | | | | | |
| ICC | 0.65 | | | | | | | 0.70 | | | | | | |
| N | 178 _participant_id_ | | | | | | | 178 _participant_id_ | | | | | | |
| Observations | 356 | | | | | | | 356 | | | | | | |
| Marginal R^2^ / Conditional R^2^ | 0.159 / 0.704 | | | | | | | 0.120 / 0.739 | | | | | | |
| AIC | 295.142 | | | | | | | 246.215 | | | | | | |

Note: Repeated-measured EEG data nested within participant, indicated by participant_id. Sex coded as 0 = female, 1 = male.

**Table S31**

*Hierarchical Linear Models of Aperiodic Offset Stability, with Session X Age interactions (Eyes-Open)*

|  | **Midline** | | | | | | | **Perimeter** | | | | | | |
| --- | --- | --- | --- | --- | --- | --- | --- | --- | --- | --- | --- | --- | --- | --- |
| *Predictors* | *Estimates* | *std. Error* | *std. Beta* | *CI* | *Statistic* | *p* | *p*  *(FDR.adj)* | *Estimates* | *std. Error* | *std. Beta* | *CI* | *Statistic* | *p* | *p*  *(FDR.adj)* |
| Intercept | 1.126 | 0.052 | 0.103 | 1.024 – 1.227 | 21.807 | **<0.001** |  | 0.665 | 0.047 | 0.063 | 0.574 – 0.757 | 14.273 | **<0.001** |  |
| Session | -0.073 | 0.028 | -0.099 | -0.128 – -0.018 | -2.611 | **0.010** | **0.013** | -0.057 | 0.025 | -0.087 | -0.106 – -0.007 | -2.255 | **0.025** | **0.025** |
| Age | -0.008 | 0.002 | -0.313 | -0.012 – -0.005 | -4.657 | **<0.001** | **<0.001** | -0.007 | 0.002 | -0.280 | -0.010 – -0.003 | -4.083 | **<0.001** | **<0.001** |
| Sex | -0.008 | 0.088 | -0.302 | -0.182 – 0.166 | -0.087 | 0.931 | 0.931 | 0.065 | 0.080 | -0.186 | -0.092 – 0.222 | 0.810 | 0.419 | 0.558 |
| Session X Sex | -0.069 | 0.048 | -0.094 | -0.163 – 0.025 | -1.455 | 0.148 | 0.197 | -0.083 | 0.043 | -0.128 | -0.168 – 0.001 | -1.942 | 0.054 | 0.197 |
| **Random Effects** | | | | | | | | | | | | | | |
| σ^2^ | 0.04 | | | | | | | 0.03 | | | | | | |
| τ_00_ | 0.08 _participant_id_ | | | | | | | 0.06 _participant_id_ | | | | | | |
| ICC | 0.65 | | | | | | | 0.65 | | | | | | |
| N | 161 _participant_id_ | | | | | | | 161 _participant_id_ | | | | | | |
| Observations | 322 | | | | | | | 322 | | | | | | |
| Marginal R^2^ / Conditional R^2^ | 0.146 / 0.702 | | | | | | | 0.111 / 0.689 | | | | | | |
| AIC | 175.276 | | | | | | | 110.496 | | | | | | |

Note: Repeated-measured EEG data nested within participant, indicated by participant_id. Sex coded as 0 = female, 1 = male.

**Table S32**

*Hierarchical Linear Models of Individual Alpha Peak Frequency Stability, with Session X Sex interactions (Eyes-Closed)*

|  | **Occipitoparietal** | | | | | | | **Frontotemporal** | | | | | | |
| --- | --- | --- | --- | --- | --- | --- | --- | --- | --- | --- | --- | --- | --- | --- |
| *Predictors* | *Estimates* | *std. Error* | *std. Beta* | *CI* | *Statistic* | *p* | *p*  *(FDR.adj)* | *Estimates* | *std. Error* | *std. Beta* | *CI* | *Statistic* | *p* | *p*  *(FDR.adj)* |
| Intercept | 10.265 | 0.093 | 0.057 | 10.083 – 10.447 | 110.915 | **<0.001** |  | 9.989 | 0.100 | -0.001 | 9.793 – 10.185 | 100.266 | **<0.001** |  |
| Session | -0.087 | 0.041 | -0.054 | -0.168 – -0.007 | -2.134 | **0.034** | 0.137 | -0.073 | 0.048 | -0.045 | -0.168 – 0.022 | -1.510 | 0.133 | 0.265 |
| Age | -0.015 | 0.004 | -0.267 | -0.023 – -0.007 | -3.787 | **<0.001** | **<0.001** | -0.021 | 0.004 | -0.348 | -0.028 – -0.013 | -5.138 | **<0.001** | **<0.001** |
| Sex | -0.027 | 0.157 | -0.165 | -0.336 – 0.282 | -0.173 | 0.863 | 0.877 | 0.052 | 0.169 | 0.003 | -0.280 – 0.385 | 0.309 | 0.758 | 0.877 |
| Session X Sex | -0.070 | 0.069 | -0.044 | -0.206 – 0.067 | -1.009 | 0.315 | 0.629 | -0.033 | 0.082 | -0.020 | -0.195 – 0.128 | -0.408 | 0.684 | 0.684 |
| **Random Effects** | | | | | | | | | | | | | | |
| σ^2^ | 0.09 | | | | | | | 0.13 | | | | | | |
| τ_00_ | 0.50 _participant_id_ | | | | | | | 0.46 _participant_id_ | | | | | | |
| ICC | 0.84 | | | | | | | 0.78 | | | | | | |
| N | 175 _participant_id_ | | | | | | | 175 _participant_id_ | | | | | | |
| Observations | 350 | | | | | | | 350 | | | | | | |
| Marginal R^2^ / Conditional R^2^ | 0.087 / 0.854 | | | | | | | 0.122 / 0.805 | | | | | | |
| AIC | 629.376 | | | | | | | 682.631 | | | | | | |

Note: Repeated-measured EEG data nested within participant, indicated by participant_id. Sex coded as 0 = female, 1 = male.

**Table S33**

*Hierarchical Linear Models of Individual Alpha Peak Frequency Stability, with Session X Sex interactions (Eyes-Open)*

|  | **Occipitoparietal** | | | | | | | **Frontotemporal** | | | | | | |
| --- | --- | --- | --- | --- | --- | --- | --- | --- | --- | --- | --- | --- | --- | --- |
| *Predictors* | *Estimates* | *std. Error* | *std. Beta* | *CI* | *Statistic* | *p* | *p*  *(FDR.adj)* | *Estimates* | *std. Error* | *std. Beta* | *CI* | *Statistic* | *p* | *p*  *(FDR.adj)* |
| Intercept | 10.230 | 0.125 | 0.102 | 9.984 – 10.476 | 81.794 | **<0.001** |  | 10.038 | 0.151 | 0.070 | 9.741 – 10.335 | 66.570 | **<0.001** |  |
| Session | 0.010 | 0.063 | 0.006 | -0.114 – 0.135 | 0.161 | 0.872 | 0.996 | 0.000 | 0.082 | 0.000 | -0.162 – 0.163 | 0.004 | 0.996 | 0.996 |
| Age | -0.009 | 0.005 | -0.137 | -0.018 – 0.001 | -1.798 | 0.074 | 0.074 | -0.012 | 0.005 | -0.169 | -0.022 – -0.002 | -2.283 | **0.024** | **0.032** |
| Sex | 0.033 | 0.214 | -0.300 | -0.389 – 0.455 | 0.155 | 0.877 | 0.877 | -0.053 | 0.258 | -0.206 | -0.562 – 0.456 | -0.204 | 0.838 | 0.877 |
| Session X Sex | -0.197 | 0.108 | -0.113 | -0.411 – 0.016 | -1.825 | 0.070 | 0.280 | -0.096 | 0.141 | -0.050 | -0.374 – 0.183 | -0.679 | 0.498 | 0.664 |
| **Random Effects** | | | | | | | | | | | | | | |
| σ^2^ | 0.19 | | | | | | | 0.33 | | | | | | |
| τ_00_ | 0.55 _participant_id_ | | | | | | | 0.56 _participant_id_ | | | | | | |
| ICC | 0.74 | | | | | | | 0.63 | | | | | | |
| N | 147 _participant_id_ | | | | | | | 147 _participant_id_ | | | | | | |
| Observations | 294 | | | | | | | 294 | | | | | | |
| Marginal R^2^ / Conditional R^2^ | 0.046 / 0.752 | | | | | | | 0.041 / 0.647 | | | | | | |
| AIC | 659.522 | | | | | | | 753.557 | | | | | | |

Note: Repeated-measured EEG data nested within participant, indicated by participant_id. Sex coded as 0 = female, 1 = male.

**Table S34**

*Hierarchical Linear Models of Alpha Peak Power Stability, with Session X Sex interactions (Eyes-Closed)*

|  | **Occipitoparietal** | | | | | | | **Frontotemporal** | | | | | | |
| --- | --- | --- | --- | --- | --- | --- | --- | --- | --- | --- | --- | --- | --- | --- |
| *Predictors* | *Estimates* | *std. Error* | *std. Beta* | *CI* | *Statistic* | *p* | *p*  *(FDR.adj)* | *Estimates* | *std. Error* | *std. Beta* | *CI* | *Statistic* | *p* | *p*  *(FDR.adj)* |
| Intercept | 1.248 | 0.052 | 0.050 | 1.146 – 1.351 | 24.012 | **<0.001** |  | 0.926 | 0.045 | 0.053 | 0.838 – 1.013 | 20.751 | **<0.001** |  |
| Session | 0.010 | 0.021 | 0.011 | -0.032 – 0.052 | 0.474 | 0.636 | 0.748 | 0.037 | 0.018 | 0.048 | 0.002 – 0.073 | 2.068 | **0.040** | 0.160 |
| Age | -0.004 | 0.002 | -0.122 | -0.009 – 0.001 | -1.657 | 0.099 | 0.397 | -0.001 | 0.002 | -0.048 | -0.005 – 0.003 | -0.642 | 0.522 | 0.696 |
| Sex | -0.076 | 0.088 | -0.144 | -0.250 – 0.097 | -0.863 | 0.389 | 0.738 | -0.045 | 0.076 | -0.152 | -0.194 – 0.104 | -0.594 | 0.553 | 0.738 |
| Session X Sex | 0.007 | 0.036 | 0.008 | -0.064 – 0.078 | 0.201 | 0.841 | 0.841 | -0.009 | 0.031 | -0.012 | -0.070 – 0.051 | -0.303 | 0.763 | 0.841 |
| **Random Effects** | | | | | | | | | | | | | | |
| σ^2^ | 0.03 | | | | | | | 0.02 | | | | | | |
| τ_00_ | 0.18 _participant_id_ | | | | | | | 0.13 _participant_id_ | | | | | | |
| ICC | 0.87 | | | | | | | 0.88 | | | | | | |
| N | 175 _participant_id_ | | | | | | | 175 _participant_id_ | | | | | | |
| Observations | 350 | | | | | | | 350 | | | | | | |
| Marginal R^2^ / Conditional R^2^ | 0.022 / 0.875 | | | | | | | 0.010 / 0.877 | | | | | | |
| AIC | 223.503 | | | | | | | 116.556 | | | | | | |

Note: Repeated-measured EEG data nested within participant, indicated by participant_id. Sex coded as 0 = female, 1 = male.

**Table S35**

*Hierarchical Linear Models of Alpha Peak Power Stability, with Session X Sex interactions (Eyes-Open)*

|  | **Occipitoparietal** | | | | | | | **Frontotemporal** | | | | | | |
| --- | --- | --- | --- | --- | --- | --- | --- | --- | --- | --- | --- | --- | --- | --- |
| *Predictors* | *Estimates* | *std. Error* | *std. Beta* | *CI* | *Statistic* | *p* | *p*  *(FDR.adj)* | *Estimates* | *std. Error* | *std. Beta* | *CI* | *Statistic* | *p* | *p*  *(FDR.adj)* |
| Intercept | 0.713 | 0.045 | 0.029 | 0.625 – 0.801 | 15.971 | **<0.001** |  | 0.527 | 0.033 | 0.062 | 0.461 – 0.593 | 15.747 | **<0.001** |  |
| Session | -0.007 | 0.020 | -0.010 | -0.047 – 0.034 | -0.322 | 0.748 | 0.748 | 0.013 | 0.016 | 0.028 | -0.018 – 0.045 | 0.835 | 0.405 | 0.748 |
| Age | -0.002 | 0.002 | -0.100 | -0.006 – 0.001 | -1.260 | 0.210 | 0.419 | 0.001 | 0.001 | 0.030 | -0.002 – 0.003 | 0.381 | 0.703 | 0.703 |
| Sex | 0.008 | 0.077 | -0.085 | -0.143 – 0.158 | 0.102 | 0.919 | 0.919 | -0.034 | 0.057 | -0.183 | -0.147 – 0.078 | -0.599 | 0.550 | 0.738 |
| Session X Sex | -0.024 | 0.035 | -0.036 | -0.093 – 0.045 | -0.688 | 0.492 | 0.841 | -0.006 | 0.028 | -0.013 | -0.061 – 0.048 | -0.234 | 0.815 | 0.841 |
| **Random Effects** | | | | | | | | | | | | | | |
| σ^2^ | 0.02 | | | | | | | 0.01 | | | | | | |
| τ_00_ | 0.09 _participant_id_ | | | | | | | 0.05 _participant_id_ | | | | | | |
| ICC | 0.82 | | | | | | | 0.79 | | | | | | |
| N | 147 _participant_id_ | | | | | | | 147 _participant_id_ | | | | | | |
| Observations | 294 | | | | | | | 294 | | | | | | |
| Marginal R^2^ / Conditional R^2^ | 0.013 / 0.822 | | | | | | | 0.008 / 0.787 | | | | | | |
| AIC | 66.404 | | | | | | | -100.001 | | | | | | |

Note: Repeated-measured EEG data nested within participant, indicated by participant_id. Sex coded as 0 = female, 1 = male.

**Table S36**

*Hierarchical Linear Models of Absolute Alpha Power Stability (Eyes-Closed)*

|  | **Midline** | | | | | | | **Perimeter** | | | | | | |
| --- | --- | --- | --- | --- | --- | --- | --- | --- | --- | --- | --- | --- | --- | --- |
| *Predictors* | *Estimates* | *std. Error* | *std. Beta* | *CI* | *Statistic* | *p* | *p*  *(FDR.adj)* | *Estimates* | *std. Error* | *std. Beta* | *CI* | *Statistic* | *p* | *p*  *(FDR.adj)* |
| Intercept | 1.469 | 0.047 | 0.104 | 1.377 – 1.562 | 31.307 | **<0.001** |  | 1.148 | 0.042 | 0.126 | 1.066 – 1.231 | 27.299 | **<0.001** |  |
| Session | -0.026 | 0.015 | -0.029 | -0.057 – 0.004 | -1.685 | 0.094 | 0.375 | -0.016 | 0.014 | -0.020 | -0.043 – 0.011 | -1.163 | 0.246 | 0.385 |
| Age | -0.004 | 0.002 | -0.122 | -0.009 – 0.001 | -1.665 | 0.098 | 0.391 | -0.002 | 0.002 | -0.065 | -0.006 – 0.002 | -0.891 | 0.374 | 0.499 |
| Sex | -0.135 | 0.069 | -0.298 | -0.271 – 0.002 | -1.939 | 0.054 | 0.065 | -0.146 | 0.062 | -0.362 | -0.269 – -0.024 | -2.352 | **0.020** | 0.065 |
| **Random Effects** | | | | | | | | | | | | | | |
| σ^2^ | 0.02 | | | | | | | 0.02 | | | | | | |
| τ_00_ | 0.18 _participant_id_ | | | | | | | 0.14 _participant_id_ | | | | | | |
| ICC | 0.89 | | | | | | | 0.90 | | | | | | |
| N | 175 _participant_id_ | | | | | | | 175 _participant_id_ | | | | | | |
| Observations | 350 | | | | | | | 350 | | | | | | |
| Marginal R^2^ / Conditional R^2^ | 0.040 / 0.899 | | | | | | | 0.037 / 0.899 | | | | | | |
| AIC | 179.248 | | | | | | | 102.733 | | | | | | |

Note: Repeated-measured EEG data nested within participant, indicated by participant_id. Sex coded as 0 = female, 1 = male.

**Table S37**

*Hierarchical Linear Models of Absolute Alpha Power Stability (Eyes-Open)*

|  | **Midline** | | | | | | | **Perimeter** | | | | | | |
| --- | --- | --- | --- | --- | --- | --- | --- | --- | --- | --- | --- | --- | --- | --- |
| *Predictors* | *Estimates* | *std. Error* | *std. Beta* | *CI* | *Statistic* | *p* | *p*  *(FDR.adj)* | *Estimates* | *std. Error* | *std. Beta* | *CI* | *Statistic* | *p* | *p*  *(FDR.adj)* |
| Intercept | 1.024 | 0.043 | 0.105 | 0.939 – 1.109 | 23.749 | **<0.001** |  | 0.727 | 0.041 | 0.117 | 0.647 – 0.807 | 17.839 | **<0.001** |  |
| Session | -0.018 | 0.017 | -0.026 | -0.052 – 0.016 | -1.065 | 0.288 | 0.385 | 0.012 | 0.016 | 0.018 | -0.019 – 0.044 | 0.766 | 0.445 | 0.445 |
| Age | -0.002 | 0.002 | -0.076 | -0.006 – 0.002 | -0.963 | 0.337 | 0.499 | 0.000 | 0.002 | 0.004 | -0.004 – 0.004 | 0.052 | 0.959 | 0.959 |
| Sex | -0.110 | 0.059 | -0.309 | -0.227 – 0.007 | -1.863 | 0.065 | 0.065 | -0.117 | 0.056 | -0.344 | -0.228 – -0.005 | -2.067 | **0.041** | 0.065 |
| **Random Effects** | | | | | | | | | | | | | | |
| σ^2^ | 0.02 | | | | | | | 0.02 | | | | | | |
| τ_00_ | 0.10 _participant_id_ | | | | | | | 0.09 _participant_id_ | | | | | | |
| ICC | 0.83 | | | | | | | 0.83 | | | | | | |
| N | 147 _participant_id_ | | | | | | | 147 _participant_id_ | | | | | | |
| Observations | 294 | | | | | | | 294 | | | | | | |
| Marginal R^2^ / Conditional R^2^ | 0.029 / 0.832 | | | | | | | 0.026 / 0.839 | | | | | | |
| AIC | 88.855 | | | | | | | 53.955 | | | | | | |

Note: Repeated-measured EEG data nested within participant, indicated by participant_id. Sex coded as 0 = female, 1 = male.

**Table S38**

*Hierarchical Linear Models of Absolute Alpha Power Stability, with Session X Age interactions (Eyes-Closed)*

|  | **Midline** | | | | | | | **Perimeter** | | | | | | |
| --- | --- | --- | --- | --- | --- | --- | --- | --- | --- | --- | --- | --- | --- | --- |
| *Predictors* | *Estimates* | *std. Error* | *std. Beta* | *CI* | *Statistic* | *p* | *p*  *(FDR.adj)* | *Estimates* | *std. Error* | *std. Beta* | *CI* | *Statistic* | *p* | *p*  *(FDR.adj)* |
| Intercept | 1.472 | 0.047 | 0.104 | 1.379 – 1.565 | 31.203 | **<0.001** |  | 1.152 | 0.042 | 0.126 | 1.069 – 1.235 | 27.251 | **<0.001** |  |
| Session | -0.028 | 0.016 | -0.029 | -0.059 – 0.003 | -1.769 | 0.079 | 0.256 | -0.018 | 0.014 | -0.020 | -0.046 – 0.009 | -1.310 | 0.192 | 0.256 |
| Age | -0.005 | 0.003 | -0.122 | -0.011 – 0.001 | -1.713 | 0.088 | 0.175 | -0.003 | 0.003 | -0.065 | -0.008 – 0.002 | -1.230 | 0.220 | 0.220 |
| Sex | -0.135 | 0.069 | -0.298 | -0.271 – 0.002 | -1.939 | 0.054 | 0.065 | -0.146 | 0.062 | -0.362 | -0.269 – -0.024 | -2.352 | **0.020** | 0.065 |
| Session X Age | 0.001 | 0.001 | 0.011 | -0.002 – 0.003 | 0.611 | 0.542 | 0.542 | 0.001 | 0.001 | 0.015 | -0.001 – 0.003 | 0.873 | 0.384 | 0.512 |
| **Random Effects** | | | | | | | | | | | | | | |
| σ^2^ | 0.02 | | | | | | | 0.02 | | | | | | |
| τ_00_ | 0.18 _participant_id_ | | | | | | | 0.14 _participant_id_ | | | | | | |
| ICC | 0.89 | | | | | | | 0.90 | | | | | | |
| N | 175 _participant_id_ | | | | | | | 175 _participant_id_ | | | | | | |
| Observations | 350 | | | | | | | 350 | | | | | | |
| Marginal R^2^ / Conditional R^2^ | 0.040 / 0.898 | | | | | | | 0.037 / 0.899 | | | | | | |
| AIC | 192.632 | | | | | | | 115.955 | | | | | | |

Note: Repeated-measured EEG data nested within participant, indicated by participant_id. Sex coded as 0 = female, 1 = male.

**Table S39**

*Hierarchical Linear Models of Absolute Alpha Power Stability, with Session X Age interactions (Eyes-Open)*

|  | **Midline** | | | | | | | **Perimeter** | | | | | | |
| --- | --- | --- | --- | --- | --- | --- | --- | --- | --- | --- | --- | --- | --- | --- |
| *Predictors* | *Estimates* | *std. Error* | *std. Beta* | *CI* | *Statistic* | *p* | *p*  *(FDR.adj)* | *Estimates* | *std. Error* | *std. Beta* | *CI* | *Statistic* | *p* | *p*  *(FDR.adj)* |
| Intercept | 1.033 | 0.043 | 0.105 | 0.947 – 1.118 | 23.842 | **<0.001** |  | 0.737 | 0.041 | 0.117 | 0.656 – 0.817 | 18.034 | **<0.001** |  |
| Session | -0.024 | 0.017 | -0.026 | -0.058 – 0.010 | -1.379 | 0.170 | 0.256 | 0.006 | 0.016 | 0.018 | -0.026 – 0.038 | 0.358 | 0.721 | 0.721 |
| Age | -0.005 | 0.003 | -0.076 | -0.011 – 0.000 | -1.861 | 0.064 | 0.175 | -0.004 | 0.003 | 0.004 | -0.009 – 0.002 | -1.359 | 0.175 | 0.220 |
| Sex | -0.110 | 0.059 | -0.309 | -0.227 – 0.007 | -1.863 | 0.065 | 0.065 | -0.117 | 0.056 | -0.344 | -0.228 – -0.005 | -2.067 | **0.041** | 0.065 |
| Session X Age | 0.002 | 0.001 | 0.041 | -0.000 – 0.005 | 1.705 | 0.090 | 0.181 | 0.002 | 0.001 | 0.049 | 0.000 – 0.005 | 2.105 | **0.037** | 0.148 |
| **Random Effects** | | | | | | | | | | | | | | |
| σ^2^ | 0.02 | | | | | | | 0.02 | | | | | | |
| τ_00_ | 0.10 _participant_id_ | | | | | | | 0.09 _participant_id_ | | | | | | |
| ICC | 0.83 | | | | | | | 0.84 | | | | | | |
| N | 147 _participant_id_ | | | | | | | 147 _participant_id_ | | | | | | |
| Observations | 294 | | | | | | | 294 | | | | | | |
| Marginal R^2^ / Conditional R^2^ | 0.031 / 0.834 | | | | | | | 0.029 / 0.842 | | | | | | |
| AIC | 99.516 | | | | | | | 63.274 | | | | | | |

Note: Repeated-measured EEG data nested within participant, indicated by participant_id. Sex coded as 0 = female, 1 = male.

**Figure S4**

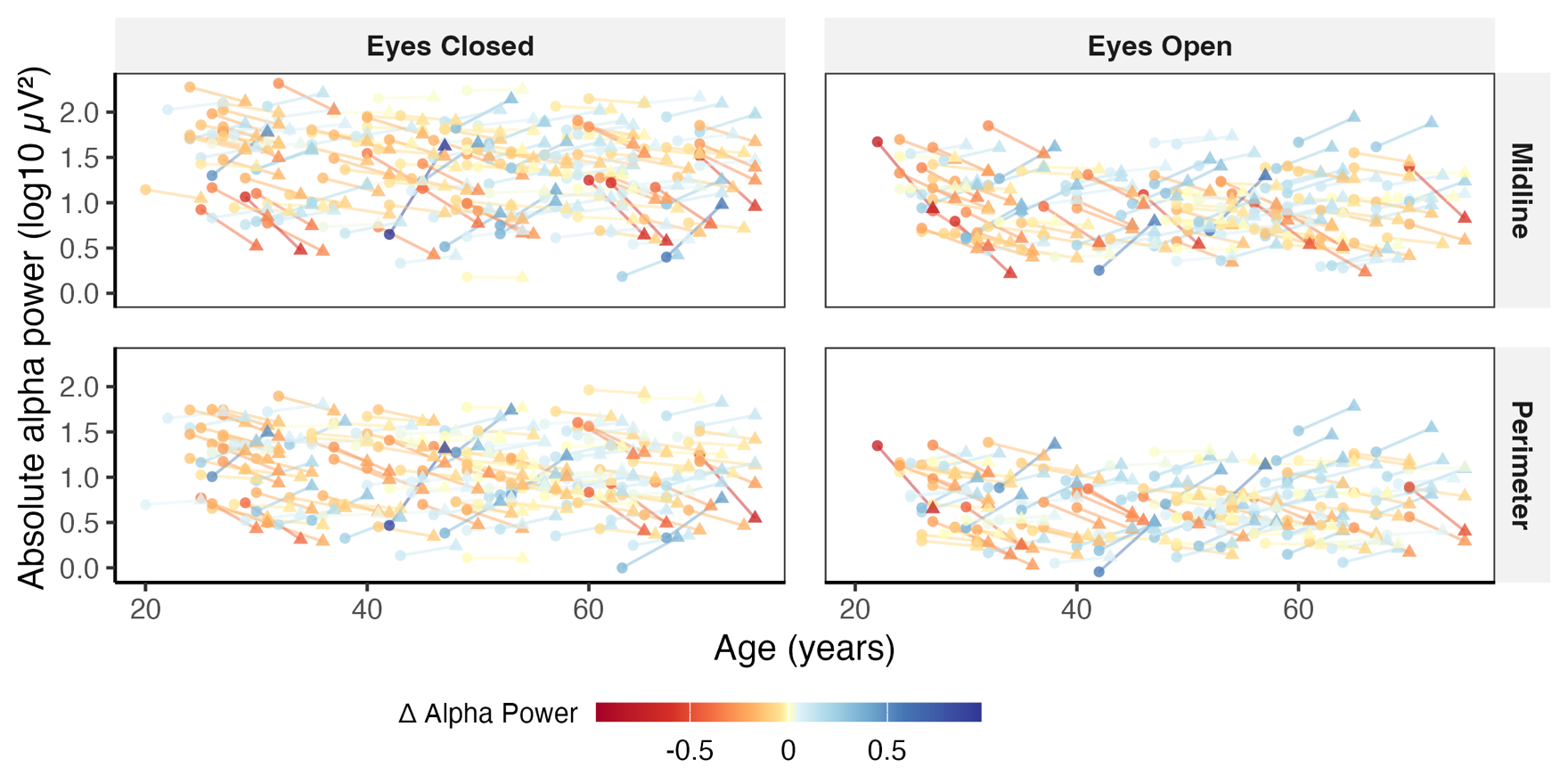
*Absolute alpha power for occipitoparietal and temporal clusters in each session*

Note. Circles indicate each participant’s measure at session 1, and triangles indicate their measure five years later at session 2. Points and lines are shaded to show the direction and magnitude of change between session 1 and session 2.

**Table S40**

*Hierarchical Linear Models of Absolute Alpha Power Stability (Eyes-Closed) with Age^2^*

|  | **Midline** | | | | | | | **Perimeter** | | | | | | |
| --- | --- | --- | --- | --- | --- | --- | --- | --- | --- | --- | --- | --- | --- | --- |
| *Predictors* | *Estimates* | *std. Error* | *std. Beta* | *CI* | *Statistic* | *p* | *p*  *(FDR.adj)* | *Estimates* | *std. Error* | *std. Beta* | *CI* | *Statistic* | *p* | *p*  *(FDR.adj)* |
| Intercept | 1.443 | 0.058 | 0.093 | 1.328 – 1.558 | 24.776 | **<0.001** |  | 1.107 | 0.052 | 0.095 | 1.005 – 1.209 | 21.358 | **<0.001** |  |
| Session | -0.026 | 0.015 | -0.029 | -0.057 – 0.004 | -1.685 | 0.094 | 0.375 | -0.016 | 0.014 | -0.020 | -0.043 – 0.011 | -1.163 | 0.246 | 0.385 |
| Age^2^ | 0.000 | 0.000 | 0.022 | -0.000 – 0.000 | 0.595 | 0.552 | 0.552 | 0.000 | 0.000 | 0.038 | -0.000 – 0.001 | 1.278 | 0.203 | 0.271 |
| Sex | -0.152 | 0.069 | -0.332 | -0.289 – -0.015 | -2.186 | **0.030** | 0.060 | -0.159 | 0.062 | -0.382 | -0.281 – -0.037 | -2.572 | **0.011** | **0.044** |
| **Random Effects** | | | | | | | | | | | | | | |
| σ^2^ | 0.02 | | | | | | | 0.02 | | | | | | |
| τ_00_ | 0.18 _participant_id_ | | | | | | | 0.14 _participant_id_ | | | | | | |
| ICC | 0.90 | | | | | | | 0.89 | | | | | | |
| N | 175 _participant_id_ | | | | | | | 175 _participant_id_ | | | | | | |
| Observations | 350 | | | | | | | 350 | | | | | | |
| Marginal R^2^ / Conditional R^2^ | 0.027 / 0.899 | | | | | | | 0.041 / 0.899 | | | | | | |
| AIC | 186.823 | | | | | | | 107.080 | | | | | | |

Note: Repeated-measured EEG data nested within participant, indicated by participant_id. Sex coded as 0 = female, 1 = male.

**Table S41**

*Hierarchical Linear Models of Absolute Alpha Power Stability (Eyes-Open) with Age^2^*

|  | **Midline** | | | | | | | **Perimeter** | | | | | | |
| --- | --- | --- | --- | --- | --- | --- | --- | --- | --- | --- | --- | --- | --- | --- |
| *Predictors* | *Estimates* | *std. Error* | *std. Beta* | *CI* | *Statistic* | *p* | *p*  *(FDR.adj)* | *Estimates* | *std. Error* | *std. Beta* | *CI* | *Statistic* | *p* | *p*  *(FDR.adj)* |
| Intercept | 0.962 | 0.053 | 0.044 | 0.858 – 1.067 | 18.075 | **<0.001** |  | 0.656 | 0.050 | 0.042 | 0.557 – 0.755 | 13.110 | **<0.001** |  |
| Session | -0.018 | 0.017 | -0.026 | -0.052 – 0.016 | -1.065 | 0.288 | 0.385 | 0.012 | 0.016 | 0.018 | -0.019 – 0.044 | 0.766 | 0.445 | 0.445 |
| Age^2^ | 0.000 | 0.000 | 0.063 | -0.000 – 0.001 | 1.833 | 0.069 | 0.138 | 0.000 | 0.000 | 0.071 | 0.000 – 0.001 | 2.376 | **0.019** | 0.075 |
| Sex | -0.111 | 0.058 | -0.313 | -0.226 – 0.005 | -1.891 | 0.061 | 0.061 | -0.111 | 0.055 | -0.332 | -0.220 – -0.002 | -2.005 | **0.047** | 0.061 |
| **Random Effects** | | | | | | | | | | | | | | |
| σ^2^ | 0.02 | | | | | | | 0.02 | | | | | | |
| τ_00_ | 0.10 _participant_id_ | | | | | | | 0.09 _participant_id_ | | | | | | |
| ICC | 0.82 | | | | | | | 0.83 | | | | | | |
| N | 147 _participant_id_ | | | | | | | 147 _participant_id_ | | | | | | |
| Observations | 294 | | | | | | | 294 | | | | | | |
| Marginal R^2^ / Conditional R^2^ | 0.044 / 0.832 | | | | | | | 0.060 / 0.838 | | | | | | |
| AIC | 91.575 | | | | | | | 53.536 | | | | | | |

Note: Repeated-measured EEG data nested within participant, indicated by participant_id. Sex coded as 0 = female, 1 = male.

**Table S42**

*Hierarchical Linear Models of Absolute Alpha Power Stability, with Session X Age^2^ interactions (Eyes-Closed)*

|  | **Midline** | | | | | | | **Perimeter** | | | | | | |
| --- | --- | --- | --- | --- | --- | --- | --- | --- | --- | --- | --- | --- | --- | --- |
| *Predictors* | *Estimates* | *std. Error* | *std. Beta* | *CI* | *Statistic* | *p* | *p*  *(FDR.adj* | *Estimates* | *std. Error* | *std. Beta* | *CI* | *Statistic* | *p* | *p*  *(FDR.adj* |
| Intercept | 1.408 | 0.063 | 0.093 | 1.283 – 1.533 | 22.194 | **<0.001** |  | 1.074 | 0.056 | 0.095 | 0.963 – 1.185 | 19.021 | **<0.001** |  |
| Session | -0.003 | 0.023 | -0.020 | -0.048 – 0.042 | -0.116 | 0.908 | 0.908 | 0.006 | 0.020 | -0.012 | -0.035 – 0.046 | 0.279 | 0.781 | 0.908 |
| Age^2^ | 0.000 | 0.000 | 0.022 | -0.000 – 0.001 | 1.288 | 0.199 | 0.199 | 0.000 | 0.000 | 0.038 | -0.000 – 0.001 | 1.880 | 0.061 | 0.081 |
| Sex | -0.152 | 0.069 | -0.332 | -0.289 – -0.015 | -2.186 | **0.030** | 0.060 | -0.159 | 0.062 | -0.382 | -0.281 – -0.037 | -2.572 | **0.011** | **0.044** |
| Session X Age² | -0.000 | 0.000 | -0.009 | -0.000 – 0.000 | -1.395 | 0.165 | 0.165 | -0.000 | 0.000 | -0.008 | -0.000 – 0.000 | -1.451 | 0.149 | 0.165 |
| **Random Effects** | | | | | | | | | | | | | | |
| σ^2^ | 0.02 | | | | | | | 0.02 | | | | | | |
| τ_00_ | 0.18 _participant_id_ | | | | | | | 0.14 _participant_id_ | | | | | | |
| ICC | 0.90 | | | | | | | 0.90 | | | | | | |
| N | 175 _participant_id_ | | | | | | | 175 _participant_id_ | | | | | | |
| Observations | 350 | | | | | | | 350 | | | | | | |
| Marginal R^2^ / Conditional R^2^ | 0.028 / 0.899 | | | | | | | 0.042 / 0.900 | | | | | | |
| AIC | 203.812 | | | | | | | 124.138 | | | | | | |

Note: Repeated-measured EEG data nested within participant, indicated by participant_id. Sex coded as 0 = female, 1 = male.

**Table S43**

*Hierarchical Linear Models of Absolute Alpha Power Stability, with Session X Age^2^ interactions (Eyes-Open)*

|  | **Midline** | | | | | | | **Perimeter** | | | | | | |
| --- | --- | --- | --- | --- | --- | --- | --- | --- | --- | --- | --- | --- | --- | --- |
| *Predictors* | *Estimates* | *std. Error* | *std. Beta* | *CI* | *Statistic* | *p* | *p*  *(FDR.adj)* | *Estimates* | *std. Error* | *std. Beta* | *CI* | *Statistic* | *p* | *p*  *(FDR.adj)* |
| Intercept | 0.911 | 0.060 | 0.044 | 0.792 – 1.029 | 15.099 | **<0.001** |  | 0.607 | 0.057 | 0.042 | 0.496 – 0.718 | 10.732 | **<0.001** |  |
| Session | 0.016 | 0.026 | -0.018 | -0.034 – 0.067 | 0.632 | 0.529 | 0.908 | 0.045 | 0.024 | 0.035 | -0.002 – 0.092 | 1.895 | 0.060 | 0.240 |
| Age^2^ | 0.001 | 0.000 | 0.063 | 0.000 – 0.001 | 2.576 | **0.010** | **0.021** | 0.001 | 0.000 | 0.071 | 0.000 – 0.001 | 3.004 | **0.003** | **0.012** |
| Sex | -0.111 | 0.058 | -0.313 | -0.226 – 0.005 | -1.891 | 0.061 | 0.061 | -0.111 | 0.055 | -0.332 | -0.220 – -0.002 | -2.005 | **0.047** | 0.061 |
| Session X Age² | -0.000 | 0.000 | -0.007 | -0.000 – 0.000 | -1.812 | 0.072 | 0.144 | -0.000 | 0.000 | -0.017 | -0.000 – 0.000 | -1.853 | 0.066 | 0.144 |
| **Random Effects** | | | | | | | | | | | | | | |
| σ^2^ | 0.02 | | | | | | | 0.02 | | | | | | |
| τ_00_ | 0.10 _participant_id_ | | | | | | | 0.09 _participant_id_ | | | | | | |
| ICC | 0.83 | | | | | | | 0.83 | | | | | | |
| N | 147 _participant_id_ | | | | | | | 147 _participant_id_ | | | | | | |
| Observations | 294 | | | | | | | 294 | | | | | | |
| Marginal R^2^ / Conditional R^2^ | 0.046 / 0.835 | | | | | | | 0.061 / 0.841 | | | | | | |
| AIC | 106.978 | | | | | | | 68.938 | | | | | | |

Note: Repeated-measured EEG data nested within participant, indicated by participant_id. Sex coded as 0 = female, 1 = male.

**Table S44**

*Hierarchical Linear Models of Absolute Alpha Power Stability, with Session X Sex interactions (Eyes-Closed)*

|  | **Midline** | | | | | | | **Perimeter** | | | | | | |
| --- | --- | --- | --- | --- | --- | --- | --- | --- | --- | --- | --- | --- | --- | --- |
| *Predictors* | *Estimates* | *std. Error* | *std. Beta* | *CI* | *Statistic* | *p* | *p*  *(FDR.adj)* | *Estimates* | *std. Error* | *std. Beta* | *CI* | *Statistic* | *p* | *p*  *(FDR.adj)* |
| Intercept | 1.467 | 0.050 | 0.104 | 1.369 – 1.565 | 29.360 | **<0.001** |  | 1.154 | 0.045 | 0.126 | 1.066 – 1.242 | 25.782 | **<0.001** |  |
| Session | -0.024 | 0.019 | -0.027 | -0.062 – 0.014 | -1.267 | 0.207 | 0.503 | -0.020 | 0.017 | -0.024 | -0.054 – 0.014 | -1.150 | 0.252 | 0.503 |
| Age^2^ | -0.004 | 0.002 | -0.122 | -0.009 – 0.001 | -1.665 | 0.098 | 0.391 | -0.002 | 0.002 | -0.065 | -0.006 – 0.002 | -0.891 | 0.374 | 0.499 |
| Sex | -0.127 | 0.085 | -0.298 | -0.294 – 0.040 | -1.499 | 0.135 | 0.195 | -0.162 | 0.076 | -0.362 | -0.312 – -0.013 | -2.133 | **0.034** | 0.135 |
| Session X Sex | -0.005 | 0.033 | -0.005 | -0.069 – 0.059 | -0.150 | 0.881 | 0.907 | 0.011 | 0.029 | 0.013 | -0.047 – 0.068 | 0.362 | 0.718 | 0.907 |
| **Random Effects** | | | | | | | | | | | | | | |
| σ^2^ | 0.02 | | | | | | | 0.02 | | | | | | |
| τ_00_ | 0.18 _participant_id_ | | | | | | | 0.14 _participant_id_ | | | | | | |
| ICC | 0.89 | | | | | | | 0.89 | | | | | | |
| N | 175 _participant_id_ | | | | | | | 175 _participant_id_ | | | | | | |
| Observations | 350 | | | | | | | 350 | | | | | | |
| Marginal R^2^ / Conditional R^2^ | 0.040 / 0.898 | | | | | | | 0.037 / 0.899 | | | | | | |
| AIC | 186.238 | | | | | | | 109.841 | | | | | | |

Note: Repeated-measured EEG data nested within participant, indicated by participant_id. Sex coded as 0 = female, 1 = male.

**Table S45**

*Hierarchical Linear Models of Absolute Alpha Power Stability, with Session X Sex interactions (Eyes-Closed)*

|  | **Midline** | | | | | | | **Perimeter** | | | | | | |
| --- | --- | --- | --- | --- | --- | --- | --- | --- | --- | --- | --- | --- | --- | --- |
| *Predictors* | *Estimates* | *std. Error* | *std. Beta* | *CI* | *Statistic* | *p* | *p*  *(FDR.adj)* | *Estimates* | *std. Error* | *std. Beta* | *CI* | *Statistic* | *p* | *p*  *(FDR.adj)* |
| Intercept | 1.014 | 0.047 | 0.105 | 0.922 – 1.107 | 21.587 | **<0.001** |  | 0.725 | 0.044 | 0.117 | 0.638 – 0.812 | 16.361 | **<0.001** |  |
| Session | -0.012 | 0.021 | -0.016 | -0.054 – 0.030 | -0.551 | 0.582 | 0.582 | 0.014 | 0.020 | 0.020 | -0.025 – 0.053 | 0.688 | 0.492 | 0.582 |
| Age^2^ | -0.002 | 0.002 | -0.076 | -0.006 – 0.002 | -0.963 | 0.337 | 0.499 | 0.000 | 0.002 | 0.004 | -0.004 – 0.004 | 0.052 | 0.959 | 0.959 |
| Sex | -0.081 | 0.080 | -0.309 | -0.239 – 0.077 | -1.007 | 0.315 | 0.315 | -0.111 | 0.076 | -0.344 | -0.260 – 0.039 | -1.457 | 0.146 | 0.195 |
| Session X Sex | -0.019 | 0.036 | -0.027 | -0.091 – 0.052 | -0.535 | 0.594 | 0.907 | -0.004 | 0.034 | -0.006 | -0.071 – 0.063 | -0.117 | 0.907 | 0.907 |
| **Random Effects** | | | | | | | | | | | | | | |
| σ^2^ | 0.02 | | | | | | | 0.02 | | | | | | |
| τ_00_ | 0.10 _participant_id_ | | | | | | | 0.09 _participant_id_ | | | | | | |
| ICC | 0.83 | | | | | | | 0.83 | | | | | | |
| N | 147 _participant_id_ | | | | | | | 147 _participant_id_ | | | | | | |
| Observations | 294 | | | | | | | 294 | | | | | | |
| Marginal R^2^ / Conditional R^2^ | 0.030 / 0.831 | | | | | | | 0.026 / 0.838 | | | | | | |
| AIC | 95.361 | | | | | | | 60.877 | | | | | | |

Note: Repeated-measured EEG data nested within participant, indicated by participant_id. Sex coded as 0 = female, 1 = male.
